# Chromosome-level genome assembly of autotetraploid *Selenicereus megalanthus* and gaining genomic insights into the evolution of trait patterning in diploid and polyploid pitaya species

**DOI:** 10.1101/2024.06.23.600268

**Authors:** Qamar U Zaman, Liu Hui, Mian Faisal Nazir, Guoqing Wang, Vanika Garg, Muhammad Ikram, Ali Raza, Wei Lv, Darya Khan, Aamir Ali Khokhar, Zhang You, Annapurna Chitikineni, Babar Usman, Cui Jianpeng, Xulong Yang, Shiyou Zuo, Peifeng Liu, Sunjeet Kumar, Mengqi Guo, Zhi-Xin Zhu, Girish Dwivedi, Yong-Hua Qin, Rajeev K. Varshney, Hua-Feng Wang

## Abstract

Yellow pitaya (*Selenicereus megalanthus*, 2n=4x=44) breeding remains severely hindered due to lacking a reference genome. Here we report yellow pitaya’s high-quality chromosome-level genome assembly and link the phenotypic trait with genomic data, based on Hi-C, ATAC, and RNA-seq data of specific tissues. We declared yellow pitaya as an autotetraploid with a 7.16 Gb genome size (harboring 27,246 high confidence genes) and majorly evolved from diploid ancestors, which remains unknown. Beyond generating the genome assembly, we explored the 3D chromatin organization which revealed insights into the genome, compartments A (648 and 519), compartments B (728 and 1064), topologically associated domains-TADs (3376 and 2031), and varying numbers of structural variations (SVs) in diploid and polyploid pitaya species, respectively. Overall, TAD boundaries were enriched with the motifs of *AP2*, *WRKY18/60/75*, *MYB63/116*, *PHL2*, and *GATA8* in both pitaya species. By linking the open chromatin genomic structure to function, we identified the major changes in betalains biosynthesis pathway in diploid and polyploid pitaya. Moreover, the higher genetic expression of *SmeADH1* [Chr11, Compartment A (135400000 - 135500000), genes inside the TAD region (135480000 - 135520000)], and lower expression of *HuDOPA* [Chr11, Compartment A (87100000 - 87200000), genes inside the TAD region (87160000-87200000)] acts as a key regulator of yellow and red color on the pericarp of polyploid and diploid pitaya, respectively. In addition, higher expression of *HuCYP76AD1* genes in diploid pitaya and lower expression of *SmeCYP76AD1* in polyploid pitaya potentially created the difference in the oxidase process that led to the production of betacyanin and betaxanthin, respectively. Furthermore, our results revealed not only the type of motifs that play a potential role in trait patterning but we also further uncovered that motif count in TAD-boundaries may impact the gene expression within the TAD regions of diploid and polyploid pitaya. Our valuable genomic resource and comparison of 3D euchromatin architecture of diploid and polyploid pitaya species will not only aid in the advancement of molecular breeding efforts but also offer insights into the organization of genomes, SVs, compartmentalization (A and B), and TADs, which have the potential to strengthen the idea of TADs-based trait improvement to achieve global food security.

## Main

Dragon fruit or pitaya belongs to the family Cactaceae (order Caryophyllales), an ancient family, that originated ∼65 million years ago^1^. The evidence of chloroplast DNA and internal transcribed sequence (ITS) divergence data suggested that cacti originated 30 million years ago in mid-tertiary^1^. The genus *Selenicereus* (formerly *Hylocereus*), with unknown origin, probably native to Central America (Southern Mexico, Salvador, Guatemala, and Costa Rica), evolved around the Early-Cretaceous (when Gondwana continent split into South America and Africa)^2^. The genus *Selenicereus* is divided into 28 species based on morphological characteristics^3^ and widely distributed from North to South America^4^. Among these species, *S. megalanthus* (2n=4x=44) and *S. undatus* (2n=22) (formerly known as *Hylocereus undatus*) are the most commercialized and consumed groups of plants comprising tremendous value with an attractive appearance of fruit, color, flavor, rich sources of vitamins, antioxidants^5^, minerals, phytochemicals (betacyanins, phenolics, polysaccharides, terpenoids, alkaloids)^6^, dietary fiber and prevent cancer^7–9^. Fruit morphology, such as size, color, number of spines, and length of areoles in cladodes are the main taxonomic evidence to differentiate between these species^7^.

Pitaya fruit was first cultivated probably by the “Aztec-civilization” back in the 13^th^ century who supported the richness of crop biodiversity in their ruled region^10^. Today, it is cultivated throughout the world’s tropical and subtropical regions, particularly Southern Mexico and South-East Asia^3^. Due to its spectacular appearance and antioxidant capacity, pitaya demand is growing and becoming popular with a substantial increase in its planting area and production. Vietnam is the biggest producer^11^ and shares 50% of the total global market (14.11 billion USD) (https://ap.fftc.org.tw/article/1596), followed by China, Mexico, Colombia, Thailand, Malaysia, India, Indonesia, Australia, Israel, the U.S.A, Pakistan, and many other countries growing on a small scale^4,7^. In addition, pitaya fruit contains potential nutrients, bioactive biochemicals, and a high level of water-soluble betalains, a good source for the food industry, and tyrosine-originated alkaloid pigment^12^. Despite their significance in agricultural sectors, pitaya fruits are well-known for being highly drought-tolerant^3^. When the earth underwent a drop in CO_2_ concentration and dryland in the Gondwana continent, Cactaceae arose and acclimatized to aridity^13^. Pitaya fruit as a climbing cactus that has developed a fascinating succulent stem, aerial roots to absorb humidity, and watery fruits.

Based on hallmark flavor, morphological and genetic fruit characteristics, diploid pitaya (*S. undatus*-red peel white flesh) and polyploid pitaya (*S.* megalanthus-yellow-peel white flesh) species taxonomically carried distinct lineage. *S. megalanthus* has yellow peel with large thorns, four or more ribbed cladode, small fruit scales, weak vegetative production, fruit small to medium in size, large seed size, white flesh inside the fruit, sweeter, parenchyma and chlorenchyma mixed as one layer (**Fig. 1a**). While *S. undatus* has three ribbed cladodes, a red-bright spineless peel with broad overlapping green fins, strong vegetative production, large fruit size, white flesh, less sweet, small seed size and comprised of separate parenchyma and chlorenchyma layers (**Fig. 1b**). Problematic classification and ploidy-level description of yellow pitaya from the last 100 years of history, raised the fundamental questions. In 1920, yellow pitaya was classified as *Mediocactus megalanthus* based on the “*Medio*” phenotypic characteristics of the *Hylocereus* and *Selenicereus* genus^4,14^. In 1953, yellow pitaya was classified into the *Selenicereus* genus^15^, but in 2003, Baur and colleagues placed it, in the *Hylocereus* genus and called it allotetraploid origin followed by chromosome doubling of closely related species of *Hylocereus* and *Selenicereus*^16^. In 2013, Plume and colleagues suggested that yellow pitaya is taxonomically autopolyploid but narrowly allotetraploid^17^. In 2017, Korotkova and colleagues abolished the *Hylocereus* genus and transferred all species into the *Selenicereus* genus^18^ (**Fig. 1c**). Despite their common and distinguished features, how these diploid and tetraploid species are genetically distinct and evolved, remains a mystery in plant biology and horticultural crops.

**Fig 1:**
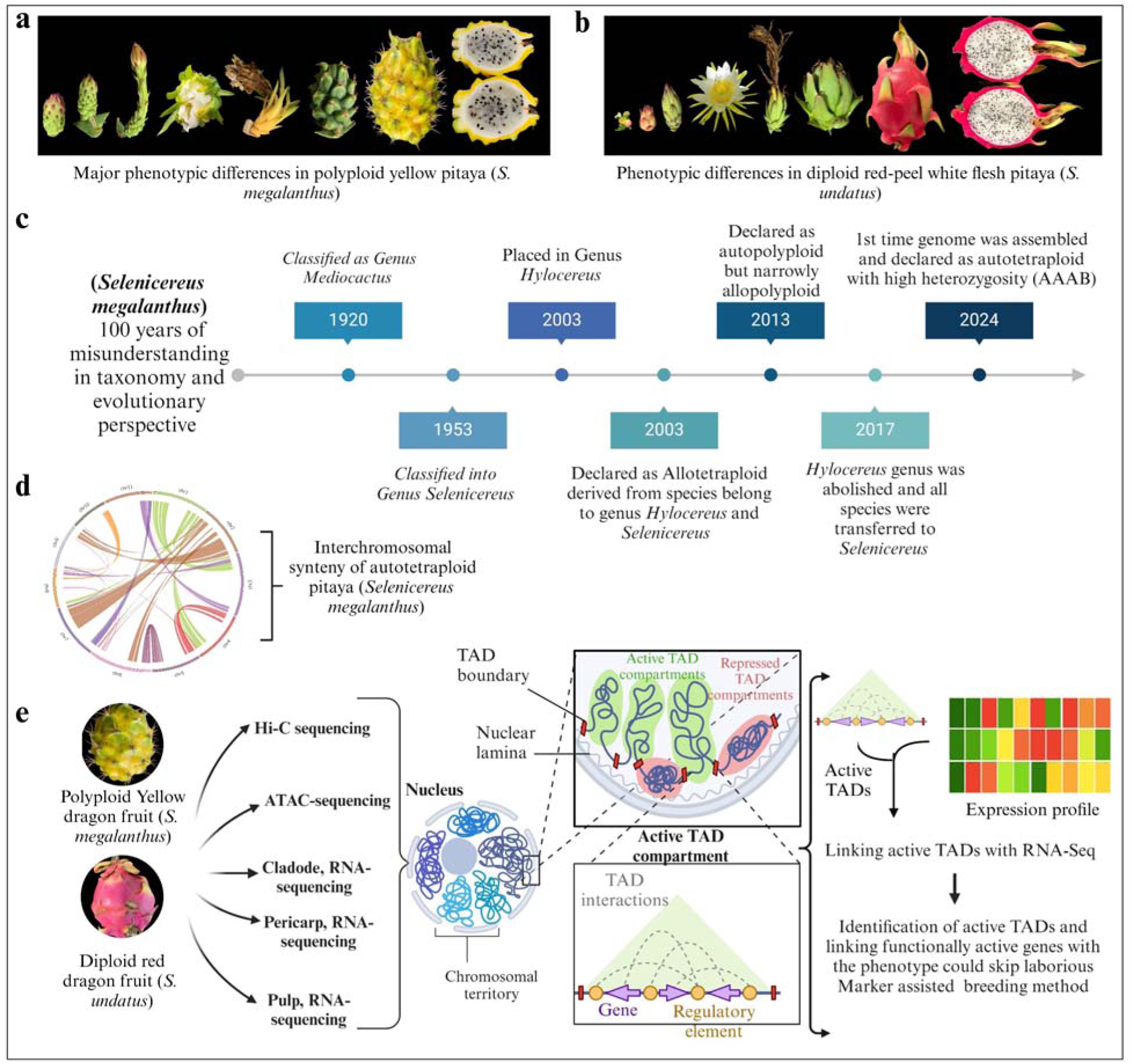
Misunderstood taxonomy, evolution of polyploid *S. megalanthus*. Comparative genomics and phenomics of diploid and polyploid pitaya. **a)** Phenotypic characteristics of yellow and **b)** red pitaya (white flesh). **c)** Schematic presentation of 100 years of misunderstood taxonomy of yellow pitaya. Our assembled genomic resource provides insights into the yellow pitaya evolution (97% autotetraploid, 3% allopolyploid). **d)** It displays the interchromosomal synteny of a newly assembled genome of yellow pitaya. **e)** Comparative genomics of diploid and tetraploid pitaya. Combined analysis of the 3D genome, TAD boundaries, and RNA-seq data revealed specific genes correlated with the respective phenotypic traits of both species (fruit color). This method can skip the laborious MAS breeding method.

Despite innovations in modern genomics tools, a high-quality reference genome of yellow pitaya was still lacking, hindering molecular breeding efforts. Mapping phenotypic variation between diploid red pitaya (red peel, thin stem/cladode, green fins) and tetraploid yellow pitaya (yellow peel, thick stem/cladode, spines) is urgently needed to accelerate the breeding program. How do DNA replication and transcription organize their role in tightly linked diploid and polyploid pitaya genomes? How are chromosome-chromosome folding, chromatin structure, topologically associated domains (TADs) borders, active (A) and inactive compartments (B) linked to the DNA-dependent processes? In general, the interaction frequency within the TAD region is significantly higher than the two adjacent TAD regions, and the TAD boundary is rich in promoter-related transcription factors (TFs), transcription start sites, housekeeping genes, tRNA genes, and short interspersed nuclear elements (SINEs), which play an important role in maintaining the stability of TAD structure^19^. To address these questions, comparative genomic studies have been conducted to understand the formation of ploidy level, evolutionary resemblance within the *Selenicereus* genus, pitaya fruit development, and adaptation to tropical and subtropical regions.

Here, we presented the high-quality chromosome-level genome sequencing and assembly of autotetraploid yellow pitaya (*S. megalanthus* L). (**Fig. 1d**) using long-read PacBio HiFi long-read sequencing. Furthermore, diploid red pitaya (2n=22) and autotetraploid yellow pitaya (2n=4x=44) were employed for Hi-C (high-throughput chromosome conformation capture) sequencing, ATAC-sequencing, and tissues specific transcriptome sequencing (**Fig. 1e**). By comparing both genomes, we identified chromosomal organization, chromatin structure, genome compartmentalization, TAD boundaries, and structural variations (SVs) throughout the diploid (*S. undatus*) and tetraploid (*S. megalanthus*) genomes. Our results declare the yellow pitaya as an autotetraploid with high heterozygosity (AAAB) and both species comparison not only provides fundamental insights into the complexities of diploid and polyploid genomes but also offers an understanding for the fast-tracking the breeding of these vital horticultural crops globally.

## Results

### Chromosome-level genome sequencing, assembly, genome characteristics, and evolution of polyploid pitaya (*S. megalanthus*)

We assembled a chromosome-level genome assembly of yellow pitaya [(*S. megalanthus*-K. Schum. Ex Vaupel-Ralf Bauer), previously known as *Hylocereus megalanthus*], using the *de-novo* genome sequencing. For a preliminary overview, DNB sequencing and *K*-mer (k-base fragment) frequency analysis were performed, and initially, we estimated the overall ∼5.64 Gb genome size of yellow pitaya. However, we also observed that *S. megalanthus* had a significant proportion of AAAB type *K*-mer pairs (64%), AABB type (21%), AB type (0.03%), AAB type, and other types (0.02%) (**Supplementary Fig. 1**). The estimated results of 21 *K*-mer with the highest fitting degree were selected as a final spectrum to predict genome characteristics. The total sequencing size was ∼201.97 Gb and we obtained 194.17 Gb of clean data. In a customized analysis, genome ploidy was evaluated using Smudgeplot software^20^ following the previous report^21^ (**Supplementary Fig. 2)**. The genome ploidy level was found to be tetraploid with a karyotype 2n=4x=44. For *de-novo* assembly, genome compartmentalization, and identification of TADs to link the RNA-seq data, we incorporated multiple sequencing technologies including PacBio HiFi, Hi-C, ATAC, and RNA-seq. PacBio_Revio sequencing platform was used by circular consensus sequencing (CCS) method, and CCS reads (97.76 Gb) were generated with an average length of 15.95 kb at N50 of 16 kb (**Supplementary Table 1**). The preliminary assembly comprised 900 contigs with a contig N50 of 13.18 Mb (**Supplementary Fig. 3**). The genome was assembled after processing PacBio HiFi and Hi-C data with Purge-Haplotigs^22^ by identifying and removing redundant heterozygous sequences, based on read depth distribution (**Supplementary Fig. 4**) and sequence similarity. The assembly was elevated into the super-scaffolds using Hi-C interaction datasets (477.11 GB, 100 X genome coverage), and 414 assembled contigs were anchored into the eleven chromosomes with a rate of 97.73% (**Fig. 2**, **Table 1**). However, the total assembled genome size was much bigger than the estimated k-mer frequency around ∼7.16 Gb (2n=4X=44) with a 37.01% GC content.

**Fig 2:**
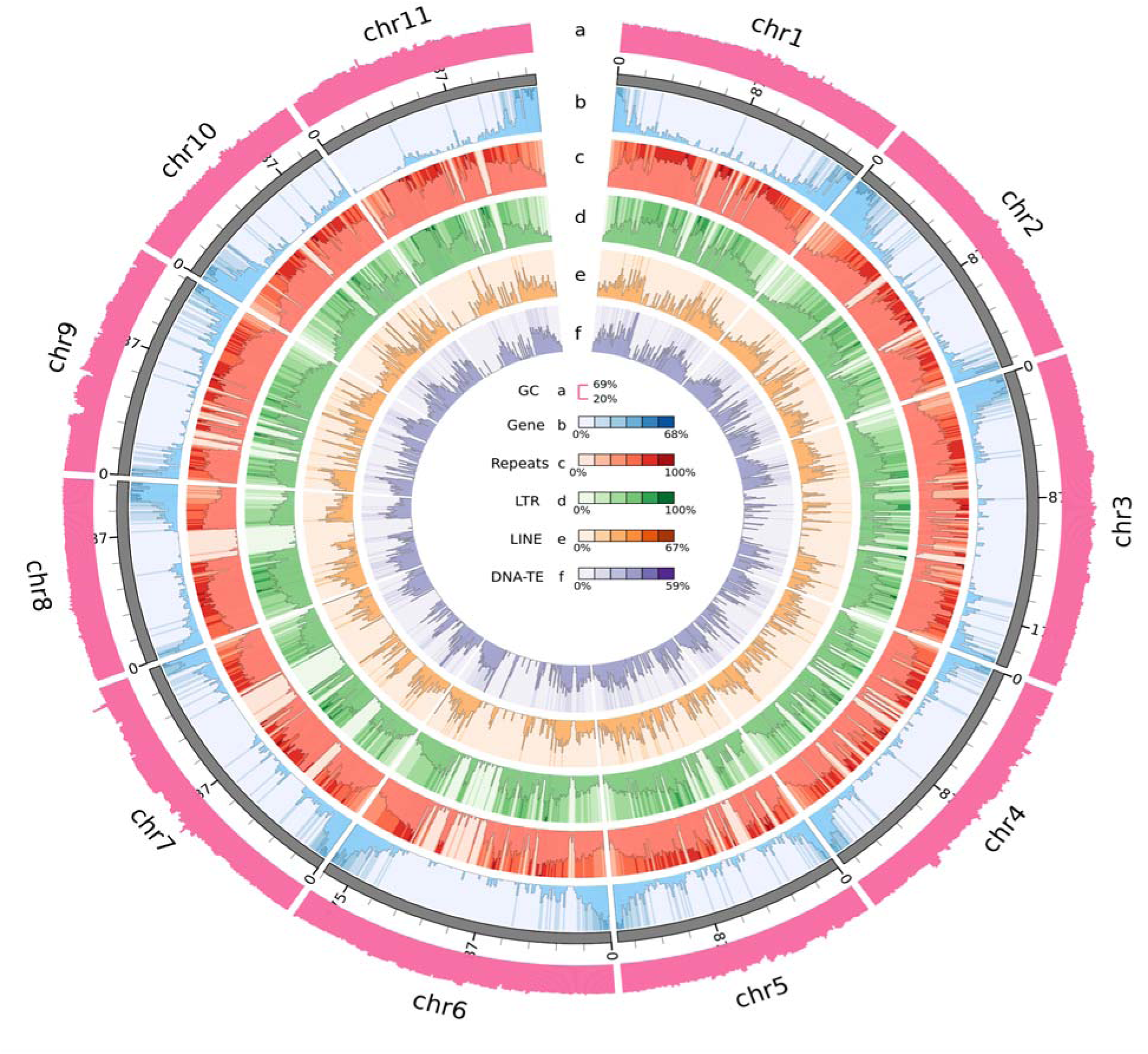
The genomic features of autotetraploid yellow pitaya genome (*S. megalanthus*). The rings indicate the 11 chromosomes and **a)** represent GC content distribution, **b)** Gene density distribution, **c)** Total repeat density distribution, **d)** long terminal repeats (LTR) distribution, **e)** LINE density distribution, and **f)** DNA-transposable elements distribution.

**Table 1:** Summary of yellow pitaya (*Selenicereus megalanthus*) genome assembly.

| Features | *Selenicereus megalanthus* |
| --- | --- |
| Total length of contigs (Gb) | 1.7916 |
| Number of assembled sequences by HiFi | 2614 |
| Number of assembled sequences by HiFi+Hi-C+Purge | 425 |
| Contig N50 (Kbps) | 7056.17 |
| GC % | 37.01 |
| Monoploid genome assembly size (Gb) | 1.79 |
| Genes with defined alleles | 27246 |
| Functionally annotated genes (91.48%) | 24924 |
| Anchor rate (%) | 97.73 |

The genome was compared to the NT database and the genome assembly was evaluated by the “compleasm-method”^23^. Multiple genome assessments supported the high-quality autotetraploid yellow pitaya genome assembly. Ortholog database (OrthoDB) was used to assess the integrity of the genome assembly. Hundreds of genomes were sampled, from which single-copy orthologou >90% genes were selected to construct a gene set with multiple phylogenetic branches.

Benchmarking Universal Single-Copy Orthologue (BUSCO, 100% of the 1614 core eukaryotic genes) analysis indicated the completeness of 94.42% with 3.72% of duplication in the assembled genome (**Supplementary Table 2**). The length of 11 chromosomes ranged from 115.62-204.98 Mb, and unanchored genomic sequences of 40.69 Mb. The assembly had contig N50 13Mb, and we applied Hi-C data to order and orient the resulting contigs. Chromosome-level genome assembly was developed, among which 97.73% was anchored on 11 chromosomes, and an overview is presented in **Fig. 3A** (**Supplementary Table 2**). Unanchored sequences could be a part of the genome that were inserted as per our assumption (**Supplementary Fig. 5**) or lateral gene transfer (LGT) during the domestication process. We have identified the AAAB pattern as a dominant component accounting for 64% of the examined K-mers (**Supplementary Fig. 2**). These results collectively point out a high level of heterozygosity in *S. megalanthus*, which might be a reason for clonal propagation^24^, or somatic mutations^25^ that may be included in a heterozygous manner within a clone, or narrow allopolyploidy.

**Fig 3:**
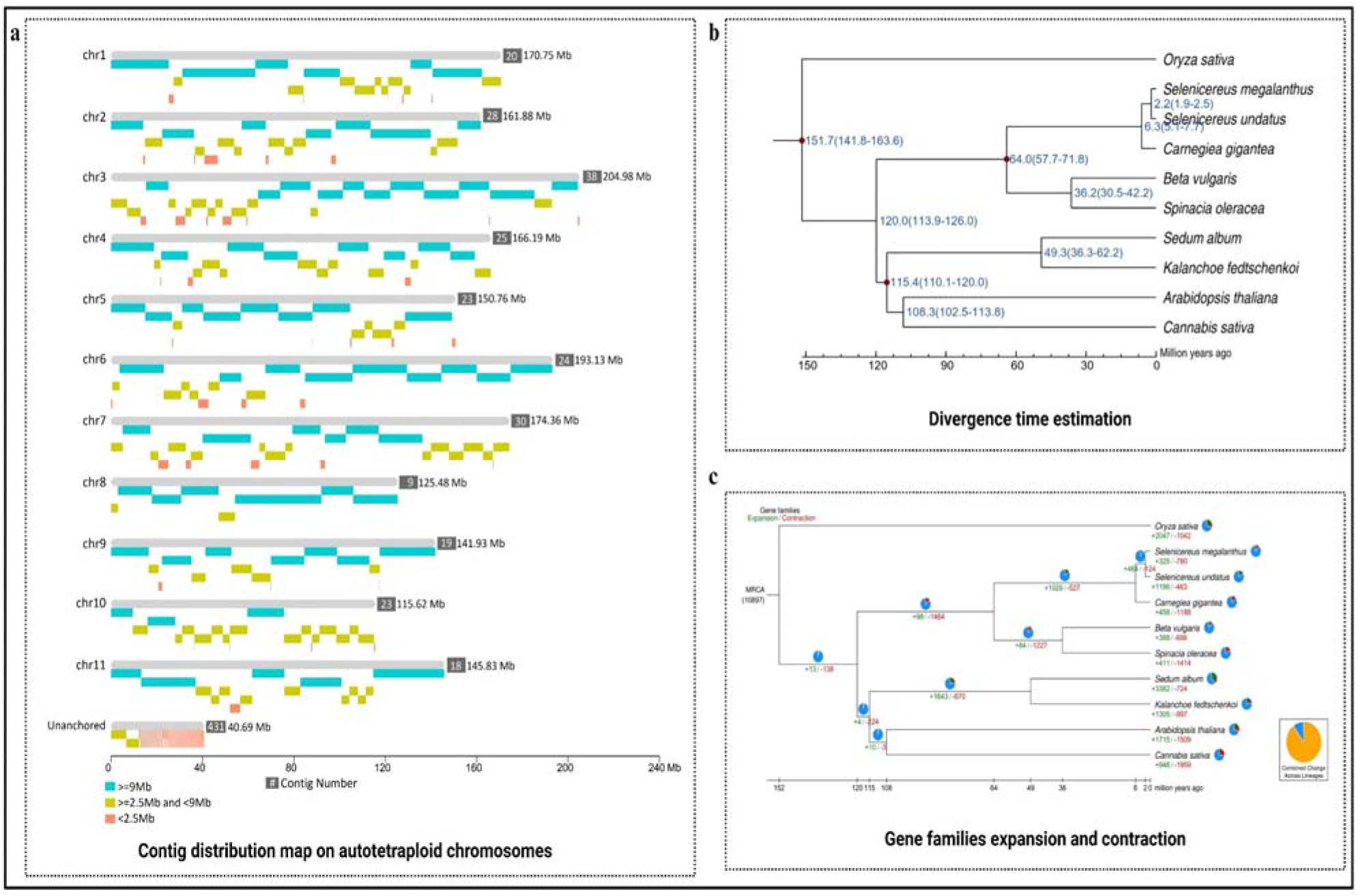
Contig distribution map on autotetraploid chromosomes, the divergence time of *S. megalanthus*, estimation of expansion and contraction of gene families. **a)** The grey color indicates the length of each chromosome and other colors represent contig with different length ranges. **b)** Divergence time of *Selenicereus megalanthus* and other plant species. The number at the node position represents the divergence time of plant species from their ancestors million years ago (MYA). Divergence time supported by the 95% highest posterior density (HPD) is also mentioned in brackets. The branch length of a phylogenetic tree represents only the number of base substitutions or genetic distance per point. **c)** Expansion and contraction of gene families in plant species. The green color indicates the gene families that were added to the genome during the evolutionary process of species, and the red color represents the loss of gene families from the plant species.

We used Hi-C data (477.11 Gb) to group the scaffold contigs and confirm the orientation of all chromosomes using their physical proximity in the yellow pitaya genome. With an interaction signal, 414 scaffolds were generated with a total length of 1.79 Gb anchored on 11 chromosomes (**Table 1**). The number of interactions between the contigs was counted and the contigs were clustered according to their number of interactions and divided into the specified class number and 11 chromosomes in the yellow pitaya. Based on the interaction frequencies between the genomic regions, they were used to group contig sequences and chromosome-level genome assembly. Subsequently, we identified 27,246 high-confidence genes in *S. megalanthus* based on *de-novo* prediction, homolog comparison, comparing homologous in biological species (*S. undatus* and *Carnegiea gigantea*), predicting transcription factors based on coding genes, BUSCO library alignment and high-quality set of genes were integrated using the BGI self-developed HiFAP software (**Supplementary Table 3**). Annotation of protein-coding genes and BUSCO assessment exhibited a high degree of completeness and accuracy (**Supplementary Table 4, Supplementary Fig. 6**). Among the final set of 27,246 genes (**Supplementary Table 3**) with 36,277 transcripts (**Supplementary Table 5**), including 24,924 (91.48%) with 33713 transcripts were functionally annotated (**Supplementary Table 6**), with an average gene length of 6255.31 bp (mean length of transcripts-7247.29 bp), average CDS length 1187.89 bp, average exon per gene 5.42, average exon length 298.64 (mean length of exon-225.43) and average intron length 1049.92 (**Supplementary Fig. 7**).

The genome sequences of ten plant species were collected and subjected to comparative genomics analysis to illustrate the evolution and divergence of *S. megalanthus*. Our results revealed 146,064 gene families, 6,751 common gene families, and 1,149 single-copy orthologs genes in ten species (**Supplementary Table 7**). These 1149 single-copy orthologs were subjected to construct the phylogenetic tree (**Fig. 3b**). Of the total 27,246 genes, *S. megalanthus* contained 17885 gene families in 25268 genes including 81 unique families. We also discovered a resolution of the phylogenetic relationship among the genomes of ten plant species. The sister species of pitaya (*S. undatus-*diploid and *S. megalanthus*-polyploid) exhibited a divergence time ∼1.9-2.2 million years ago (MYA) (**Fig. 3b**). However, the divergence time was 5.1-7.7 MYA between pitaya sister species and closely related cactus (*Carnegiea gigantea*). *S. megalanthus* exhibited numerous expanded (325) and contracted (780) gene families among these selected plant species. In addition, we found that 464 gene families were commonly expanded and 124 contracted between diploid and polyploid pitaya species (**Fig. 3c**). Importantly, these expanded gene families were enriched in gene ontology (GO) terms like “protein dimerization activity”, “terpene synthase activity”, “diterpenoid biosynthetic pathway” and “glycosyltransferase activity” in *S. megalanthus* suggesting their role in the biosynthesis of “betaxanthin” to produce the yellow pitaya fruit (**Supplementary Fig 8**). We used the WGDI software to calculate the synonymous substitution rate (*Ks*), the peaks corresponding to the existence of genome-doubling, and computed the collinearity blocks which can reduce the error caused by the tandem repeats. The distribution of *Ks* of each homologous gene pair within *S. megalanthus* exhibited a single peak, revealing a single whole genome duplication event in yellow pitaya. The most recent event of WGD occurred dating back to 1.9-2.2 MYA, contributing to the tetraploidy in *S. megalanthus* (**Supplementary Fig. 8)**. Collinearity between *S. undatus* and *S. megalanthus* has illuminated the genomic evolution and relatedness of two pitaya species (**Supplementary Fig. 9).** Hence we discovered 777 syntenic blocks (average 37 syntenic genes per block) on both pitaya species, suggesting conserved gene order across the genomes (**Supplementary Table 8).** The adaptive evolution and divergence of species are ultimately attributed to the evolution of genes. Synonymous mutations (*Ks*) represent the changes in the gene sequence but not in the amino acid sequence. Nonsynonym mutations (*Ka*) represent the genetic sequence changes that alter the amino acids as well. The nucleotide substitution rate per site per year was calculated using the formula “r=K/2T where T is the divergence time (2.2 MYA) and K was calculated as 0.01008145 (K= −0.75ln (1-4λ/3). The insertion time of the LTR was calculated with the formula *T=K/2r* (or r=K/2T) in which r represents the substitution rate per site per year. The nucleotide substitution rate per site per year “r” was computed as 2.29 X 10^-9^.

### The 3-dimensional genome structure of diploid and polyploid pitaya species

The eukaryotic chromatin genome is hierarchically packed in chromatin loops, A and B compartments, and TADs. We generated Hi-C data, to investigate the diploid and polyploid pitaya 3D genome architecture. Before aligning the clean data to their corresponding reference genome of diploid^26^ and polyploid pitaya (newly assembled genome), reads containing adapter sequences were filtered and subjected to verify the distribution of sequence base content (**Supplementary Fig. 10 & Fig. 11**), base mass distribution (**Supplementary Fig. 12**), and GC content (**Supplementary Fig. 13**) in all biological repeats. In the total raw data, 958 to 997 million clean reads (filter reads + mapped reads) were produced in diploid pitaya. In comparison, 1,582 to 1,916 million clean reads were generated in polyploid pitaya species (**Supplementary Table 9**). To ensure the quality, accuracy, and reproducibility of Hi-C sequencing libraries, we analyzed the correlation between the parallel samples of diploid (0.992) and polyploid (0.989) pitaya species using GenomeDISCO software^27^. Hi-C analysis was employed to construct a 3D conformation of the genome by aligning pairwise reads at different loci on the reference genome of both pitaya species. To construct the interaction matrix, the genome was divided into multiple bins of equal size, and then filtered valid pairs (Q30) (**Supplementary Table 10**) were assigned to these bins. Due to the uneven distribution of GC content and enzyme cleavage sites on the reference genomes, the original interaction matrix was corrected using the Juicer^28^, and a 100-kb resolution genome-wide interaction matrix (**Supplementary Fig. 14**) and a single chromosome interaction matrix with a resolution of 40 kb were generated for diploid (*S. undatus*) (**Supplementary Fig. 15**) and polyploid (*S. megalanthus*) (**Supplementary Fig. 16**) pitaya species. To validate the results credibility, a resolution chart was generated which showed that 80% of the bins could reach 1000 contacts (**Supplementary Fig. 17A & 17B**), and the amount of the data generated by Hi-C can support the subsequent analysis as suggested by Rao and colleagues (2014)^29^. In addition, a genome-wide interaction attenuation curve was generated to investigate the relationship between the distance and interaction intensity of *cis*-interaction sites (**Supplementary Fig. 18A & 18B**).

According to the single-chromosome interaction matrix, the Hi-C based 3D genome structure of diploid and polyploid pitaya was partitioned into their active (A) and inactive (B) compartments. The PC1 value of the bin was acquired by the principal component analysis (PCA)^30^ and the C-score method of bin^31^. The results of both pitaya species illustrated that the B compartment was more compact than the A compartment. After analyzing the genomic compartments using the C-score method, the positive and negative bins were identified as compartment A and compartment B to obtain the distribution of the compartments on each chromosome. A single chromosome interaction matrix with a resolution of 100 kb was constructed for diploid and polyploid pitaya, guiding that compartment A was comprised of chromatin active region corresponding to the gene density, active histones, and open chromatin. While compartment B corresponds to the chromatin inactive region of the genome which exhibited low gene density, few active histones, and compact chromatin (**Fig 4a & Fig 4b**). The number and length distribution of the genomic compartments were computed by combining the bins of the consecutive compartment A (4,697 in *S. undatus* and 6,114 in *S. megalanthus*) and compartment B (8,576 in *S. undatus* and 8,834 in *S. megalanthus*) in both pitaya species (**Table 2**). In total, we found 648 compartment A, 728 compartment B in diploid pitaya (*S. undatus*), and 519 compartment A, 1,064 compartment B in polyploid pitaya (*S. megalanthus*). In diploid pitaya, compartment A comprised of 19,811 genes out of 4,697 bins and compartment B contained 7,696 genes out of 8,576 bins. However, polyploid pitaya exhibited that compartment A carried 19,956 genes out of 6,114 bins and compartment B kept 6,829 genes in 8,834 bins (**Supplementary Table 11)**.

**Fig 4:**
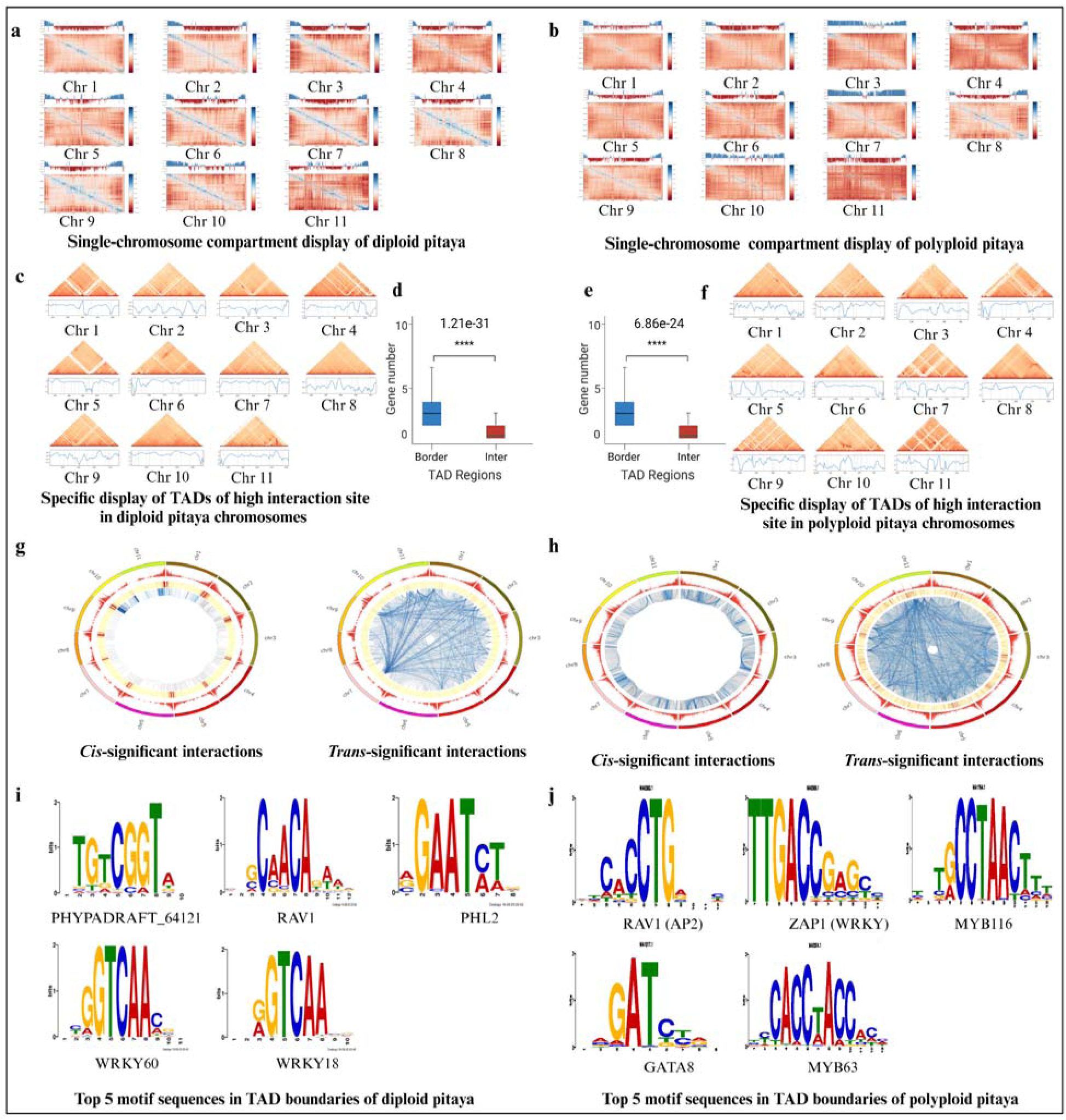
3-D genome structural analysis of diploid and polyploid pitaya. **a)** Single chromosome compartments of *S. undatus.* **b)** Single chromosome compartments of *S. megalanthus.* [The upper part of Figures 4a & 4b denotes the distribution of the values of compartments A/B. Blue bars represent the active (A) part of the genome while the red color bars exhibit the inactive (B) part of the genome. The lower part of the figure denotes the correlation matrix converted from a single chromosome interaction matrix. The color bar represents the correlation coefficient]. **c)** Single chromosome TADs of *S. undatus*. The horizontal axis denotes the position on th reference genome (Unit: Mb). The upper part of the figure represents the interaction at th specific position of single chromosomes. In the lower part of the figure, the blue line shows the insulation score, and the gray line denotes the TAD boundary. **d)** Number of genes within the TAD boundaries of diploid pitaya. **e)** Number of genes within the TAD boundaries of polyploid pitaya. [The horizontal axis in figures 4d & 4e denotes the number of genes within TAD (Inter) and at the TAD boundary (Border). The significance test results in **** represent 0.0001 ≥ p]**. f)** Number of genes within the TAD boundaries of polyploid pitaya. The horizontal axis denotes the number of genes within TAD (Inter) and at the TAD boundary (Border). The significance test results in **** represent 0.0001 ≥ p**. g)** Single chromosome TADs of *S. megalanthus*. The horizontal axis denotes the position on the reference genome (Unit: Mb). The upper part of the figure represents the interaction at the specific position of single chromosomes. In the lower part of the figure, the blue line shows the insulation score, and the gray line denotes the TAD boundary. **h)** Circle plot of genome-wide *Cis/trans*-significant interaction sites in diploid pitaya. **i)** Circle plot of genome-wide *Cis/trans*-significant interaction sites in polyploid pitaya. In Figures 4h & 4i, the name and number of chromosomes are presented clockwise in the figure. The red color exhibits the number of genes on each chromosome. From dark blue to light blue lines indicate the large to small p-value and cis-significant interaction locus on each chromosome. Red bars indicate the significant trans-interactions with other chromosomes. The blue color, from dark to light exhibits the number of reads supporting the interactions from large to small. **j)** Top 5 motif sequences in *S. undatus*. **k)** Top 5 motif sequences in *S. megalanthus*. In Figures 4j & 4k, the horizontal axis exhibits the length of the motif (Unit: bp), while the longitudinal axis represents the frequency distribution of ATGC bases.

**Table 2:**
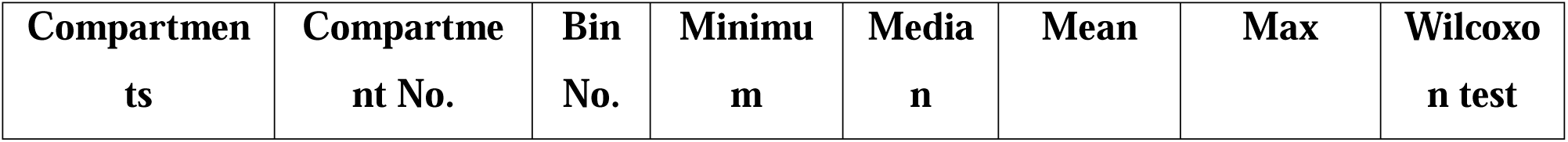
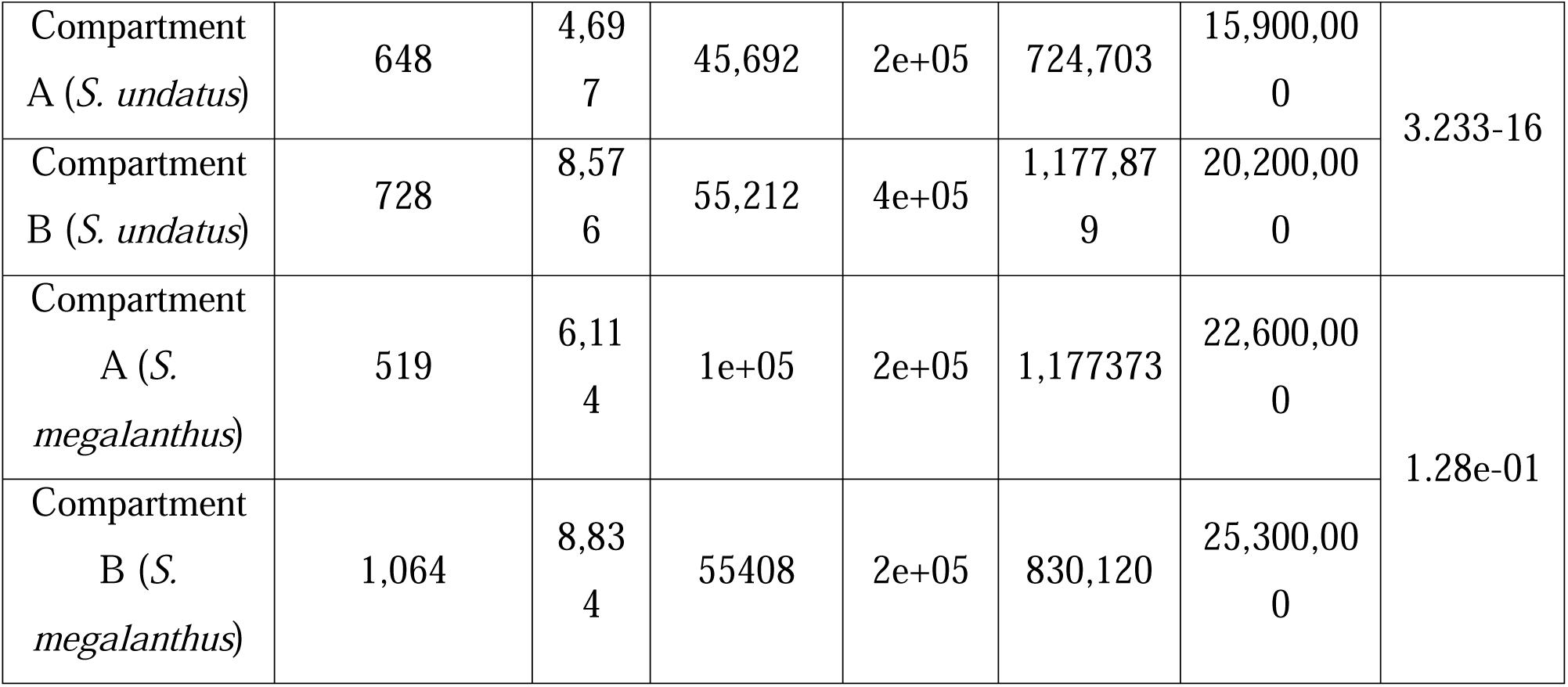
Statistical table of the number and length distribution of compartments.

The length distribution of compartment A and compartment B was compared and displayed in the box plot, which showed highly significant differences (0.0001 ≥ p) determined by Wilcoxon’s rank-sum test (**Supplementary Fig. 19A & 19B**). In addition, Rao and colleagues (2014)^29^ reported that GC content in the genome sequence might be associated with the structure of compartment A and compartment B. However, in our case of A and B compartments of both species, we could not find any significant differences in GC content distribution (**Supplementary Fig. 20A & 20B**). Further analysis illustrated the distinct topologically associated domains (TADs) organization on a chromosome which showed significantly higher interaction frequency within the TAD region than in two adjacent TAD regions. Using the insulation score method^32^, TAD boundaries were computed in both pitaya species (**Fig. 4c & 4f)**. The insulation score reflected the interaction between two sides of each bin on pitaya reference genome. TAD boundary was marked, when the insulation score was lower on both sides of the bin, which was considered as interaction isolation of the bin from other TADs. The single chromosome interaction matrix of TAD was constructed at 40-kb resolution for both species (**Supplementary Fig. 21 & Fig. 22**) including their specific TADs region of higher interaction (**Fig 4c. & Fig. 4f**). In total, 3376 TADs were identified in diploid pitaya genome and 2031 TAD were predicted in polyploid pitaya genome (**Supplementary Table 12**) with the mean length of TAD 401056 bp/pc and 861897 bp/pc, respectively.

To identify the number of genes within the TADs and TAD boundaries, we divided all bins in the pitaya species genome into two categories, including the genes located inside the TAD (Inter), the genes located at the TAD boundary (Border), and the significance of gene number distribution was computed using Wilcoxon’s rank sum test (**Fig. 4d & 4e)**. The data discovered that 24,384 (30,487 bins) and 24,775 (41,756 bins) genes were distributed inside the TAD of diploid and polyploid pitaya species), respectively. However, 4,045 (3,365 bins) and 2,162 (2,021 bins) genes were located at the TAD boundaries of diploid and polyploid pitaya species, respectively (**Supplementary Table 13 & Supplementary** Fig. 23). Furthermore, we computed GC content distribution in all the bins in TAD and TAD boundaries of both species but we could not find significant differences in the distribution of GC content (**Supplementary Table 14**, **Supplementary Fig. 24**). The ability of TAD boundaries to functionally restrict the genome applied to both active and inactive compartments (A and B) of the genome. TADs are formed by *cis*-acting loop extrusion factors (cohesions) and enriched for the insulator binding protein (*CTCF*) in mammals^33^ however, no protein with insulator function has been reported in plants^19,34^ except *TEOSINTE BRANCHED 1*, *CYCLOIDEA* and *PCF1* (*TCP1*) TFs that has been a promising candidate to replace *CTCF* binding protein^35^. In plants, higher-order chromatin loops are formed between distal regulatory elements and promoters to exert functions, which allows enhancers direct contact with the genes^36,37^. In five plant species (maize, rice, sorghum, foxtail millet, and tomato), TAD showed enriched *cis*-interactions within the same domain and comprised of mammalian TAD-like genetic and epigenetic features changes at the border, which are enriched for active genes, open chromatins, active histone marks and are depleted for transposable elements and DNA methylation^38^. To test the significance of all interaction sites, we used Fit-Hi-C software^39^, and the interaction between the two adjacent bins was discarded. DNA loci within the same chromosomes or between the chromosomes could interact significantly so that domains with different biological functions could perform transcriptional regulation together. AT 10 kb resolution, genome-wide cis-trans significant interaction sites were obtained according to their *p-*value (≤ 0.01) and number of reads supporting the interaction, respectively. The first 25,000 pairs of significant interacting loci were displayed in a circle plot. The integrative analysis of Hi-C and cis-trans regulatory element datasets illustrated 952,744 significant *cis*-regulatory elements (10-kb resolution), and 4,913,539 *trans*-interaction sites were obtained in *S. undatus* (**Fig. 4g**). However, significant 9,634,251 *cis*-interaction loci and 35,7664 *trans*-interaction sites were sorted from *S. megalanthus* genome (**Fig. 4h**).

Generally, TAD boundaries are enriched by structural proteins, *CCCTC*-binding (*CTCF*) factors, and modifications associated with active transcription in mammalian species and are considered to be formed by the active loop extrusion, during which DNA is extruded through a loop until boundary proteins stall the extrusion at the TAD boundaries^19,40^. However, TADs in plants are slightly different than those in animals and are subdivided into four groups with different epigenetic features including active, repressive, polycomb silenced, and intermediate type^38^. The reference genome of diploid and polyploid pitaya species were motif scanned in the JASPAR database (http://jaspar.genereg.net/) using MEME Suite software (http://meme-suite.org/) to obtain Motif annotation information for the whole genome. The number of TAD boundaries with the motif was calculated as the percentage of total TAD boundaries and sorted from large to small percentages. Our results demonstrated that TAD boundaries of the diploid pitaya genome (*S. undatus*) were significantly enriched by *PHYPADRAFT_64121* (90.55%), *RAV1* (90.28%), *AT3G24120* (90.01%), *WRKY60* (89.72%), and *WRKY18* (89.6%) motifs. However, TAD boundaries polyploid pitaya genome (*S. megalanthus*) were enriched by *PHYPADRAFT_64121* (91.21%), *RAV1* (90.41%), *WRKY75* (89.67%), *WRKY60* (89.61%), and *MYB3* (89.53%) motifs (**Supplementary Table 15** & **Fig. 4j**). In rice, DNA GC-rich motifs bound by plant-specific TFs associated with euchromatic epigenetic marks and active gene expression were identified as belonging to the *TCP* family at TADs boundaries^41^. In pepper, TAD-like domain structures were identified and 60% of the pepper genome corresponds to the repressed regions, enriched by repetitive sequences and heterochromatin marks (*H3K9me2*).

### Genome-wide comparative analysis of diploid and polyploid pitaya

We performed a comparative genomic analysis of diploid (red peel) and polyploid (yellow peel) pitaya to investigate the spatial 3D structures, including their compartments and TADs regulating the gene expression. Interaction matrices of diploid and polyploid pitaya were converted into the Z-scores matrices^32^ and obtained the interaction differences by subtracting the matrices and the heatmap was constructed for diploid (**Fig. 5a)** and polyploid pitaya (**Fig. 5b**) species at 200 kb resolution and the interaction matrices of single chromosomes were displayed. At 200 kb resolution, both species were compared to measure the overall differences in distance and attenuation frequency^42,43^. We observed that the distance between both genomes increases as the interaction frequency decreases which indicates the differences and complexity in the spatial structure of both genomes (**Supplementary Fig. 25 & Supplementary Fig. 26**). In genome-wide comparison, we generated saddle plot to measure the degree of euchromatin (compartment A) and constitutive heterochromatin (compartment B) of both pitaya species which showing strong interactions at compartments of A-A ranged from 1.2-1.22, and exhibited somehow strong chromatin interaction in polyploid pitaya than the diploid pitaya (**Fig. 5c**). In addition the ploidy level in yellow pitaya has also effected the B-B compartments and exhibited a higher frequency of contact (3.35) than the diploid pitaya (3.1). As the compartments have plasticity and could convert from compartments A/B state in different biological states by bringing large-scale changes in gene expression^44^. By comparing the homology of the genes in both diploid and polyploid pitaya species, we performed a comparative analysis at 100-kb resolution. Our results displayed that the dynamic 3D genome architecture of both species showed switching of the A/B compartments and reorganization of TADs during their evolutionary process (**Fig. 5d** & **Supplementary Table 16**). Subsequently, we calculated the dynamic changes in TAD structures by the relative signal of TAD boundaries **(Supplementary Table 17).** Repeatability comparison of TAD boundaries demonstrated that 626 TAD boundaries were overlapping in both species among which 1,395 TAD boundaries were unique in diploid pitaya while 1,454 unique TAD boundaries were found in polyploid pitaya (**Fig. 5e**). Genes located within the TAD boundaries of diploid were functionally annotated (**Supplementary Table 13**) and significantly enriched GO entries and KEGG pathways were obtained in both pitaya species. Most of the genes were associated with the cellular process, cell, cell and binding functions and signal transduction pathways (**Supplementary Fig. 27**) (**Fig. 5h**). By using Fit-Hi-C analysis^39^, the interaction between two loci was statistically analyzed. The differences between the *cis*-significant interactions and *trans*-significant interactions in the whole genome of diploid and polyploid species were significant. We also observed 362,158 overlapping significant *cis* (**Fig. 5f**) and 144,1230 *trans*-interactions (**Fig. 5g**) in both pitaya species. Gene expression between the differential compartments of both species illustrated that the maximum number of genes exhibited their expression in A2A compartments (**Supplementary Table 18 and Fig. 5i**). In addition, the distribution of genes between conserved compartments (A2A and B2B) and switching compartments (A2B and B2A) were compared in both species. A2A compartments showed a stronger correlation with gene density than the B2B. Among the switching compartments, B2A exhibited higher gene density than A2B compartments (**Fig. 5j**).

**Fig 5:**
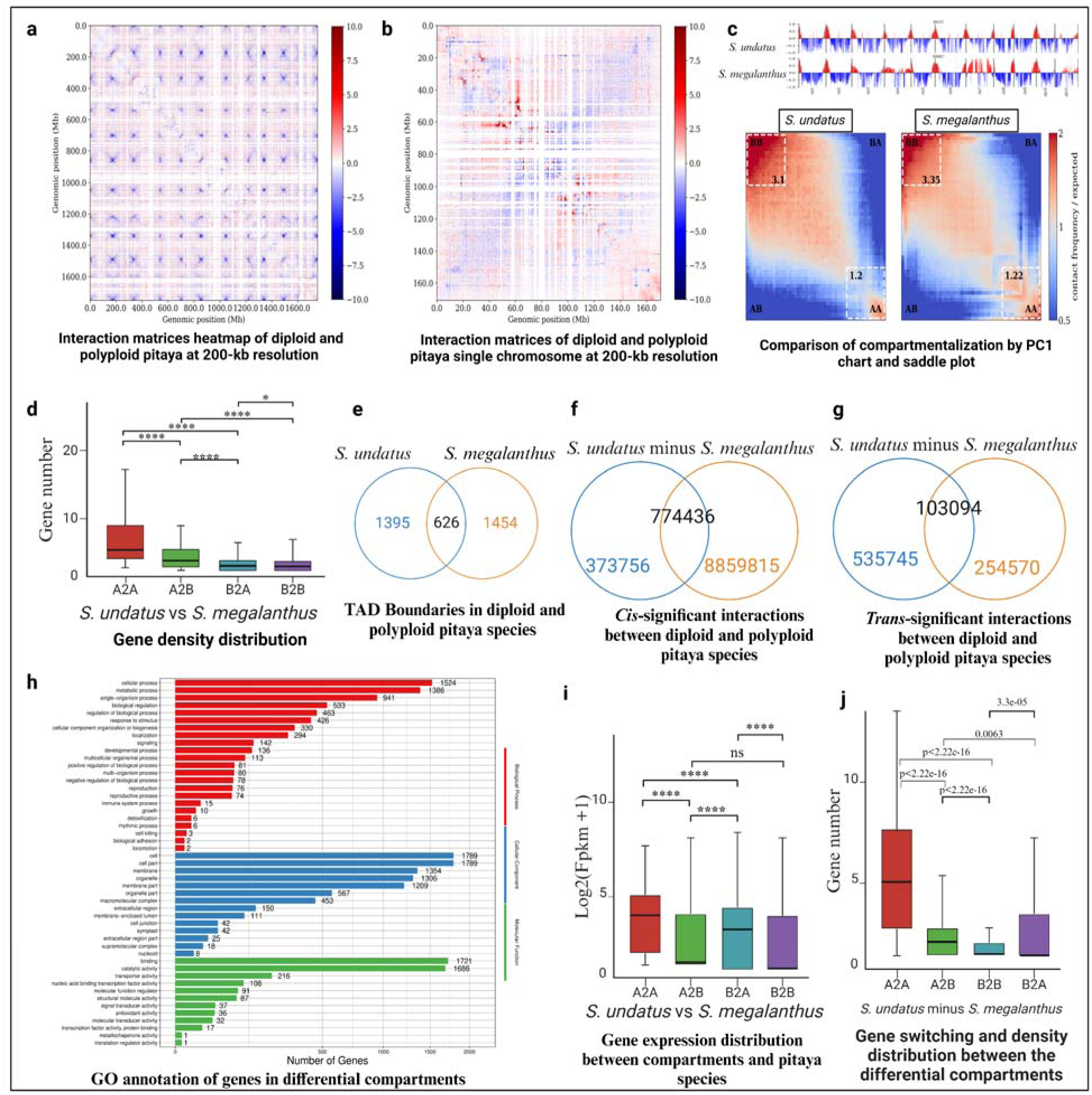
Genome-wide comparative analysis of diploid and polyploid pitaya species. **a)** Interaction heat map at 200-kb resolution exhibits the differences between genomes of polyploid and diploid pitaya (differences in *S. megalanthus* minus interaction differences in *S. undatus*). The color bar including red color indicates that the interaction intensity of *S. megalanthus* was greater than that of *S. undatus*. Blue color denotes the opposite effect, higher value exhibiting the differences in interaction intensity of the locus at diploid and polyploid genomes. Horizontal and vertical axes showed the position of the genome. **b)** Reduced heat map of the interaction of single chromosomes at 200-kb resolution of both pitaya species. The red color bar indicates th interaction intensities of *S. megalanthus* greater than *S. undatus*. A greater value in the blue bar showed the higher differences in the interaction intensity between both species. **c)** Comparison chart and compartment saddle plot. The red color on the comparison chart exhibits compartment A and blue bars represents compartment B. In the saddle plot, the color indicates the interaction signals of compartment AA, BB and AB in diploid and polyploid pitaya. **d)** Gene density distribution between the differential compartments of diploid and polyploid pitaya. The horizontal axis shows four types of compartments while the vertical axis exhibits the distribution of gene density within the region. **e)** Venn plot of common and unique TAD boundaries in diploid (*S. undatus*) and polyploid (*S. megalanthus*) pitaya species. **f)** Venn diagram of *cis*-significant interactions including common and unique ones between two species. **g)** Venn diagram of *trans*-significant interactions including common and unique between two species **h)** GO annotation genes in differential compartments of diploid and polyploid species. A horizontal axis exhibits the number of genes annotated to the GO terms. The vertical axis denotes GO terms in three categories including biological processes, cellular components, and molecular functions. **i)** Distribution of gene expression between genomic compartments. The horizontal axis has been divided into *S. undatus* and *S. megalanthus*. The vertical axis denotes the distribution of gene expression in the compartments of A2A, A2B, B2A, and B2B. Note: The significant test results **** represent 0.0001 ≥ p. However, “ns” represents the not-significant results. **j)** Gene density distribution of differential compartments and gene switching between the compartments of diploid and polyploid pitaya (*S. undatus* minus *S.* megalanthus). The horizontal axis exhibits the four compartment switches and the horizontal axis represents the gene number distribution in conserved compartments (A2A and B2B) and switching compartments (A2B and B2A).

### Structural variations in diploid and polyploid pitaya genomes

Polyploids are formed by the duplication of a single genome (autopolyploid) or the combination of two or more sets of chromosomes to design a new species (allopolyploids)^45^ and undergo rapid structural and functional alterations. SVs play an important role in species evolution and assist in climate change adaptation. We compared the autotetraploid yellow pitaya genome with the available genome of closely related biological species of red-peel white-flesh pitaya (*S. undatus*). In autotetraploid pitaya, we found 41,409 and 82,394 presence-absence variations (PAV), 1245 inversions, 15,725 translocations, 23,325,494 SNPs, 2,294,810 InDels and 19,928 copy number variations (CNVs). Most of the PAVs (31,506-66,158), inversion (710), translocations (13,533), and CNVs (14,890) were found in the intergenic region of the yellow pitaya genome (**Supplementary Table 19**). These varying numbers of SVs in yellow pitaya produced genetic and phenotypic differences as compared to other pitaya species (**Fig. 1A)**. Notably, the diploid genome of pitaya has had extensive syntenic relationship with yellow pitaya. Heterozygosity and extensive accumulation of duplications, 2533 frameshift insertions, translocations at genomic regions of 3′UTR (11), 5′UTR(3), downstream (230), exonic (1017), intergenic (13,533), intronic (639), splicing (1) and upstream (280) regions of autotetraploid yellow pitaya, might have led to an increase in the genome size.

Large inversions were found on Chr 1, Chr 2, Chr 3, Chr 4, Chr 5, Chr 7, Chr 8, and Chr 9. On the other hand, a large number of translocations were noticed on almost all chromosomes of autotetraploid pitaya. We found genome duplication events on almost all chromosomes of yellow pitaya, but very rare duplicated events were observed on Chr 10 (**Fig. 6**). Homologous exchanges occur by replacing one genomic segment with another copy number or ancestrally duplicated regions. Traditionally, autopolyploids are considered to arise within a single species by doubling the homologous genomes (AAAA)^46^. As the high-density SNP could be difficult in the heterozygous genome of tetraploid yellow pitaya, our analysis revealed a descending count of SNPs ranging from 25,752,287 SNPs on Chr 1 and the lowest 15,658,201 SNPs on Chr 11. In addition, Chr 1 exhibited high SNP density approaching 48Mb and between 112 to 128 Mb regions possibly indicative of high recombination or potential adaptive evolutionary hotspots, and consistently low-density area on Chr 5, 8, and 11 representing evolutionarily conserved regions. Interestingly, the terminal ends of most of the chromosomes generally showed a trend of reduced SNP density, potentially indicating conserved regions or structural elements for chromosomal stability (**Supplementary Fig. 28**). We found that the genome size of diploid pitaya was estimated as 2n=1.41 Gb^26^. However, the yellow pitaya genome is much bigger than the estimation (i:e, 7.16 Gb) indicating the extensive genome organization and diverse SVs as compared to their diploid parent. Clonal propagation^24^ or somatic mutations^25^ might also have a role that results in a heterozygous manner within a clone of autotetraploid pitaya. Synthetic autopolyploids of *Phlox drummondii* exhibited immediate loss of DNA (up to 25%) in the third generation which led to a substantial decrease in genome size and an increase in seed set in plants^47^. Rapid deletion of DNA sequences from autopolyploid genomes followed by genome doubling, possibly sustains the bivalent pairing and part of the adaptive measures. Therefore, it is noteworthy to highlight that DNA loss is not indeed necessary in every autopolyploid. In Arabidopsis, no structural change and DNA loss were reported in the first generation of synthetic autopolyploid^48^. Genome size increases also arise from the amplification repeats, independent of polyploidization^49^. Initially, we estimated the genome size of yellow pitaya as 5.14 Gb, but we found the genome size of yellow pitaya n=1.79 Gb (2n=4x=7.16 Gb), which was much bigger than the estimation. Antagonistic to previous reports, autopolyploid retain or lose the DNA segments after ploidy events. But in the case of yellow pitaya, it seems that the genome faces a lot of duplication events or gone through a series of crossing and hybridization (see **Supplementary Figure 5** for more arguments) events which might have led to an increase in the genome size as compared to the unknown ancestral parent.

**Fig. 6:**
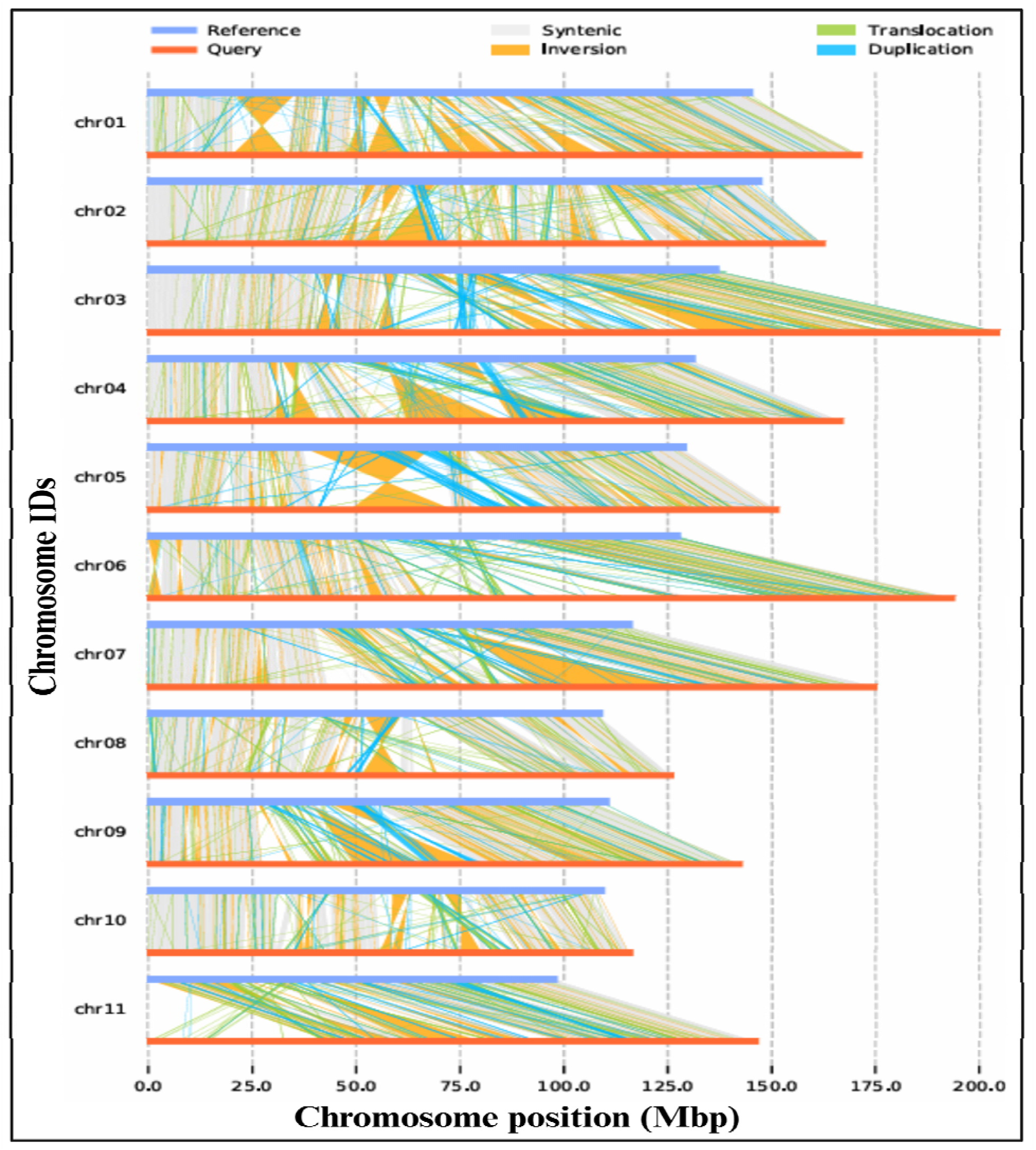
Genome structural variations in autotetraploid polyploid pitaya. The blue bar exhibits the reference genome of yellow pitaya but the orange bar represents the diploid genome of red-peel pitaya. The blue bar (reference sequence) denotes the genome of *S. undatus*. The orange bar (query sequence) represents the genome of yellow pitaya (*S. megalanthus*).

### Genome-level chromatin accessibility of diploid and polyploid pitaya

Assay of transposase-accessible chromatin sequencing (ATAC-seq) has been widely used technique to accurately map accessible chromatin of the genome, and *cis*-regulatory elements, and uncover the underlying mechanism of gene regulation^50^. In this study, we performed ATAC-seq by following the sucrose precipitation method^51^, and Illumina-PE150 sequencing was performed for both diploid and polyploid pitaya species, and 0.375 billion clean reads were generated (**Supplementary Table 20**). After filtering the data, we proceed with further bioinformatics analysis to identify the enrichment of ATAC-seq reads after considering the quality parameters including base content distribution (**Supplementary Fig. 29**), base mass distribution (**Supplementary Fig. 30**), GC content distribution, and insert length distribution (**Supplementary Fig. 31 & Supplementary Table 21**). Most of the ATAC-seq reads were enriched at the transcriptional start sites (TSS) (**Fig. 7a**) in diploid and polyploid pitaya genomes. In our results, we observed ∼43186 ATAC-seq peaks in diploid pitaya and detected ∼30,276 in polyploid pitaya by using MACS3 software (**Supplementary Table 22**). We compare the overlap of the biological replicates (**Fig. 7b**) and the distribution of the peaks was analyzed at 3 kb upstream and 3 kb downstream at the TSS region. The enriched region of the genome was selected to visualize the peak assay where the ATAC-seq data signal was significantly enriched in both pitaya species (**Supplementary Fig. 32**). In addition, the distribution trend of ATAC-seq signal in the gene body region (from TSS to TES) and the chromatin openness near the TSS regions of diploid and polyploid pitaya was plotted (**Supplementary Fig. 33**). The peaks detected between the biological repeats of both pitaya species were combined to obtain the peak set within the group. In the results, we obtained 125,321 peaks for diploid pitaya and 30,276 peaks for polyploid pitaya. In addition, peaks from both species were compared, and detected 13,136 overlapping peaks (**Supplementary Fig. 34**). By comparing the signal values of the ATAC-seq data between the diploid and polyploid pitaya species, a region with significant differences in signal values was detected using the R package DiffBind and named as differentially accessible region (DAR). To compare the biological differences between the two species including the chromatin openness that regulates the downstream gene regulatory network and leads to the ultimate phenotype of the pitaya plants, we compared the DAR of both species. In results, we obtained a total of 13,977 DARs between the two pitaya genomes including 3,355 gains of DAR and 10,622 loss of DARs. The signal value of DAR of each biological repeat was clustered (**Figure 7b**) which showed a large number of differentiations in DAR between both pitaya species. To obtain the position, the DARs were annotated on the genome of diploid and polyploid pitaya, and we found that only >5% of the DARs increased the chromatin accessibility on the promoter region (gain DAR) and >24% of the DAR decreased the chromatin accessibility at promoter region (loss DAR) (**Fig. 7c**). Therefore, we paid more attention to the DAR associated with the promoter region by comparing both pitaya species. The results revealed that 159 genes (gain DARs) and 2362 genes (loss DAR) were associated with the promoter regions of pitaya species (*S. undatus* vs *S. megalanthus*). The function of the distributed genes and their coordinated pathways were determined through GO and KEGG pathway analysis (**Supplementary Fig. 35**-38) which revealed genes mainly enriched in response to “cation transport, ATP metabolic process, and carbohydrate biosynthetic process in gain of DAR. The top20 GO terms contained in the promoter regions of the genome were annotated and clusterProfiler was used to display in word-cloud which revealed enriched GO terms associated with metabolites, response to a toxic substance, ATP synthesis, triphosphate biosynthesis, herbicide response, and energy. By contrast, GO terms enriched in loss of DAR were comprised of glucose pathways, circadian, stimulus-response, and photosynthesis in the promoter region of both pitaya species (**Fig. 7d**). Using the JASPAR database and MEME suit tool, motif scanning, and enrichment analysis of gain and loss of DAR was carried out, and we found 109 enriched motifs in gain of DAR and 425 motifs in loss of DAR. We noticed that *WRKY75/40*, *UIF1*, A*T5G02460*, *GATA10*, *OBP3/4* and *PHYPADRAFT-140773/153324* were the most enriched motifs for gain DAR. Antagonistically, we found that *ABI5*, *EmBP-1*, *TCP8*, *bZIP910*, and *AP1* were enriched motifs for loss DAR (**Supplementary Fig 39 & Fig. 40**). We further investigated how many genes associated with DAR or differentially expressed genes (DEGs) overlapped in the promoter region of both pitaya species. Binding of TFs to DNA sequences usually activates gene expression which has a strong correlation between gene expression and the chromatin openness in the vicinity of the genes. By comparing DAR-associated genes and DEGs of both pitaya species (*S. undatus* vs *S. megalanthus*), we found 15 genes overlapped for gain-DAR vs up-DEGs (**Fig. 7e**), 6 genes for gain-DAR vs down-DEGs, 146 genes for loss-DAR vs up-DEGs and 296 genes for loss-DAR vs down-DEGs (**Supplementary Fig. 41**). Expression analysis of genes associated with gain DAR showed significantly higher expression in the promoter region of *S. megalanthus* as compared to *S. undatus* (**Fig. 7f**). ATAC-seq files of both species and RNA-seq data were jointly visualized by integrated genome viewer (IGV v2.4.18)^52^ browser to view the results of any region of interest at the genome-wide level. We discovered significant differences in chromatin accessibility between diploid and polyploid pitaya species (**Fig. 7g**).

**Fig. 7.**
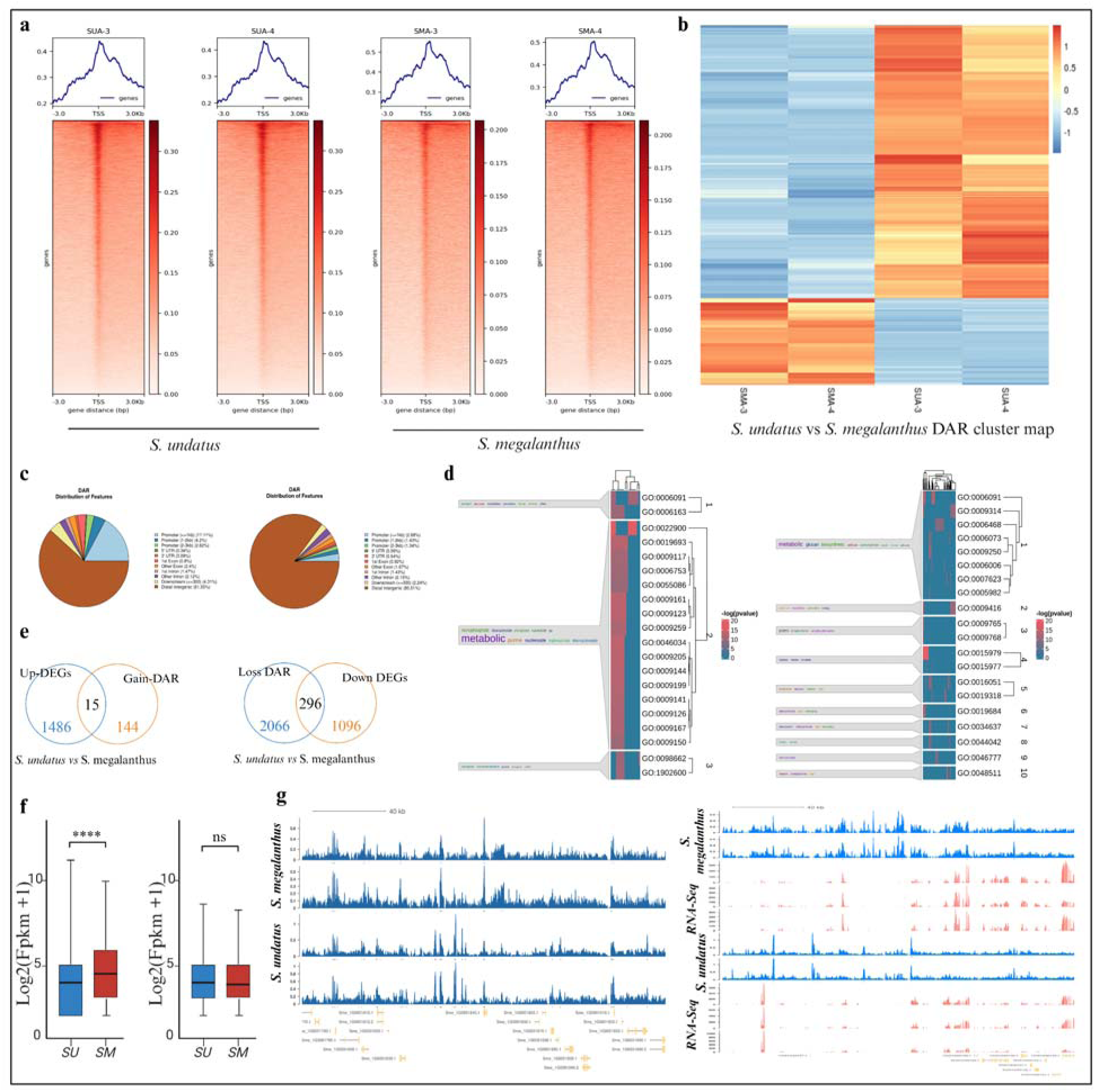
The ATAC-seq analysis on whole genomes of diploid and polyploid pitaya. **a)** Signal enrichment plot of ATAC-seq in the TSS region of *S. undatus* and *S. megalanthus* genome. The horizontal axis of the heat map is 3kb upstream and downstream of TSS and the vertical axis is the gene. The horizontal axis of the trend chart is the range of 3KB upstream and downstream of the TSS, and the vertical axis is the signal enrichment at the genomic position. **b)** DAR cluster map of the diploid and polyploid pitaya. The horizontal is the name of the pitaya species and the vertical axis is the DAR obtained by comparing the two species. The color in the figure showed the magnitude of the signal values of the DAR in each species. **c)** Distribution of gain of DAR and loss of DAR. The map showed the location of DAR on genomes and the proportion of DAR on the location. **d)** GO terms in Gain DAR and loss DAR-associated genes located on the promoter region. Genes are clustered according to their term and their function is displayed on the left side. **e)** Comparison of DAR-associated genes and differentially expressed genes to identify the overlapping genes in the promoter region. **f)** Box plot of DAR-related gene expression. **g)** Visuliation of multiomics data revealed significant differences in chromatin accessibility between diploid and polyploid pitaya species.

### Comprehensive analysis of chromatin architecture of diploid and polyploid pitaya species to develop unique phenotype

To identify the tissue-specific differential genes and open chromatin structure of the pitaya genome, we link the Hi-C, and tissue-specific pericarp RNA-seq data of both pitaya species by following the Liu and colleagues^53^. As the subcompartment switching is correlated with changes in expression level, we classified all 40-kb genomic bins into three groups based on changes in the subcompartment. All 11 chromosomes were classified into compartment A (open chromatin and high RNA expression region), compartment B (closed chromatin and low RNA expression), TAD, and TAD boundaries in red-peel pitaya-*S. undatus* and yellow-peel pitaya-*S. megalanthus* species. In results we discovered 16,685 genes located in compartment A and 5,279 genes in compartment B of yellow pitaya (**Supplementary Table 23)**. Similarly, we found 17,759 genes expressed in the compartment of A and 5,978 genes in compartment of B in red-peel pitaya (**Supplementary Table 24)**. By separating the genes of pitaya species in active and inactive compartments, local chromatin active domains i:e TADs and TAD boundaries were identified as enriched with insulator-binding proteins. Results in, we found 20,453 genes in TADs and 2,264 genes in the TAD boundaries of yellow pitaya (**Supplementary Table 25**). Similarly, we predicted that 22,603 genes in TADs and 1,436 genes in TAD boundaries were expressed red-peel pitaya (**Supplementary Table 26**). This methodology directly assisted us in pointing out the genes involved in the betalains pathway producing the yellow color in *S. megalanthus* and the red color *S. undatus* fruit. We first tested whether the “down” and “up” bins overlap with more DEGs and then correlated this classification with fold changes in expression level between tissues. We adopted the following approaches for dealing with replicates including evaluating the consistency of results between independent analysis completed for each replicate and using only the chromatin features shared by both replicates for the analysis. To identify a reorganization of TADs and loops that were correlated with alteration in expression in pitaya pericarp, we divided the genome into compartments (A/B). After compartmentalization, genes were identified into TADs and TAD-boundaries and checked whether it is located on inter-TADs or TAD-boundaries. We correlated this classification of TAD features in gene expression profiles, linked the genes with annotation of the genome, and predicted the genes playing their role in the betalains pathway.

We found that the genes involved in betalains biosynthesis in diploid pitaya showed higher sequence similarity with yellow pitaya genomes at the same chromosome numbers except for a few genes but varied in copy number (**Supplementary Table)**. In addition, most of the genes exhibited differential expression in diploid and polyploid pitaya, highlighting the morphological differences between both pitaya species. Notably, we found few specific genes in diploid red peel (*HU10G00264*, *HU09G01263*, *HU07G00551* and *HU08G02124*) and polyploid yellow peel (*Sme_10G0002470*, *Sme_9G0013340*, *Sme_7G0004800* and *Sme_8G0020650*) pitaya involved in the shikimate pathway regulating the betalains biosynthesis (**Fig 8)**. The biosynthesis of betalains involves the conversion of tyrosine by the predicted genes of diploid (*HU02G00163*, *HU10G00810*, *HU10G00173*, *HU07G00908*, *HU02G00811*, *HU05G00517* and *HU11G01112*) and polyploid pitaya (*Sme_2G0001630*, *Sme_10G0007740*, *Sme_10G0001580*, *Sme_7G0008480*, *Sme_2G0008230*, *Sme_5G0004900* and *Sme_11G0010540*), a precursor molecule, through two enzymatic reactions. Specifically, tyrosine is synthesized from prephenate^54^ which was regulated by genes regulated in diploid (*HU01G02069* and *HU07G00319*) and polyploid (*Sme_1G0021320* and *Sme_7G0002630*) pitaya by the action of prephenate dehydrogenase, and from arogenate by arogenate dehydrogenase (*ADH*)^54^. Furthermore, the joint analysis predicted the genes involved in cytochrome P450 enzymes hydroxylate tyrosine, producing the *L-DOPA* in diploid (*HU10G01783*, *HU08G01572* and *HU08G01571*) and polyploid pitaya (*Sme_10G0019300*, *Sme_8G0014610*, *Sme_8G0014620*) (**Fig. 8a)**. *L-DOPA* undergoes ring-opening oxidation, a process facilitated by *L-DOPA 4,5-dioxygenase* (*DODA*), leading to the creation of the intermediate molecule known as *4,5-seco-DOPA*^55^. Our results revealed the genes in diploid (*HU07G00239* and *HU07G00240*) and polyploid (*Sme_7G0001870*, *Sme_7G0001880*) species regulating and the conversion of *DOPA* to *cyclo-DOPA*. For the synthesis of red violent betacyanin, we predicted the genes (*HU06G02516*, *HU04G00198*, *HU07G01678*, *HU07G00475*) in red-peel pitaya (*S. undatus*), that may have the potential to engage in the regulatory pathways and produces enzymes that spontaneously conjugate with the imino group of *cyclo-DOPA*, culminating the formation of betalain pigments i.e., red-purple betacynanin^55^. Antagonistically, in autotetraploid pitaya, we revealed that the *Sme_6G0025350*, *Sme_4G0001820*, *Sme_7G0017600* and *Sme_7G0004100* genes belong to *SmeABA2*, *SmeFG2/3* and *SmeHD-ZIP* families playing the potential role to spontaneously condense the betalamic acid with the amino group to produce yellow betaxanthins rather than with *cyclo-DOPA* derivatives^56^. By comparing the gene expression of the betalains pathway in both species, we observed major changes in gene expression regulating the betalains pathways and expressed across the pericarp of pitaya fruit. The genes directly regulating the *shikimate* pathway (*Sme_10G0002470* and *Sme_7G0004800*) in yellow pitaya exhibited 3-fold higher expression as compared to genes (*HU10G00264* and *HU07G00551*) expressing in pericarp red-peel pitaya. We found non-significant expression in genes of both species regulating prephenate in the betalains pathway (**Fig. 8b)**. In a betalains pathway, we identified genes with varying degrees of expression in both pitaya species, leading two contrasting phenotypes. We found the genes belonging to *CYP76AD1* that regulate the enzyme to oxidase tyrosine into L-DOPA and produce betacyanins in diploid pitaya showed 2-fold higher expression (*HU07G00500*-*CYP76AD1*), as compared to the genes (*Sme_7G0004430*-*CYP76AD1*) in the fruit pericarp of yellow pitaya (**Fig. 8b).** In addition, the homologs copy in diploid pitaya (*HU01G02117*-*CYP736A12*) exhibited 1.2 fold higher expression that the gene (*Sme_1G0021900*-*CYP736A12*) in yellow pitaya. In addition, we found that a homologous copy of the *CYP736A12* gene is located in compartment A and compartment B of the yellow pitaya genome. During the evolutionary and polyploidization process of yellow pitaya, duplication of *CYP736A12* gene copy in compartment A/B and reduced expression of *CYP36A12* genes in yellow pitaya as compared to red-peel pitaya ultimately led to the loss of capacity to oxidase the tyrosine into *L-DOPA* and instead produced betaxanthin.

**Figure 8.**
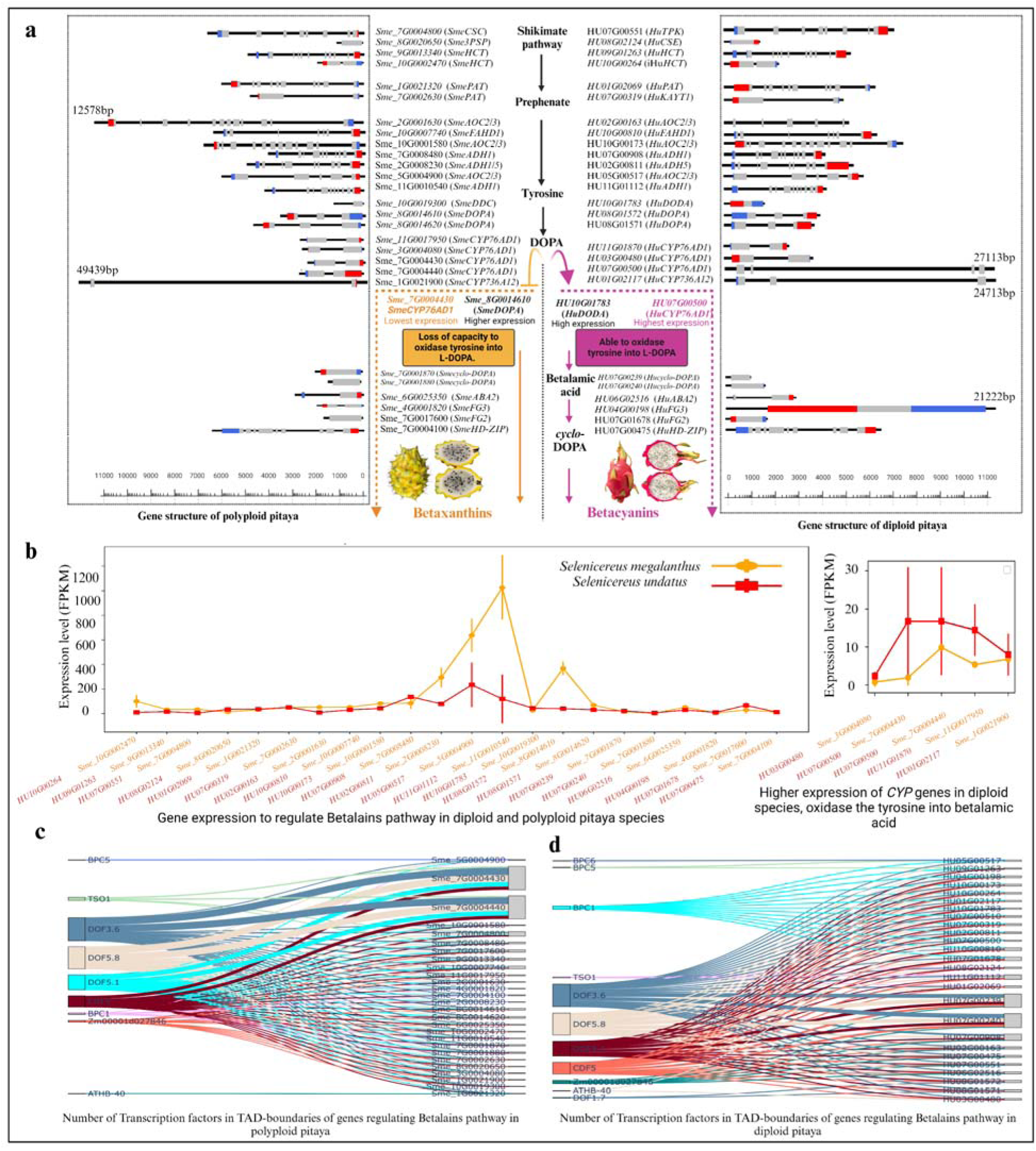
Linking tissue-specific gene expression to open chromatin of diploid and polyploid pitaya. **a)** Genes involved in betalains biosynthesis in diploid pitaya showed higher sequence similarity with yellow pitaya genomes at the same chromosome numbers except for a few genes and also varied in copy number. Genes are exhibited that may have a potential role in red-peel pitaya (*S. undatus*), to engage in the regulatory pathways and produce enzymes that spontaneously conjugate with the imino group of *cyclo-DOPA*, culminating in the formation of betalain pigments i.e., red-purple betacyanin. In autotetraploid pitaya, genes belonging to *SmeABA2*, *SmeFG2/3* and *SmeHD-ZIP* families play the potential role of spontaneously condensing the betalamic acid with the amino group to produce yellow betaxanthins rather than with *cyclo-DOPA* derivatives **b)** The genes directly regulating the *shikimate* pathway (*Sme_10G0002470*, *Sme_7G0004800*) in yellow pitaya exhibited 3-fold higher expression as compared to genes (*HU10G00264*, *HU07G00551*) expressing in pericarp red-peel pitaya. A homologous copy of the *CYP736A12* gene is located in compartment A and compartment B of the yellow pitaya genome. During the evolutionary and polyploidization process of yellow pitaya, duplication of *CYP736A12* gene copy in compartment A/B and reduced expression of *CYP36A12* genes in yellow pitaya as compared to red-peel pitaya ultimately led to the loss of capacity to oxidase the tyrosine into *L-DOPA* and instead produced betaxanthin. **c)** TFs identified from the 40-Kb region of TAD boundaries and top 5 TFs are presented in polyploid pitaya **d)** TFs identified from the 40-Kb region of TAD boundaries in diploid pitaya and top 5 TFs are shown in the figure.

These observation raises the question, how chromosome-chromosome folding, chromatin structure, TADs borders, active (A) and inactive compartments (B) are linked to produce two different phenotypes in pitaya species? Does motif count in TAD boundaries affect the gene expression in TADs? As earlier, in our results, we showed the major differences in genomic organization, compartmentalization, and differences in TADs of both species. How these changes affected the TAD boundaries or vice versa led to the changes in gene expression and produced two different phenotypes. Here, we asked which TF or motifs are associated with these TAD boundaries of both pitaya species that led to changes in gene expression. How do the long-term evolutionary rearrangements of TAD boundaries alter gene expression? As the TAD boundaries were enriched with TFs and housekeeping genes. We analyzed the gene position in TADs and mapped the TAD boundaries on diploid and polyploid pitaya species genomes (**Supplementary Tables 23 & 24**). We identified the 40-Kb region of TAD boundaries for diploid (**Supplementary Table 27)** and polyploid (**Supplementary Table 28**) species in between the TADs in which our genes of interest were located (**Supplementary Table 29**), and playing a role in the betalains pathway. We identified all TFs from the 40-Kb region of TAD boundaries and motif count was computed to explore the top 5 TFs from TAD boundaries of both pitaya species (**Supplementary Table 30).** We found variations in motif types in TAD boundaries of diploid and polyploid pitaya that were regulating the changes in expression profile and ultimately producing the yellow and red-peel fruit color. Our results revealed that expression of the genes regulating the betalains pathway was highly correlated with the motif count in the TAD boundaries. As **Figure 8b** exhibited, those genes exhibited extensive variation in expression, and showed higher differences in motif counts in TAD boundaries. Among all the genes, the *Sme_11G0010540* gene exhibited higher expression within the TADs and lower number of TF in TAD-boundaries of yellow pitaya as compared to the red-peel pitaya (**Fig. 8c)**. However, *Sme_8G0014610* gene showed lower level gene expression within the TAD and a higher number of TFs in TAD boundaries in yellow pitaya as compared to the red-peel pitaya. In addition, we found that *CYP76AD1* genes (*Sme_7G0004430*, *Sme_7G0004440*) exhibited the highest number (same TFs repeated) of TFs [*DOF3.6* (11890), *DOF5.8* (11650), *DOF5.1* (7780), *CDF5* (5605) and *TSO1* (2220)] but showed lowest genes expression in yellow pitaya as compared to the red-peel pitaya (**Fig. 8d)**. That’s why these genes lost the capability to oxidase the tyrosine and produced the betaxanthins. These findings suggest that TAD-boundaries have had the role of regulator and repressive at once and mechanistically control the genes within the TADs. Furthermore, we found that the *Hu01G02117* gene of diploid pitaya is located in compartment A of the genome. In contrast, its counterpart gene (*Sme_1G0021900*) of yellow pitaya is located in compartment A and compartment B of polyploid pitaya. This might be a reason for the low expression of *CYP736A12* genes in polyploid pitaya that led to the loss of capacity to oxidase the tyrosine and ultimately produced yellow color fruit.

## Discussion

Autopolyploidy in yellow pitaya and whole genome duplications might be the reason that leads to the phenotypic difference of morphological and metabolic changes in tetraploid pitaya including the fruit color, cladode thickness, size of fruit, and seed size. Chromosome number, genome size, genes, and their expression evolve following polyploidization of the genome enabling the diversification of plant species and novel traits. In yellow dragon fruit, polyploid genome and gene redundancy seem to originate from unreduced gametes of diploid plants. Most of the formation of the unreduced gametes is associated with temperature instability (low/high) in various organisms, including plants^57–59^, fish, and amphibians^60^. Production of unreduced gametes is governed by a few genes that are heritable and increase as the environmental stress increases. The evolution of polyploid has also been led by grafting. The *Cacataceae* family can easily be grafted using centuries-old traditional methods. Nuclear genomes can move from rootstock to scion. As a result of this movement, some cells became polyploid which led to a fully grown shoot that have fertile flowers^61^. In our results, we found the difference of 3% genome identified by K-mer analysis as allopolyploid might be aligned with the prediction of Tel-Zur and colleagues (2022)^4^ narrowly allopolyploid and 64% heterozygosity in the genome predicted by our Smudgeplot analysis, might be the reason for grafting, series of crossing of intraspecific hybrids originated from multiple origins^10,46^ or lateral gene transfer (LGT) of DNA without sexual reproduction. Maybe, autotetraploid yellow pitaya involved crosses of two diverged populations of the same species with genetically distinct, but structurally similar chromosomes by forming an “interracial autopolyploid”. These genomic changes and increased genetic diversity alter the morphology, physiology, and ecology of the yellow pitaya, gaining further contribution to speciation and adaptation to environmental changes. Multiple origins of polyploid are common and genetic diversity has increased due to different maternal lineages^46^. Multiple alleles per locus allow polyploids to thrive in a diverse and harsh environment, making them highly tolerant to stress factors and allowing them to adapt and persist across heterogeneous landscapes in the long run^46^. Many regulatory networks are known to be highly dosage-dependent and duplicated genomes with multiple alleles provide extra dosage. In addition, selection under domestication may favor the loss or gain of particular genes due to human interferences^62^. In contrast, the selection of a few loss of function mutations is regarded as beneficial in breeding and crop improvement as *GA20-OX* laid the foundation of dwarf wheat and Green-Revolution. In the synthetic autopolyploid of *Phlox drummondii*, immediate loss of 17-25% of total DNA was observed up to the third generation^47^. The decrease in genome size not only enhanced the seed set (30-66%) but also helped in the stabilization of tetraploid. However, more research is needed to document the patterns and genome loss/gain under the domestication of pitaya and assess the utility of the genes that were exposed through inbreeding and others remain as cryptic. So the possible explanations for the establishment of autopolyploid have been proposed including vegetative reproduction and perennial habit along with the outcrossing matting system^63,64^ and we proposed three assumptions of yellow pitaya WGD process as mentioned in **Supplementary figure 5**.

In diploid and polyploid pitaya species, the unusual class of pigments known as betalains have accelerated research discoveries and shed light on the evolution and biosynthesis pathways fast-tracked by omics data and synthetic biology. Plant pigmentation, particularly in *Caryophyllales*, has captivated researchers due to the exclusive production of betalains or anthocyanins, but never both. Betalains, a prominent class of plants’ secondary metabolites, hold considerable significance across various facets of our lives. To identify the underlying mechanism of phenotypic differences between yellow and red-peel pitaya, 3D folding of genomic regions and packaging in chromatin loops, TADs, A/B compartments^65^, and chromosome territories (CTs)^66^ were explored. Using the insulation score method^32^, TAD boundaries were computed in both pitaya species. In total, 3376 TADs were identified in the diploid pitaya genome and 2031 TADs were predicted in the polyploid pitaya genome with the mean length of TAD 401056 bp/pc and 861897 bp/pc, respectively. TADs are hard-wired features of chromosomes, and a group of TADs can form larger A and B compartments. TADs play an important role in organizing the genomic functional domains and regulating the genes in spatial and temporal patterns^35^. The loci within the same TAD interact with each other frequently, and then they interact with other adjacent TADs. These self-interacting domains are separated by boundaries and serve as a barrier to the spread of activity within the genome^19^. Our results revealed that the dynamic 3D genome architecture of both species showed switching of the A/B compartments and reorganization of TADs during their evolutionary process. Repeatability comparison of TAD boundaries demonstrated that 626 TAD boundaries were overlapping in both species among which 1,395 TAD boundaries were unique in diploid pitaya while 1,454 unique TAD boundaries were found in polyploid pitaya. We also observed that 24,384 (30,487 bins) and 24,775 (41,756 bins) genes were distributed inside the TAD of diploid and polyploid pitaya species), respectively. However, 4,045 (3,365 bins) and 2,162 (2,021 bins) genes were located at the TAD boundaries of diploid and polyploid pitaya species, respectively. The ability of TAD boundaries to functionally restrict the genome applied to both active and inactive compartments (A and B) of the genome. TADs are formed by *cis*-acting loop extrusion factors (cohesions) and enriched for the insulator binding protein (*CTCF*) in mammals^33^, however, no protein with insulator function has been reported in plants ^19,34^ except *TEOSINTE BRANCHED 1*, *CYCLOIDEA* and *PCF1* (*TCP1*) TFs that have been a promising candidate to replace *CTCF* binding protein^35^. Moreover the results revealed that TAD boundaries of the diploid pitaya genome (*S. undatus*) were significantly enriched by diverse motifs including *PHYPADRAFT_64121* (90.55%), *RAV1* (90.28%), *AT3G24120* (90.01%), *WRKY60* (89.72%), and *WRKY18* (89.6%). However, TAD boundaries polyploid pitaya genome (*S. megalanthus*) was enriched by *PHYPADRAFT_64121* (91.21%), *RAV1* (90.41%), *WRKY75* (89.67%), *WRKY60* (89.61%), and *MYB3* (89.53%) motifs. In plants, higher-order chromatin loops are formed between distal regulatory elements and promoters to exert functions, which allows enhancers to direct contact with the genes^36,37^. In five plant species (maize, rice, sorghum, foxtail millet and tomato) TAD showed enriched *cis*-interactions within the same domain and comprised of mammalian TAD-like genetic and epigenetic features changes at the border which are enriched for active genes, open chromatins, active histone marks and are depleted for transposable elements and DNA methylation^38^. Plant boundaries lack *CTCF* boundary homologs and TAD boundaries, but are enriched with *TCP* and *bZIP* TFs binding sites, raising the possibility of involvement in TAD formation^67^. In Arabidopsis, the disruption of the loop between the promoter and transcription termination site results in a decrease *FLC* gene expression^68^. In *Marchantia polymorpha*, *TCP1* knockout mutants, genes located in *TCP1-rich* TADs exhibited larger changes in expression levels as compared to the genes outside of the TADs. However, neither *TCP1-bound* TAD borders nor *TCP1-rich* TADs exhibited changes in chromatin pattern that posits that *TCP1* is not essential for TAD formation in *Marchantia polymorpha* ^69^. In rice, DNA GC-rich motifs bound by plant-specific TFs associated with euchromatic epigenetic marks and active gene expression were identified as belonging to the TCP family at TADs boundaries^41^. In pepper, like mammals, TAD-like domain structures were identified. In total, 60% of the pepper genome corresponds to the repressed regions that were enriched by repetitive sequences and heterochromatin marks (*H3K9me2*). Pepper TADs-like domains were enriched by genes that may act to connect genes via gene-to-gene loops leading to the spatial clustering of active genes^70^.

Within the pitaya TADs, we found that the genes involved in betalains biosynthesis of diploid pitaya showed higher sequence similarity with yellow pitaya genomes at the same chromosome numbers except for a few genes but varied in copy number. Additionally, most of the genes exhibited differential expression in diploid and polyploid pitaya, creating differences in both pitaya fruit morphology. We found a few specific genes in diploid red peel (*HU10G00264*, *HU09G01263*, *HU07G00551* and *HU08G02124*) and polyploid yellow peel (*Sme_10G0002470*, *Sme_9G0013340*, *Sme_7G0004800* and *Sme_8G0020650*) pitaya involved in the shikimate pathway that regulates the betalains biosynthesis. We identified all TFs from TAD boundaries and motif count was computed to explore the top 5 TFs from TAD boundaries of both pitaya species. We found variations in motif types in TAD boundaries of diploid and polyploid pitaya that were regulating the changes in expression profile and ultimately producing the yellow and red-peel fruit color. Our results discovered that expression of the genes regulating the betalains pathway was highly correlated with the motif count in the TAD boundaries.

The availability of the diploid pitaya (*S. undatus*)^3,26^ the genome has expanded the toolbox and trait improvement efforts. Nonetheless, the lack of the reference genome of polyploid pitaya (*S. megalanthus*), the controversial debate over ploidy level, significantly hinders the studies focused on its evolutionary process and potential for genetic improvement of yellow peel pitaya. High-quality reference genomes and annotation provide fundamental resources for plant breeding and opportunities for crop improvement by identifying trait-specific genes. In this study, the first high-quality reference genome with 11 chromosomes was generated for yellow pitaya fruit using PacBio-HiFi-Seq. In addition, Hi-C, ATAC, and RNA-seq were also carried out for diploid red pitaya (*S. undatus*) and autopolyploid yellow pitaya (*S. megalanthus*). Diploid and polyploid pitaya genomes were compared to identify the SVs, A and B compartments, TADs, and their associated TAD boundaries to identify the genes regulating phenotypic traits. Linking the 3D genome structure of pitaya and transcriptome sequencing of pitaya pericarp tissues assisting us in identifying the genes regulating phenotypic variations in pitaya species. These genomic resources enabled us to potentially associate phenotypic differences of diploid and tetraploid pitaya species with the active TADs of the genomes. In short, our results advance our understanding of optimizing and innovating plant breeding strategies to enhance crop traits by exploiting genomic architectural domains and meeting sustainable food security targets.

## Materials and Methods

### Sample preparation for sequencing

Plant material of yellow pitaya [(*Selenicereus megalanthus*) previously known as *Hylocereus megalanthus* (K. Schum. Ex Vaupel) Ralf Bauer] and red peel white pulp pitaya [(*Selenicereus undatus*- Britton & Rose), previously known as *Hylocereus undatus*)] for all types of sequencing were collected from Hainan-Shengda Modern Agriculture Development Company (latitude/longitude 19°N/110°E), Qionghai, Hainan, China.

### DNB sequence survey

The young cladode of the yellow pitaya (*S. megalanthus*) was collected from the field for a DNA nano Ball (DNB) sequence survey. Samples were prepared for the DNB survey using the Beijing Genomics Institute (BGI) sequencing group platform (zero-PCR DNBSEQTM). Total DNA was extracted from fresh cladodes using the standard cetyltrimethylammonium bromide (CTAB) method. After checking the quality and quantity of the DNA, sequencing was carried out at the BGI DNBSEQTM platform. Samples were randomly interrupted by an ultrasonic system (Covaris) and fragments were obtained around 350 base pairs (bp). The DNA fragments were repaired by base A at the 3 end. The ligated library was isolated, and the circle library was generated by rolling circle amplification (RCA) to DNB. The library was sequenced, and raw data was obtained in FASTQ format. Using fastp-v0.20.04 software, clean reads were generated after removing the adapter and low-quality reads using Trimmomatic software (v 0.38)^71^. We took multiple *K*-mers, for the analysis. The *K*-mer frequency was quickly counted by Jellyfish- (v2.2.6) software (http://www.cbcb.umd.edu/software/jellyfish)^72^ and genome size, heterozygosity, repeat sequences, and sequence depth were evaluated using GenomeScope2 software (https://github.com/tbenavi1/genomescope2.0)^20^.

### PacBio HiFi sequencing

For the yellow pitaya (*S. megalanthus*) genome, genomic DNA was extracted from cladodes (plant photosynthetic branches/flattened leaves like a stem). The initial sample was extracted from DNA, which was fragmented using the Megaruptor system to obtain an average size of 15- 20K. We prepared the hairpin SMRTbell-shaped structure after ligating the adapter with DNA and removing the linear and damaged DNA. The BluePippin was used to select the DNA fragments with the required target size. Qualified libraries were sequenced using the BGI Sequel II system (PacBio platform) in HiFi mode.

### Chromosome conformation capture (Hi-C) library construction and sequencing

Hi-C libraries of the diploid (*S. undatus*) and polyploid (*S. megalanthus*) pitaya were constructed by following previously published protocols^73,74^ to identify the genome-wide chromatin interactions. Young cladodes were mixed, and formaldehyde solution (1%) was used to cross- link the DNA of the spatial adjacent sites by following the protocols of Wuhan-Fraser Gene Information Co., Ltd. After cell lysis, the cross-linked chromatin adjacent sites were digested using the specific restriction endonuclease enzyme DpnII. The sticky ends of nucleic acids generated by DpnII were labeled with biotin-dCTP covalently, resulting in produced blunt-ended repaired DNA strands. DNA was physically fragmented and biotin-dCTP labeled fragments were purified using streptavidin beads. T4 DNA polymerase was used to remove the terminal alkaloid tag of the unlinked fragments. The purified product was amplified and sequencing libraries were constructed after inspecting the quality with Qubit 2.0 (identify the library concentration) and Agilent 2100 (detect the integrity of the DNA fragments and insert size) to ensure the output of high-quality Hi-C library. Hi-C libraries were sequenced with an Illumina Hiseq (BGI PE150) platform. The raw reads generated from Hi-C were filtered and trimmed using the trim-galore (v.0.6.7) (https://www.bioinformatics.babraham.ac.uk/projects/trim_galore/). The Hi-C reads of diploid pitaya (*S. undatus*) were aligned with the reference genome^26,75^ and yellow pitaya Hi-C reads were aligned with the newly assembled reference genome of polyploid yellow pitaya (*S. megalanthus*) using the Burrows-Wheeler alignment algorithm (BWA)^76^. To ensure the quality of Hi-C sequenced data, HiCpro alignment (v.2.8.0) was carried out to obtain the read pairs that were uniquely aligned at both ends and obtained the final read pairs ^77^.

### ATAC-sequencing

An assay for Transposase-Accessible Chromatin with high throughput sequencing (ATAC-seq) was carried out for both species of pitaya including diploid (*S. undatus*) and polyploid (*S. megalanthus*) to analyze the chromatin regions accessible to Tn5 transposase^78^. The sucrose precipitation method was used for the ATAC-seq experiment^79^. Young cladode tissues from diploid and polyploid pitaya species were prepared for cell suspension to obtain the nucleus. Tn5 transposase cleaves only open chromatin and simultaneously adds sequencing adapters to both ends of the cleaved DNA fragments. The digested DNA fragments were recovered, amplified by PCR, and libraries size was selected by a gel-free double-sided size-selection protocol using the Agencourt AMPure XP beads (Cat. 63881) at 0.5X and 1.2X. Libraries were sequenced using next-generation high-throughput sequencing (Illumina PE150). The raw data were filtered using trimmomatic software^71^. Clean reads of diploid pitaya were aligned with the reference genome (http://www.pitayagenomic.com/download.php)^26,75^ and clean reads of polyploid pitaya were aligned with the reference genome (newly assembled genome of *Selenicereus megalanthus*) using Bowtie 2 (v 2.2.6). By comparing the signal values of both species, DAR analysis was carried out by R-package DiffBind with DESeq2 and set the filtering threshold to log2FoldChange >0.58 and FDR <0.05. For downstream analysis, we followed all the data processing protocols mentioned by Zhang and colleagues (2021)^80^.

### Transcriptome Sequencing

Young-developing cladode (30 days old stem), fruit pericarp (when the fruit turns its color), and pulp (before maturity) were sampled from both pitaya species (*S. undatus* and *S. megalanthus*) for transcriptome sequencing. Total RNA was extracted from the stem, pericarp, and fruit pulp samples. The concentration and purity of the RNA were detected by Nanodrop 2000, and the RIN value was determined by Agilent 2100. Nanodrop was used to detect the purity (OD260/280), concentration, and normal nucleic acid absorption peak. The total 2 µg RNA with a concentration of ≥ 300 ng/μL, the OD260/280 was between 1.8∼2.2. Raw reads were generated using the BGI Illumina platform.

### Assembly quality assessment

Genome size was estimated using a DNB-sequence survey at 21-mer frequency distribution based on 194.17 Gb cleaned BGI short reads. High-quality genome assembly, and high fidelity (HiFi) reads were used to assemble the *S. megalanthus* genome. To assess the quality of the yellow pitaya reference genome, contiguity, completeness, and correctness (3-Cs) were evaluated. Contiguity was measured as contig N50, the longest contig that comprised 50% of the total genome length. These contigs were scaffolded onto chromosomes using Hi-C reads. Completeness (Integrity and accuracy of the assemblies) was measured using Benchmarking Universal Single-Copy Orthologs (BUSCO)^81^ with a more than 95% score.

### Reference genome assembly and chromosome construction

For the yellow pitaya genome assembly, genomic DNA was sequenced using the BGI PacBio platform (Sequel II system). SMRT bell library was constructed and sequenced. Guppy basecaller (ver 4.1.1)^82^ was employed for calling the reads using the high-accuracy base calling model and the resulting fastq files were used to generate genome assembly (Total length 3,754,928,053 bp) (**Supplementary Table 31**). Clean PacBio subreads were polished using LoRDEC (v.0.7)^83^ with yellow pitaya short reads and scaffolded with Hi-C to anchor the assembled contigs into pseudochromosome molecules. In addition, we used the Purge_Haplotigs software to initialize the genome assembly, identified redundant heterozygous contigs, and removed read depth, distribution, and sequence similarity^22^. The accuracy of the long-read sequencing was also maintained by the depth of the sequencing. The N50 parameters were computed for high-quality genome assembly. Redundant heterozygous contig was identified using the purge-haplotigs^22^ and removed based on the depth distribution of reads and sequence similarity. The BWA software and Minimap2 were used to align the short-read and long-read data of the genome. We calculated the genome mapping rate as 96.96% with the coverage rate of 99.81%. After the genome assembly, genome annotation was carried out through different bioinformatics tools to define sequence elements by *de* novo^84^, homology, sequence location, and function. Repeat sequences including the tandem repeat (microsatellites)^85^ and interspersed repeats (Transposon elements i:e retrotransposons and DNA transposons) were predicted using the TRF software, PILER (http://www.drive5.com/piler)^86^ repeat database (Repbase - GIRI (girinst.org)^87^, repeat modeler software (https://www.repeatmasker.org/) and LTR finder^88^. To predict the structural coding genes, various tools were used including the identification of homologs of closely related species and *de-novo* prediction by GlimmerHMM^89^. Transcriptome data from three tissues including the stem, fruit pericarp, and fruit pulp was aligned using the HISAT2^90^, and transcripts were predicted using the String-Tie^91^. ISOseq3 was used for PacBio full-length transcript acquisition and Transdecoder (https://github.com/TransDecoder/TransDecoder) was subjected to coding region prediction. Using the MAKER2, the predicted gene set was integrated into a non-redundant and complete gene set^92^. Functional annotation of proteins in the genes set was mainly carried by protein databases (SWISS-PROT^93^, InterProScan^94^, Gene Ontology^95^, KEGG^96^, and KOBAS-i^97^) to identify the gene function, metabolic pathway, and structural domains.

### *S. megalanthus* transposable elements and gene annotation

From the assembled genome, nonredundant protein-coding genes were functionally annotated using the MAKER^98^ (v.2.31.11) pipeline by combining *de novo*, homologous searches, and transcriptome data. The transposable elements were predicted *de novo* method TE annotator (EDTA) (1.9.6)^99^. TE library derived from EDTA, *de novo* transcriptome assembly of 3 tissues (stem, pericarp, and pulp), and gene predictions were generated by the fast and accurate algorithm of AUGUSTUS (v.3.4.0)^100^ and GeneMark-EP^+^(v.4.6.3) (http://topaz.gatech.edu/GeneMark/license_download.cgi)^101^. The final new yellow pitaya transcripts were merged with MAKER gene models to produce the final set of transcripts, by combining all the data sets and eliminating the transposable elements related domains, and redundant elements^98^. *Selenicereus undatus* (http://www.pitayagenomic.com/download.php)^3,26,75^, *Carnegiea gigantea* (GCA_029747015.1_UA_SGP5p_2), *Arabidopsis thaliana* (GCA_000001735.1_TAIR10)^102,103^, *Oryza sativa* (GCF_001433935.1_IRGSP-1.0)^104^, *Solanum lycopersicum* (GCF_000188115.5_SL3.1), and *Solanum tuberosum* (GCF_000226075.1_SolTub_3.0)^105^ species were used for homology prediction in yellow pitaya (**Supplementary Table 32**). Functional annotation of proteins in the genes set was mainly carried by protein databases (SWISS-PROT^93^, InterProScan^94^, Gene Ontology^95^, KEGG^96^, and KOBAS-i^97^) to identify the gene function with the cut-off value of 1e^-5^, to obtain the protein function, metabolic pathway, and structural domains.

### Hi-C heat maps of red (diploid) and yellow (polyploid) pitaya

Clean data of polyploid yellow pitaya was aligned with the newly constructed reference genome. For diploid white flesh pitaya, Hi-C clean data was aligned with the reference genome available at http://www.pitayagenomic.com/download.php (accessed on December 13, 2023) using the Juicer software^28^. Messy and noisy reads including single-ended, low-quality, and unaligned reads were filtered. For Hi-C analysis, valid pairs reflect the interaction information between loci on the reference genome and invalid interaction pairs, including self-circle, dangling ends, and PCR duplicates were discarded. The quality of the Hi-C libraries was assessed by indicating the proportion of valid pairs in non-PCR replicates. After comparison and filtering, the Valid Pairs (Q30) obtained were used for subsequent construction of the interaction matrix and downstream analysis, and the comparison statistics of each sample (**Supplementary Table 33)**. 3D conformation of the genome was constructed by aligning the pairwise reads at different loci of the reference genome, requiring a large number of paired-end reads to fully characterize all interactions throughout the genome. The correlation of Hi-C data between parallel samples was analyzed using the GenomeDISCO software^27^. To build the interaction matrix, the reference genome was divided into the size of the bins, then assigned to the filtered valid pairs (Q30), and the double-ended reads were compared to two bins. Due to the uneven distribution of GC content and enzyme cleavage sites on the reference genome, and uneven length of digested fragments, the original interaction matrix was corrected using the Juicer software^28^. A 100-kb resolution genome-wide interaction matrix was constructed, and a corrected interaction matrix was displayed in the form of a heatmap. In addition, a single chromosome interaction matrix with a resolution of 40-kb was constructed for diploid and polyploid pitaya species.

Interaction differences between the diploid and polyploid genomes of pitaya were demonstrated. Interaction matrices of diploid and polyploid pitaya were converted into Z-score matrices^106^. Z- score matrices of both genomes were subtracted, and the final interaction matrices were displayed as a heatmap with 200 kb resolution. The interaction frequency and distance were computed by comparing both genomes at 200 kb resolution^42,43^.

### Compartment identification in both pitaya species

Genome compartments were investigated to find the potential link between the regions within a single chromosome after constructing an interaction map^30^. According to the single chromosome interaction matrix at 100-kb resolution, the PC1 value of bin was obtained by the C-score method of bin^107^, and all the bins on the reference genome of polyploid yellow and diploid red pitaya were divided into two regions including active regions (compartment A) and inactive regions (compartment B). The C-score method was used to analyze the genomic compartments and the positive and negative bins were referred to as compartment A and B to compute the distribution of compartments on each chromosome. The bins of the consecutive compartment A or compartment B were combined to calculate the number and length distribution of compartment A/B in the whole genome. The length distribution of the compartment was displayed in the form of the boxplot. We computed the significance of the length distribution of Compartment A and Compartment B after merging and obtained the p-value by Wilcoxon’s rank-sum test^108^. The number of genes in each bin in the whole genome of compartment A and compartment B of the sample was counted. The distribution of gene numbers in compartments were displayed in the form of box plots, and the significance of the distribution of genes between compartment A and compartment B was tested. To identify the association of GC content of genomic sequence with compartments (A/B) structure^29,109^, GC content in each bin was counted and the GC content distribution was displayed in the form of box plot and test for significance by Wilcoxon test^108^.

### TAD structural analysis, classification of genes, and motif analysis in red and yellow pitaya species

Topologically associating domains (TADs) boundaries were calculated using the insulation score method using cworld-dekker^106^ which reflects the interaction between the two sides of each bin. The interaction isolation between the two sides of the bin was recognized as the TAD boundary. TAD boundary was obtained at 40-kb resolution on each chromosome of diploid and polyploid species of pitaya. All bins in the whole genome were divided into two categories; those bins located at the TAD boundary (Border) and those located inside the TAD (inter), and the number of genes in each bin in the TAD boundary and TAD of the sample was counted, and the distribution of the gene number between the TAD boundary and inside the TAD was displayed in the form of box plot and significance of the gene number was tested using Wilcoxon test^108^. To study the transcription factors and structural proteins, motif analysis was carried out using JASPAR database (http://jaspar.genereg.net/) and MEME Suite software (http://meme-suite.org/) from the whole genome of diploid and polyploid pitaya. The number of TAD boundaries with the motif was counted and the top 5 motifs were displayed.

### Gene function annotation in differential compartments region

The gene sets located in the compartment region were functionally annotated and the 20 most significant enrichments were selected and shown on GO and KEGG scatter plots.

### Analysis of significant interaction sites in diploid and polyploid pitaya species

At 10-kb resolution, the Fit-Hi-C software (v1.0.1) (L resolution) was used to analyze the significant interactions between the two loci within the genome of pitaya species^39^. The significant cis and trans interaction sites were obtained based on the number of reads supporting the interaction by p-value ≤ 0.01 and q-value ≤ 0.01. The corresponding cumulative probability and interacting loci were sorted. If the *P* value and *q* value were <0.01 and the read count was >2 were identified as significant interactions^110^.

### Comparative genome structural variation and genome evolution analysis

Genome alignment of reference and query genomes was performed using the Nucmer embedded in the MUMmer4 toolkit with parameters of“-c 40 -l 90”^111^. The alignments were filtered by running the Delta-filter with parameters of “ -1 ”. The Delta alignment output was parsed using Show-coords with parameters of “ -THrd ”. Then, SNPs and InDels (1-50 bp) were identified by running Show-SNPs (-ClrTH).SyRI(v.1.6.3) software was used to identify Inversion, Translocation, CNV, and PAV^112^. All variants were annotated using ANNOVAR software^113^. For yellow pitaya genome evolutionary analysis, orthologous gene families of *S. megalanthus*, closely related cactus species [*S. undatus-*(http://www.pitayagenomic.com/download.php), (*Carnegiea gigantea*- https://www.ncbi.nlm.nih.gov/datasets/genome/GCA_029747015.1/, GCA_029747015.1_UA_SGP5p_2)] and plant species including *Beta vulgaris* (https://phytozome-next.jgi.doe.gov/info/Bvulgaris_EL10_1_0, Bvulgaris_548_EL10_1.0), *Spinacia oleracea* (https://phytozome-next.jgi.doe.gov/info/Soleracea_Spov3, Soleracea_575_Spov3), *Sedum album* (https://genomevolution.org/CoGe/SearchResults.pl?s=Sedum%20album&p=genome, V3), *Arabidopsis thaliana*(https://phytozome-next.jgi.doe.gov/info/Athaliana_TAIR10, Athaliana_167_TAIR10), *Kalanchoe fedtschenkoi* (https://phytozome-next.jgi.doe.gov/info/Kfedtschenkoi_v1_1, Kfedtschenkoi_382_v1.1), *Oryza sativa* (https://phytozome-next.jgi.doe.gov/info/Osativa_v7_0, Osativa_323_v7.0), *Cannabis sativa* (https://www.ncbi.nlm.nih.gov/datasets/genome/GCF_029168945.1/, GCF_029168945.1_ASM2916894v1) were obtained using the OrthoMCL (V1.4, http://orthomcl. org/orthomcl/). A super alignment matrix was obtained by aligning the sin gle-copy families using the MUSCLE (v.3.8.31) and following the pipeline reported in literature^114–116^.

Synonymous substitution rate (*Ks*) distribution for paralogs was identified from autotetraploid yellow pitaya (*S. megalanthus*) using the 11 chromosomes and orthologs were identified from pitaya species (*S. undatus* and *S. megalanthus*) using the MCScanX software (http://chibba.pgml.uga.edu/mcscan2/). By distinguishing the synonymous substitution (*Ks*) and nonsynonymous substitution (*Ka*) rate, we measure the selection pressure at protein level which is generally equal to 1 (Ka/Ks=1). We used WGDI software to calculate the *Ks* values, filtered the data based on collinear blocks (reduce the error caused by tandem repeats) and draw the distribution of *Ks* frequency to observe the number of peaks to observe the WGD events.

We selected single-copy gene families to align using MUSCLE (v3.8.31, http://www.drive5.com/muscle/). To identify the single base difference, we use vcf2dis software for each gene family, and the nucleic acid difference rate was calculated 0.010014.

### Linking Hi-C data with RNA-Seq

To identify whether subcompartment switching is correlated with changes in expression level, we classified all 40-kb genomic bins into three groups based on changes in the subcompartment of pitaya pericarp tissues. To link the Hi-C data with RNA-seq, we followed the previous studies^70,53^ and approaches to explore the reorganization of chromatin domains (TADs) and loops is correlated with changes in expression level. We correlated this classification of TAD features and loops with the changes in gene expression profiles, including DEGs and fold changes between tissues.

## Acknowledgments

This study was funded by the Hainan Province Science and Technology Special Fund (ZDYF2022XDNY190), the Project of Sanya Yazhou Bay Science and Technology City (Grant Number: SCKJ-JYRC-2022-83), Innovational Fund for Scientific and Technological Personnel of Hainan Province, Collaborative Innovation Center of Nanfan and High-Efficiency Tropical Agriculture, Hainan University Funding (XTCX2022NYB09) and the Hainan Provincial Natural Science Foundation of China (421RC486). . It was also supported by the Food Futures Institute of Murdoch University to RKV.

## Contributions

Q.U.Z, M.F.N., A.R., H.-F.Q., and R.K.V. designed the research. Q.U.Z., M.G. M.F.N. M.I., D.K., L.H., A.A.K., S.K., B.U. analyzed the data. L.W., G.W., X.Y., S.Z., Z.Y., P.L., Z.X.Z. generated plant materials. H.-F.W., R.K.V. supervised the whole-genome sequencing work. Q.U.Z. wrote the article with input from the other authors.

## Ethics declarations

Competing interests

The authors declare no competing interests.

## Data availability

All raw sequence data of PacBio, ATAC, and RNA-Seq have been deposited in the National Center for Biotechnology Information-NCBI (SRA database under BioProject accession <u>PRJNA1117350</u>) and Hi-C data are available at the China National Center for Bioinformation-CNCB (Under GSA database Bioproject ID <u>PRJCA026647</u>). Reference genome assembly and annotation files of *Selenicereus megalanthus-Sme* (Sme.fasta.gz, Sme gene annotation.xls, Sme annotation_gene_transcript.xls, Sme.gene.cds, Sme.gene.gff and Sme.gene.pep.) are available at https://figshare.com/s/6f98e7b31f4af6bc3f62.

## Supplementary information

Supplementary Figures (1-41) and Supplementary Tables (1-33) are available as attached files.

## References

1 Hershkovitz, M. A. & Zimmer, E. A. J. T. On the evolutionary origins of the cacti. 46, 217–232 (1997).

2 Khan, D. et al. The evolutionary history and distribution of cactus germplasm resources, as well as potential domestication under a changing climate. Journal of Systematics and Evolution n/a 10.1111/jse.13042

3 Zheng, J., Meinhardt, L. W., Goenaga, R., Zhang, D. & Yin, Y. The chromosome-level genome of dragon fruit reveals whole-genome duplication and chromosomal co-localization of betacyanin biosynthetic genes. Horticulture Research 8, 63 (2021). 10.1038/s41438-021-00501-6

4 Tel-Zur, N. Breeding an underutilized fruit crop: a long-term program for Hylocereus. Horticulture Research 9 (2022). 10.1093/hr/uhac078

5 Tenore, G. C., Novellino, E. & Basile, A. Nutraceutical potential and antioxidant benefits of red pitaya (Hylocereus polyrhizus) extracts. Journal of Functional Foods 4, 129–136 (2012). 10.1016/j.jff.2011.09.003

6 Jiang, H. et al. Nutrition, phytochemical profile, bioactivities and applications in food industry of pitaya (Hylocereus spp.) peels: A comprehensive review. Trends in Food Science & Technology 116, 199–217 (2021). 10.1016/j.tifs.2021.06.040

7 Abirami, K. et al. Distinguishing three Dragon fruit (Hylocereus spp.) species grown in Andaman and Nicobar Islands of India using morphological, biochemical and molecular traits. Scientific Reports 11, 2894 (2021). 10.1038/s41598-021-81682-x

8 Wu, L.-c., et al. Antioxidant and antiproliferative activities of red pitaya. Food Chemistry 95, 319–327 (2006). 10.1016/j.foodchem.2005.01.002

9 Paśko, P. et al. Dragon Fruits as a Reservoir of Natural Polyphenolics with Chemopreventive Properties. Molecules 26 (2021). 10.3390/molecules26082158

10 Watanabe, K. in Crop Production under Stressful Conditions: Application of Cutting-edge Science and Technology in Developing Countries (eds Makie Kokubun & Shuichi Asanuma) 177-193 (Springer Singapore, 2018).

11 Morillo-Coronado, A. C., Manjarres-Hernández, E. H., Saenz-Quintero, Ó. J. & Morillo-Coronado, Y. Morphoagronomic Evaluation of Yellow Pitahaya (Selenicereus megalanthus Haw.) in Miraflores, Colombia. 12, 1582 (2022).

12 Gandía-Herrero, F. & García-Carmona, F. J. N. P. The dawn of betalains. 227, 664–666 (2020).

13 Arakaki, M. et al. Contemporaneous and recent radiations of the world’s major succulent plant lineages. 108, 8379–8384 (2011).

14 Britton, N. L. & Rose, J. N. The Cactaceae: descriptions and illustrations of plants of the cactus family. Vol. 3 (Courier Corporation, 1963).

15 Moran, R. J. G. H. Selenicereus megalanthus (Schumann) Moran. 8, 325 (1953).

16 Bauer, R. A synopsis of the tribe Hylocereeae F. Buxb. (Hunt, 2003).

17 Plume, O. et al. Testing a hypothesis of intergeneric allopolyploidy in vine cacti (Cactaceae: Hylocereeae). 38, 737–751 (2013).

18 Korotkova, N., Borsch, T. & Arias, S. J. P. A phylogenetic framework for the Hylocereeae (Cactaceae) and implications for the circumscription of the genera. 327, 1–46 (2017).

19 Dixon, J. R. et al. Topological domains in mammalian genomes identified by analysis of chromatin interactions. Nature 485, 376–380 (2012). 10.1038/nature11082

20 Ranallo-Benavidez, T. R., Jaron, K. S. & Schatz, M. C. GenomeScope 2.0 and Smudgeplot for reference-free profiling of polyploid genomes. Nature Communications 11, 1432 (2020). 10.1038/s41467-020-14998-3

21 Ma, L. et al. Diploid and tetraploid genomes of Acorus and the evolution of monocots. Nature Communications 14, 3661 (2023). 10.1038/s41467-023-38829-3

22 Roach, M. J., Schmidt, S. A. & Borneman, A. R. Purge Haplotigs: allelic contig reassignment for third-gen diploid genome assemblies. BMC Bioinformatics 19, 460 (2018). 10.1186/s12859-018-2485-7

23 Huang, N. & Li, H. compleasm: a faster and more accurate reimplementation of BUSCO. Bioinformatics 39 (2023). 10.1093/bioinformatics/btad595

24 Van Drunen, W. E. & Husband, B. C. Evolutionary associations between polyploidy, clonal reproduction, and perenniality in the angiosperms. New Phytologist 224, 1266–1277 (2019). 10.1111/nph.15999

25 Otto, S. P. The Evolutionary Consequences of Polyploidy. Cell 131, 452–462 (2007). 10.1016/j.cell.2007.10.022

26 Chen, J.-y., et al. A chromosome-scale genome sequence of pitaya (Hylocereus undatus) provides novel insights into the genome evolution and regulation of betalain biosynthesis. Horticulture Research 8, 164 (2021). 10.1038/s41438-021-00612-0

27 Ursu, O. et al. GenomeDISCO: a concordance score for chromosome conformation capture experiments using random walks on contact map graphs. Bioinformatics 34, 2701–2707 (2018). 10.1093/bioinformatics/bty164

28 Durand, N. C. et al. Juicer Provides a One-Click System for Analyzing Loop-Resolution Hi- C Experiments. Cell Systems 3, 95–98 (2016). 10.1016/j.cels.2016.07.002

29 Rao, S. S. et al. A 3D map of the human genome at kilobase resolution reveals principles of chromatin looping. Cell 159, 1665–1680 (2014). 10.1016/j.cell.2014.11.021

30 Lieberman-Aiden, E. et al. Comprehensive mapping of long-range interactions reveals folding principles of the human genome. Science 326, 289–293 (2009). 10.1126/science.1181369

31 Zheng, X. & Zheng, Y. CscoreTool: fast Hi-C compartment analysis at high resolution. Bioinformatics 34, 1568–1570 (2017). 10.1093/bioinformatics/btx802

32 Crane, E. et al. Condensin-driven remodelling of X chromosome topology during dosage compensation. Nature 523, 240–244 (2015). 10.1038/nature14450

33 Alipour, E. & Marko, J. F. Self-organization of domain structures by DNA-loop-extruding enzymes. Nucleic Acids Res 40, 11202–11212 (2012). 10.1093/nar/gks925

34 Szabo, Q., Bantignies, F. & Cavalli, G. Principles of genome folding into topologically associating domains. 5, eaaw1668 (2019). doi:10.1126/sciadv.aaw1668

35 Tourdot, E. & Grob, S. Three-dimensional chromatin architecture in plants – General features and novelties. European Journal of Cell Biology 102, 151344 (2023). 10.1016/j.ejcb.2023.151344

36 Doğan, E. S. & Liu, C. Three-dimensional chromatin packing and positioning of plant genomes. Nat Plants 4, 521–529 (2018). 10.1038/s41477-018-0199-5

37 Li, E. et al. Long-range interactions between proximal and distal regulatory regions in maize. 10, 2633 (2019).

38 Dong, P. et al. 3D Chromatin Architecture of Large Plant Genomes Determined by Local A/B Compartments. Mol Plant 10, 1497–1509 (2017). 10.1016/j.molp.2017.11.005

39 Ay, F., Bailey, T. L. & Noble, W. S. Statistical confidence estimation for Hi-C data reveals regulatory chromatin contacts. Genome Res 24, 999–1011 (2014). 10.1101/gr.160374.113

40 Pei, L., Li, G., Lindsey, K., Zhang, X. & Wang, M. Plant 3D genomics: the exploration and application of chromatin organization. 230, 1772–1786 (2021). 10.1111/nph.17262

41 Liu, C., Cheng, Y.-J., Wang, J.-W. & Weigel, D. Prominent topologically associated domains differentiate global chromatin packing in rice from Arabidopsis. Nature Plants 3, 742–748 (2017). 10.1038/s41477-017-0005-9

42 Du, Z. et al. Allelic reprogramming of 3D chromatin architecture during early mammalian development. Nature 547, 232–235 (2017). 10.1038/nature23263

43 Ke, Y. et al. 3D Chromatin Structures of Mature Gametes and Structural Reprogramming during Mammalian Embryogenesis. Cell 170, 367–381.e320 (2017). 10.1016/j.cell.2017.06.029

44 Wang, M. et al. Evolutionary dynamics of 3D genome architecture following polyploidization in cotton. Nature Plants 4, 90–97 (2018). 10.1038/s41477-017-0096-3

45 Schiessl, S.-V., Katche, E., Ihien, E., Chawla, H. S. & Mason, A. S. The role of genomic structural variation in the genetic improvement of polyploid crops. The Crop Journal 7, 127–140 (2019). 10.1016/j.cj.2018.07.006

46 Parisod, C., Holderegger, R. & Brochmann, C. Evolutionary consequences of autopolyploidy. New Phytologist 186, 5–17 (2010). 10.1111/j.1469-8137.2009.03142.x

47 Raina, S. N. et al. Associated chromosomal DNA changes in polyploids. Genome 37, 560–564 (1994). 10.1139/g94-080

48 Ozkan, H., Tuna, M. & Galbraith, D. No DNA loss in autotetraploids of Arabidopsis thaliana. Plant Breeding 125, 288–291 (2006).

49 Bhadra, S., Leitch, I. J. & Onstein, R. E. From genome size to trait evolution during angiosperm radiation. Trends in Genetics 39, 728–735 (2023). 10.1016/j.tig.2023.07.006

50 Zhang, L. et al. 3D genome structural variations play important roles in regulating seed oil content of Brassica napus. Plant Communications 5, 100666 (2024). 10.1016/j.xplc.2023.100666

51 Tannenbaum, M. et al. Regulatory chromatin landscape in Arabidopsis thaliana roots uncovered by coupling INTACT and ATAC-seq. Plant Methods 14, 113 (2018). 10.1186/s13007-018-0381-9

52 Robinson, J. T., et al. Integrative genomics viewer. Nature Biotechnology 29, 24–26 (2011). 10.1038/nbt.1754

53 Liu, Z. et al. Linking genome structures to functions by simultaneous single-cell Hi-C and RNA-seq. Science 380, 1070–1076 (2023). 10.1126/science.adg3797

54 Lopez-Nieves, S. et al. Relaxation of tyrosine pathway regulation underlies the evolution of betalain pigmentation in Caryophyllales. New Phytologist 217, 896–908 (2018). 10.1111/nph.14822

55 Timoneda, A. et al. The evolution of betalain biosynthesis in Caryophyllales. New Phytol 224, 71–85 (2019). 10.1111/nph.15980

56 Schliemann, W., Kobayashi, N. & Strack, D. The Decisive Step in Betaxanthin Biosynthesis Is a Spontaneous Reaction1. Plant Physiology 119, 1217–1232 (1999). 10.1104/pp.119.4.1217

57 De Storme, N., Copenhaver, G. P. & Geelen, D. Production of Diploid Male Gametes in Arabidopsis by Cold-Induced Destabilization of Postmeiotic Radial Microtubule Arrays Plant Physiology 160, 1808–1826 (2012). 10.1104/pp.112.208611 %J Plant Physiology

58 Mason, A. S., Nelson, M. N., Yan, G. & Cowling, W. A. Production of viable male unreduced gametes in Brassica interspecific hybrids is genotype specific and stimulated by cold temperatures. BMC Plant Biology 11, 103 (2011). 10.1186/1471-2229-11-103

59 Pécrix, Y. et al. Polyploidization mechanisms: temperature environment can induce diploid gamete formation in Rosa sp. Journal of Experimental Botany 62, 3587–3597 (2011). 10.1093/jxb/err052 %J Journal of Experimental Botany

60 Mable, B. K., Alexandrou, M. A. & Taylor, M. I. Genome duplication in amphibians and fish: an extended synthesis. 284, 151–182 (2011). 10.1111/j.1469-7998.2011.00829.x

61 Gaut, B. S., Miller, A. J. & Seymour, D. K. Living with Two Genomes: Grafting and Its Implications for Plant Genome-to-Genome Interactions, Phenotypic Variation, and Evolution. Annual Review of Genetics 53, 195–215 (2019). 10.1146/annurev-genet-112618-043545

62 Dwivedi, S. L. et al. Evolutionary dynamics and adaptive benefits of deleterious mutations in crop gene pools. Trends in Plant Science 28, 685–697 (2023). 10.1016/j.tplants.2023.01.006

63 Stebbins, G. L. Variation and evolution in plants. (Columbia University Press, 1950).

64 Soltis, P. S. & Soltis, D. E. The role of genetic and genomic attributes in the success of polyploids. Proc Natl Acad Sci U S A 97, 7051–7057 (2000). 10.1073/pnas.97.13.7051

65 Lieberman-Aiden, E. et al. Comprehensive Mapping of Long-Range Interactions Reveals Folding Principles of the Human Genome. 326, 289–293 (2009). doi:10.1126/science.1181369

66 Cremer, T. & Cremer, M. J. C. S. H. p. i. b. Chromosome territories. 2, a003889 (2010).

67 Ouyang, W., Xiong, D., Li, G. & Li, X. Unraveling the 3D Genome Architecture in Plants: Present and Future. Molecular Plant 13, 1676–1693 (2020). 10.1016/j.molp.2020.10.002

68 Crevillén, P., Sonmez, C., Wu, Z. & Dean, C. A gene loop containing the floral repressor FLC is disrupted in the early phase of vernalization. Embo j 32, 140–148 (2013). 10.1038/emboj.2012.324

69 Karaaslan, E. S. et al. Marchantia TCP transcription factor activity correlates with three- dimensional chromatin structure. Nature Plants 6, 1250–1261 (2020). 10.1038/s41477-020-00766-0

70 Liao, Y. et al. The 3D architecture of the pepper genome and its relationship to function and evolution. Nature Communications 13, 3479 (2022). 10.1038/s41467-022-31112-x

71 Bolger, A. M., Lohse, M. & Usadel, B. J. B. Trimmomatic: a flexible trimmer for Illumina sequence data. 30, 2114–2120 (2014).

72 Marçais, G. & Kingsford, C. A fast, lock-free approach for efficient parallel counting of occurrences of k-mers. Bioinformatics 27, 764–770 (2011). 10.1093/bioinformatics/btr011

73 Liu, C. In Situ Hi-C Library Preparation for Plants to Study Their Three-Dimensional Chromatin Interactions on a Genome-Wide Scale. Methods Mol Biol 1629, 155–166 (2017). 10.1007/978-1-4939-7125-1_11

74 Burton, J. N. et al. Chromosome-scale scaffolding of de novo genome assemblies based on chromatin interactions. 31, 1119–1125 (2013).

75 Chen, C. et al. Pitaya Genome and Multiomics Database (PGMD): A Comprehensive and Integrative Resource of Selenicereus undatus. Genes (Basel) 13 (2022). 10.3390/genes13050745

76 Li, H. & Durbin, R. J. b. Fast and accurate short read alignment with Burrows–Wheeler transform. 25, 1754–1760 (2009).

77 Servant, N. et al. HiC-Pro: an optimized and flexible pipeline for Hi-C data processing. 16, 1–11 (2015).

78 Buenrostro, J. D., Giresi, P. G., Zaba, L. C., Chang, H. Y. & Greenleaf, W. J. Transposition of native chromatin for fast and sensitive epigenomic profiling of open chromatin, DNA- binding proteins and nucleosome position. Nature Methods 10, 1213–1218 (2013). 10.1038/nmeth.2688

79 Tannenbaum, M. et al. Regulatory chromatin landscape in Arabidopsis thaliana roots uncovered by coupling INTACT and ATAC-seq. Plant Methods 14, 113 (2018). 10.1186/s13007-018-0381-9

80 Zhang, X. et al. Chromatin spatial organization of wild type and mutant peanuts reveals high-resolution genomic architecture and interaction alterations. Genome Biology 22, 315 (2021). 10.1186/s13059-021-02520-x

81 Simão, F. A., Waterhouse, R. M., Ioannidis, P., Kriventseva, E. V. & Zdobnov, E. M. BUSCO: assessing genome assembly and annotation completeness with single-copy orthologs. Bioinformatics 31, 3210–3212 (2015). 10.1093/bioinformatics/btv351

82 Di Tommaso, P. et al. Nextflow enables reproducible computational workflows. 35, 316–319 (2017).

83 Salmela, L. & Rivals, E. J. B. LoRDEC: accurate and efficient long read error correction. 30, 3506–3514 (2014).

84 Price, A. L., Jones, N. C. & Pevzner, P. A. De novo identification of repeat families in large genomes. Bioinformatics 21 Suppl 1, i351–358 (2005). 10.1093/bioinformatics/bti1018

85 Benson, G. Tandem repeats finder: a program to analyze DNA sequences. Nucleic Acids Res 27, 573–580 (1999). 10.1093/nar/27.2.573

86 Edgar, R. C. & Myers, E. W. PILER: identification and classification of genomic repeats. Bioinformatics 21 Suppl 1, i152–158 (2005). 10.1093/bioinformatics/bti1003

87 Bao, W., Kojima, K. K. & Kohany, O. Repbase Update, a database of repetitive elements in eukaryotic genomes. Mobile DNA 6, 11 (2015). 10.1186/s13100-015-0041-9

88 Xu, Z. & Wang, H. LTR_FINDER: an efficient tool for the prediction of full-length LTR retrotransposons. Nucleic Acids Res 35, W265–268 (2007). 10.1093/nar/gkm286

89 Majoros, W. H., Pertea, M. & Salzberg, S. L. TigrScan and GlimmerHMM: two open source ab initio eukaryotic gene-finders. Bioinformatics 20, 2878–2879 (2004). 10.1093/bioinformatics/bth315

90 Kim, D., Paggi, J. M., Park, C., Bennett, C. & Salzberg, S. L. Graph-based genome alignment and genotyping with HISAT2 and HISAT-genotype. Nature Biotechnology 37, 907–915 (2019). 10.1038/s41587-019-0201-4

91 Pertea, M. et al. StringTie enables improved reconstruction of a transcriptome from RNA-seq reads. Nature Biotechnology 33, 290–295 (2015). 10.1038/nbt.3122

92 Holt, C. & Yandell, M. MAKER2: an annotation pipeline and genome-database management tool for second-generation genome projects. BMC Bioinformatics 12, 491 (2011). 10.1186/1471-2105-12-491

93 Bairoch, A. & Apweiler, R. The SWISS-PROT protein sequence database and its supplement TrEMBL in 2000. Nucleic Acids Res 28, 45–48 (2000). 10.1093/nar/28.1.45

94 Zdobnov, E. M. & Apweiler, R. InterProScan – an integration platform for the signature- recognition methods in InterPro. Bioinformatics 17, 847–848 (2001). 10.1093/bioinformatics/17.9.847

95 Ashburner, M. et al. Gene Ontology: tool for the unification of biology. Nature Genetics 25, 25–29 (2000). 10.1038/75556

96 Kanehisa, M. & Goto, S. KEGG: kyoto encyclopedia of genes and genomes. Nucleic Acids Res 28, 27–30 (2000). 10.1093/nar/28.1.27

97 Bu, D. et al. KOBAS-i: intelligent prioritization and exploratory visualization of biological functions for gene enrichment analysis. Nucleic Acids Res 49, W317–w325 (2021). 10.1093/nar/gkab447

98 Li, X. et al. Origin and evolution of the triploid cultivated banana genome. Nature genetics (2023). 10.1038/s41588-023-01589-3

99 Ou, S. et al. Benchmarking transposable element annotation methods for creation of a streamlined, comprehensive pipeline. 20, 1–18 (2019).

100 Stanke, M., Diekhans, M., Baertsch, R. & Haussler, D. Using native and syntenically mapped cDNA alignments to improve de novo gene finding. Bioinformatics 24, 637–644 (2008). 10.1093/bioinformatics/btn013 %J Bioinformatics

101 Brůna, T., Lomsadze, A. & Borodovsky, M. GeneMark-EP+: eukaryotic gene prediction with self-training in the space of genes and proteins. NAR Genomics and Bioinformatics 2 (2020). 10.1093/nargab/lqaa026

102 Swarbreck, D. et al. The Arabidopsis Information Resource (TAIR): gene structure and function annotation. Nucleic Acids Res 36, D1009–1014 (2008). 10.1093/nar/gkm965

103 Sloan, D. B., Wu, Z. & Sharbrough, J. Correction of Persistent Errors in Arabidopsis Reference Mitochondrial Genomes. Plant Cell 30, 525–527 (2018). 10.1105/tpc.18.00024

104 Kawahara, Y. et al. Improvement of the Oryza sativa Nipponbare reference genome using next generation sequence and optical map data. Rice (N Y) 6, 4 (2013). 10.1186/1939-8433-6-4

105 Xu, X. et al. Genome sequence and analysis of the tuber crop potato. Nature 475, 189–195 (2011). 10.1038/nature10158

106 Crane, E. et al. Condensin-driven remodelling of X chromosome topology during dosage compensation. Nature 523, 240–244 (2015). 10.1038/nature14450

107 Zheng, X. & Zheng, Y. CscoreTool: fast Hi-C compartment analysis at high resolution. Bioinformatics 34, 1568–1570 (2018). 10.1093/bioinformatics/btx802

108 Haynes, W. in Encyclopedia of Systems Biology (eds Werner Dubitzky, Olaf Wolkenhauer, Kwang-Hyun Cho, & Hiroki Yokota) 2354-2355 (Springer New York, 2013).

109 Lajoie, B. R., Dekker, J. & Kaplan, N. The Hitchhiker’s guide to Hi-C analysis: practical guidelines. Methods 72, 65–75 (2015). 10.1016/j.ymeth.2014.10.031

110 Zhang, L. et al. 3D genome structural variations play important roles in regulating seed oil content of Brassica napus. Plant Communications, 100666 (2023). 10.1016/j.xplc.2023.100666

111 Marçais, G. et al. MUMmer4: A fast and versatile genome alignment system. PLOS Computational Biology 14, e1005944 (2018). 10.1371/journal.pcbi.1005944

112 Goel, M., Sun, H., Jiao, W.-B. & Schneeberger, K. SyRI: finding genomic rearrangements and local sequence differences from whole-genome assemblies. Genome Biology 20, 277 (2019). 10.1186/s13059-019-1911-0

113 Wang, K., Li, M. & Hakonarson, H. ANNOVAR: functional annotation of genetic variants from high-throughput sequencing data. Nucleic Acids Res 38, e164 (2010). 10.1093/nar/gkq603

114 Liao, N. et al. Chromosome-level genome assembly of bunching onion illuminates genome evolution and flavor formation in Allium crops. Nature Communications 13, 6690 (2022). 10.1038/s41467-022-34491-3

115 Hu, G. et al. Two divergent haplotypes from a highly heterozygous lychee genome suggest independent domestication events for early and late-maturing cultivars. Nature Genetics 54, 73–83 (2022). 10.1038/s41588-021-00971-3

116 Marchant, D. B. et al. Dynamic genome evolution in a model fern. Nat Plants 8, 1038–1051 (2022). 10.1038/s41477-022-01226-7

