## Supplementary Figures 1-41 for "Chromosome-level genome assembly of autotetraploid *Selenicereus megalanthus* and gaining genomic insights into the evolution of trait patterning in diploid and polyploid pitaya species"

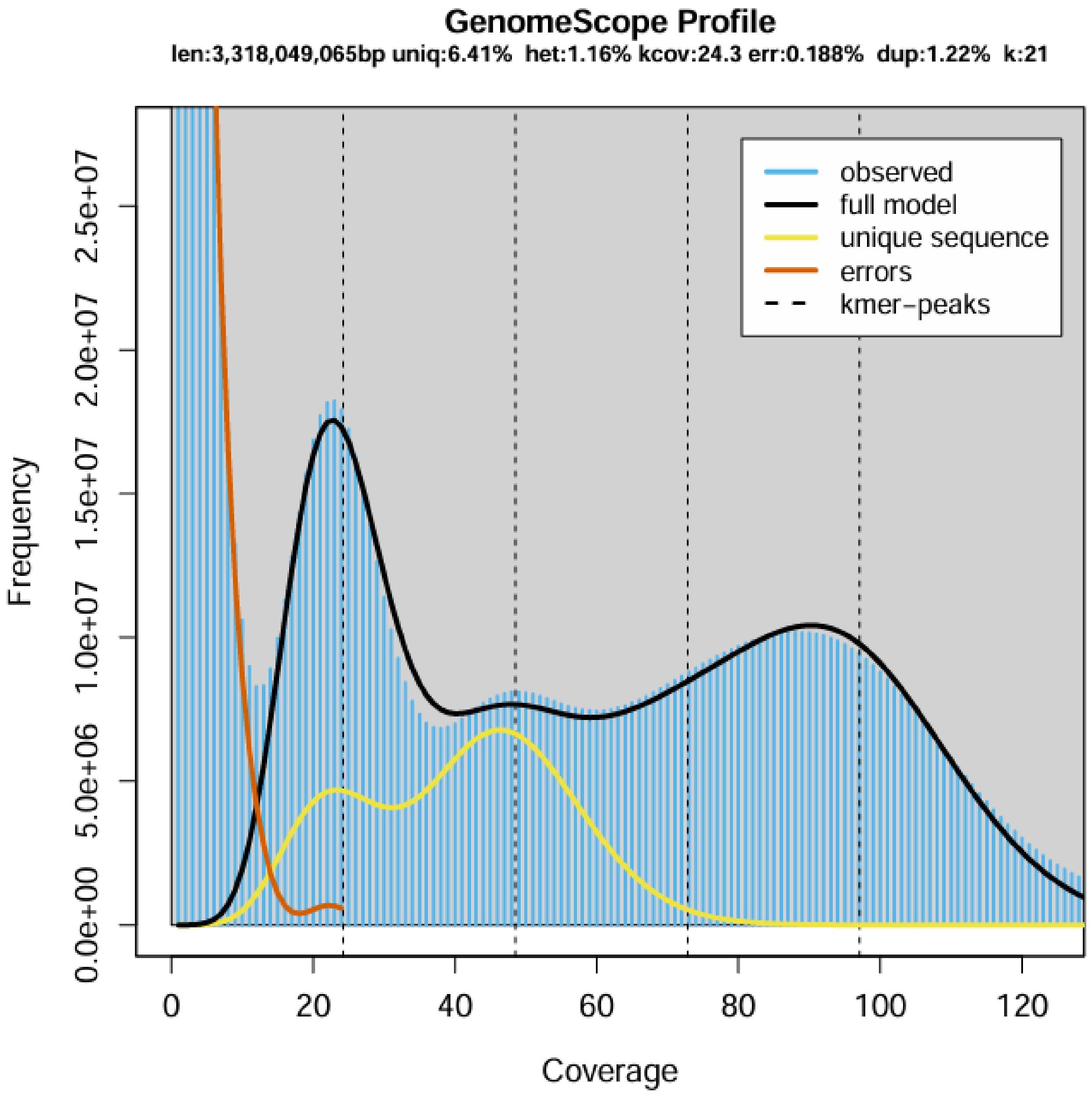


**Supplementary Figure 1**: **21-mer frequency distribution curve of yellow pitaya (*S. megalanthus*).** The genome size of the yellow pitaya was estimated as 4.56Gb as autotetraploid with AAAA K-mer pairs up to 97% and narrowly allopolyploid AAAB/AABB/ABCD *K*-mer pairs up to 3% ratio. The X-axis is the depth, and the Y-axis is the proportion that exhibits the frequency at which depth is divided by the total frequency of all depths.

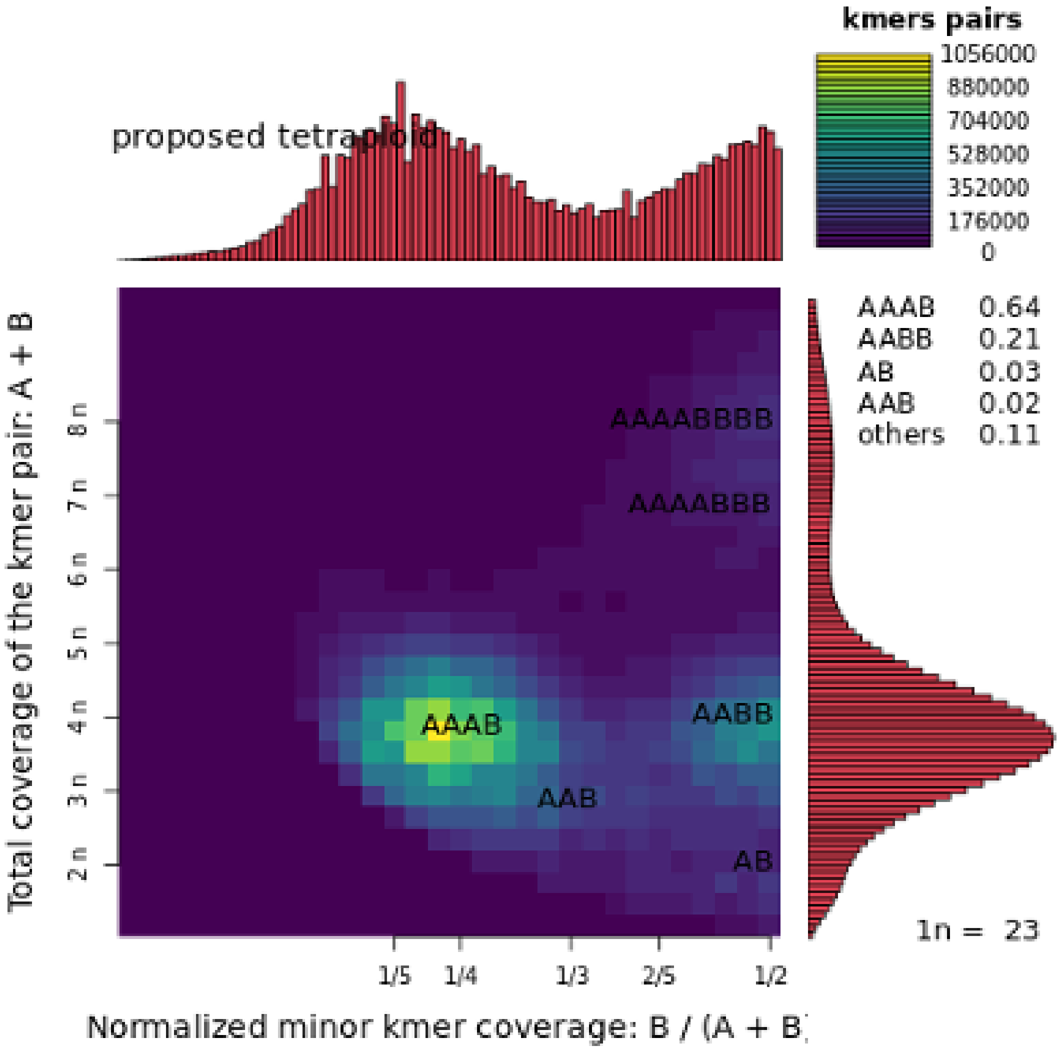


**Supplementary Figure 2: Smudgeplot for the tetraploid yellow pitaya (*Selenicereus megalanthus*).** Two-dimensional heatmaps to predict the ploidy level from clean reads using Smudgeplot. The color intensity represents the approximate amount of the K-mers per bin ranging from purple dark to yellow light. The estimated ploidy level is shown on the Y-Axis of the heatmap and various ploidies on the right side of the heatmap.

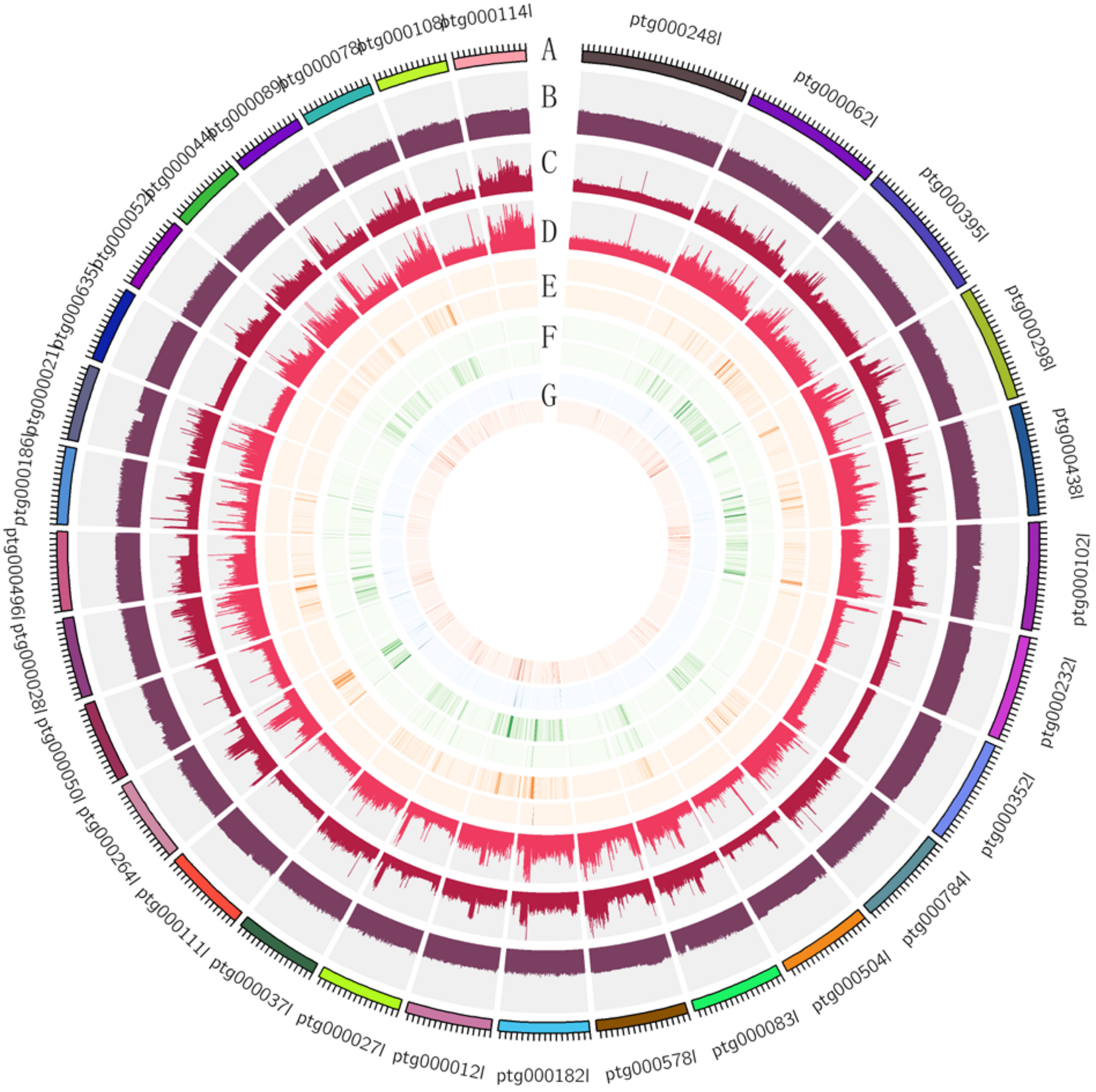


**Supplementary Figure 3: Genome circular map presented** as **A)** genomic information, **B)** distribution of GC content **C)** Short read depth distribution **D)** Long read depth distribution **E)** Outer ring exhibiting homozygous SNPs and inner ring represents heterozygous SNPs **F)** Outer ring exhibiting homozygous InDel and inner ring represents heterozygous InDel **G)** Complete alignment of BUSCO gene distribution on the genome. Single copy BUSCO exhibited in blue color and duplicated BUSCO shown in red color.

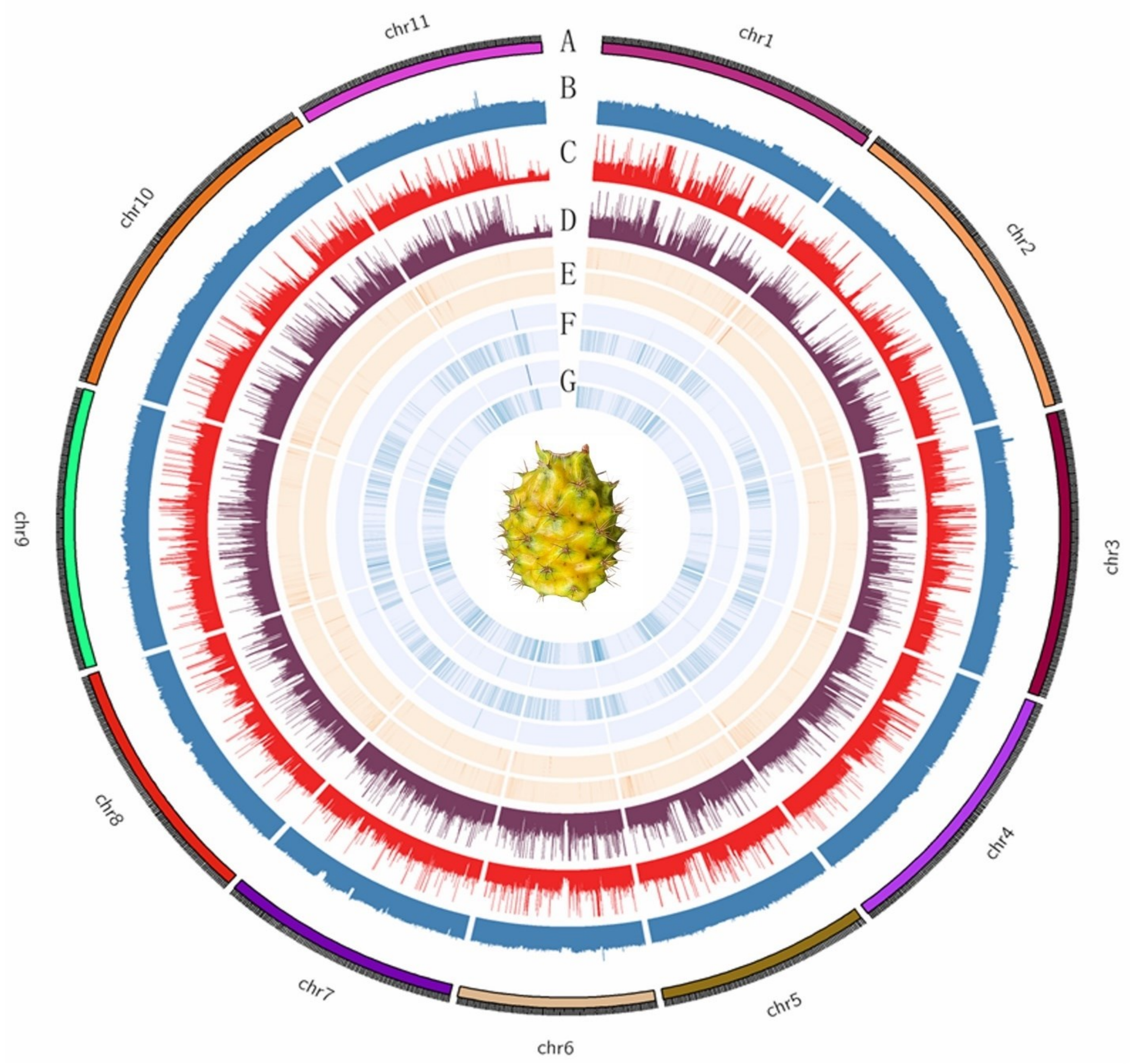


**Supplementary Figure 4: The genomic features of autotetraploid pitaya genome. A)** Genomic information, **B)** GC content distribution, **C)** Short read depth distribution, **D)** Long read depth distribution, **E)** Distribution of BUSCO genes on the yellow pitaya genome was completely aligned with a single copy gene in the outer circle and the duplicated gene in the inner circle, **F)** Outer ring represents the homozygous SNP density distribution and the inner ring denotes the heterozygous SNP density distribution, **G)** Outer ring represents the homozygous InDel density distribution and the inner ring denotes the heterozygous InDel density distribution.

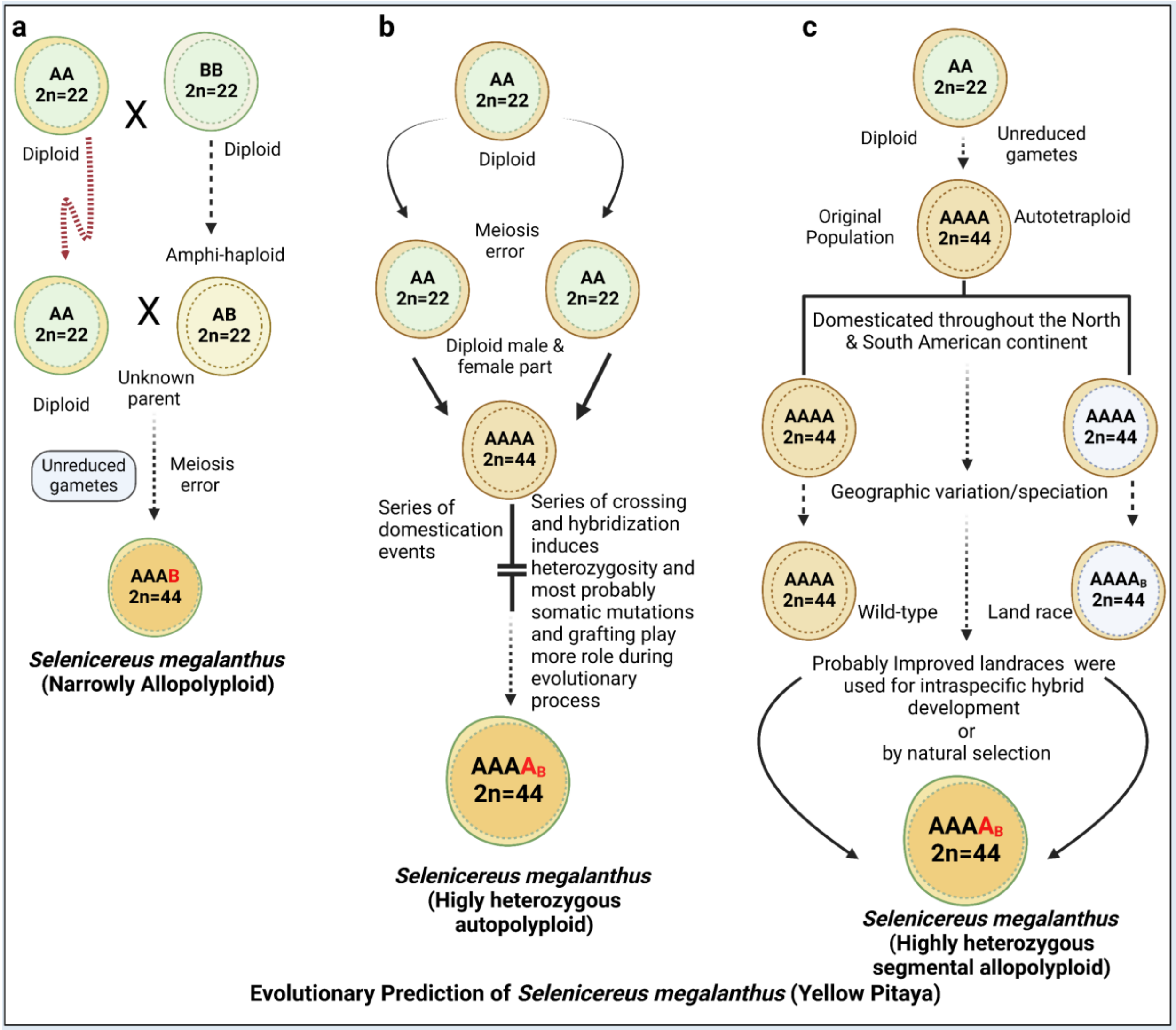


**Supplementary Figure 5: Predicting the ploidy level and divergent evolution in yellow pitaya (*Selenicereus* megalanthus) based on assumptions.** In our Smudgeplot analysis, we identified the AAAB pattern as a dominant component accounting for 64% of the examined K-mers. These results collectively point out a high level of heterozygosity in *Selenicereus megalanthus* which might be a reason for clonal propagation, or somatic mutations may be included in a heterozygous manner within a clone, or narrow allopolyploid. We propose three assumptions, **a)** Unknown parents with the genotype of AA and BB crossed to produce an Amphi-haploid. Amphi-haploid crossed again with diploid parent AA and transferred the same chromosome complement as somatic cells of the parent individual. From this cross, we can predict the yellow pitaya genotype as AAAB which is majorly autotetraploid but narrowly allopolyploid. **b)** Due to the failure of meiotic cell division to separate sister chromatids into daughter nuclei, produced a plant with two sets of chromosomes. A series of domestication events and somatic mutations produced a highly heterozygous genome. **c)** Our 3rd assumption is, after meiotic failure, plants were grown in specific areas. After the splitting of the Gondwana continent into South America and Africa, plants were produced distantly. Due to clonal propagation, environmental mutations, domestication and somatic mutation events created distant organisms and evolutionary speciation within the species. Most probably, Aztec people produced the hybrid by crossing the distant pitaya plants of closely related yellow pitaya.

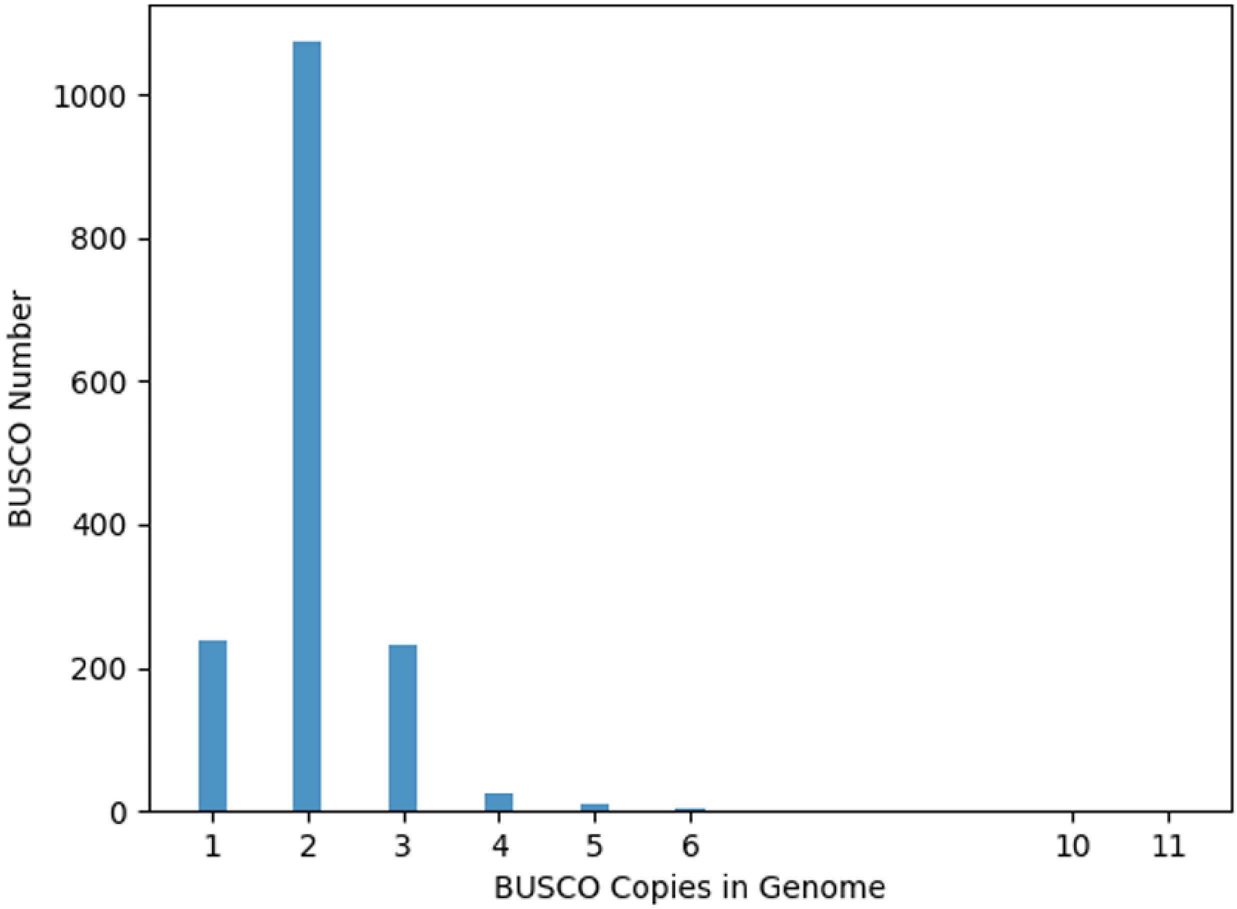


**Supplementary Figure 6: Distribution map of the BUSCO copy number of the *Selenicereus megalanthus* genome.** The results predicted the number of BUSCO copies in the yellow pitaya genome with 1 copy, 2 copies, 3 copies, and more copies as shown on x-axis of the graph.

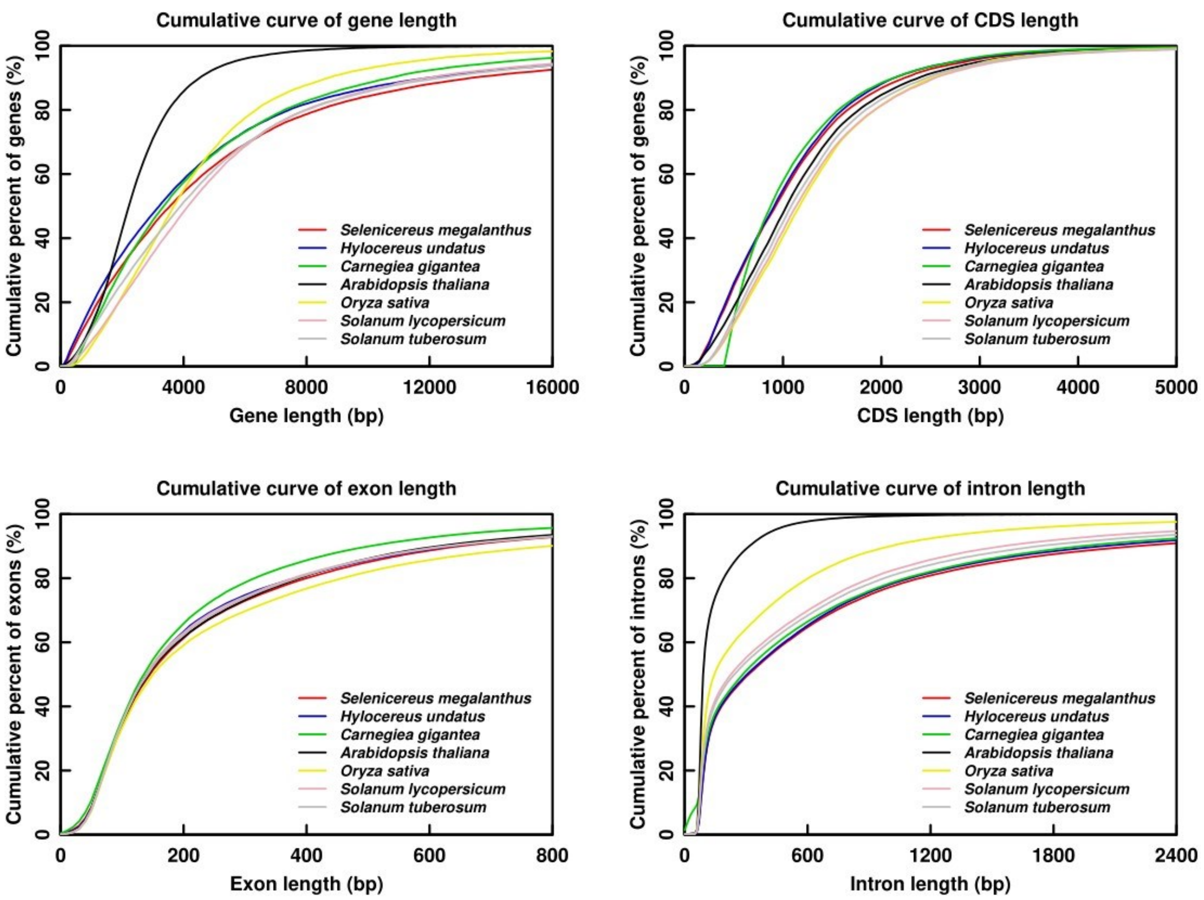


**Supplementary Figure 7: Cumulative distribution of gene set with closely related species.** Species includes yellow pitaya (*Selenicereus megalanthus*), red peel white flesh pitaya (*Selenicereus undatus*), Saguaro cactus (*Carnegiea gigantea*), Arabidopsis (*Arabidopsis thaliana*), tomato (*Solanum lycopersicum*), and potato (*Solanum tuberosum*).

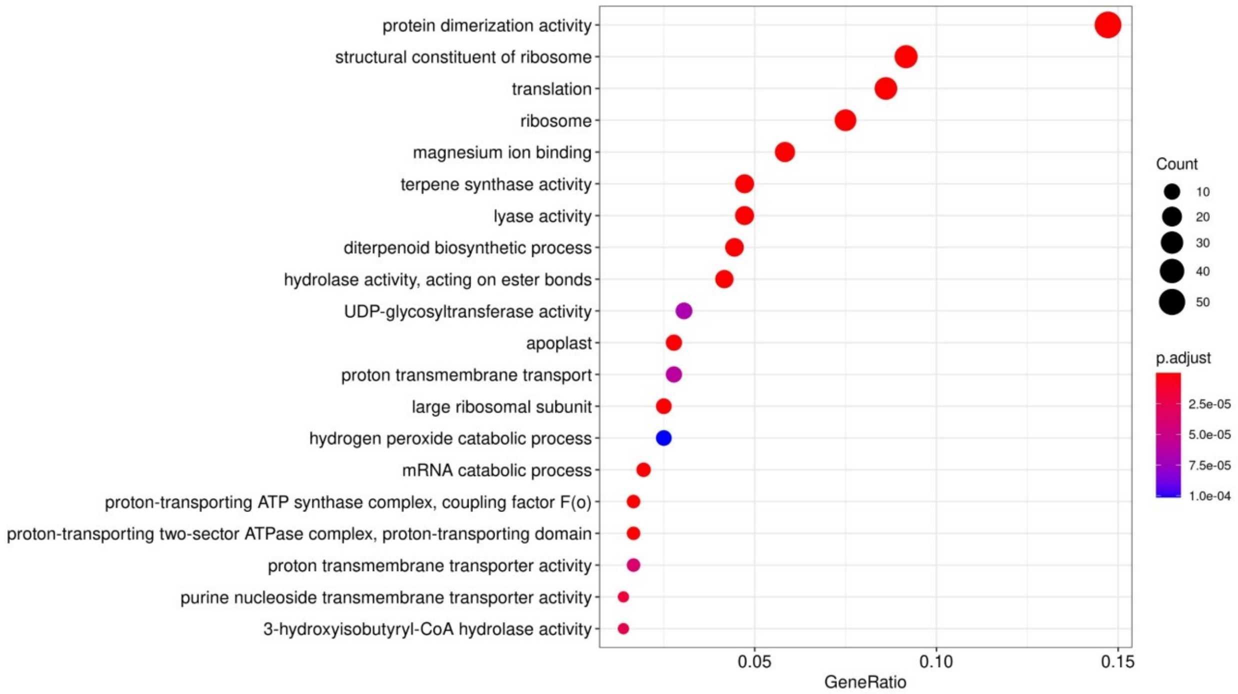


**Supplementary Figure 7: GO enrichment bubble map.** The x-axis is the enrichment ratio and the y-axis represents the GO-terms.

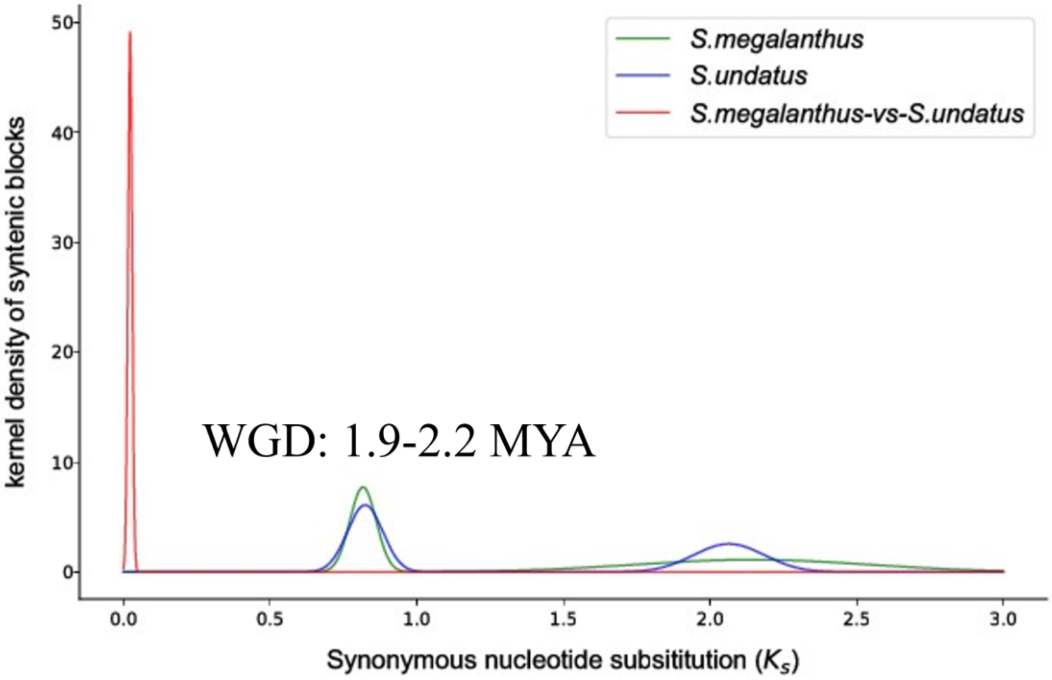

**Supplementary Figure 8**: Ks distribution map. The peak value of the combined map between the species represents species divergence and the peak value of the combined map within a species represents the whole genome duplication event of *S. megalanthus*.

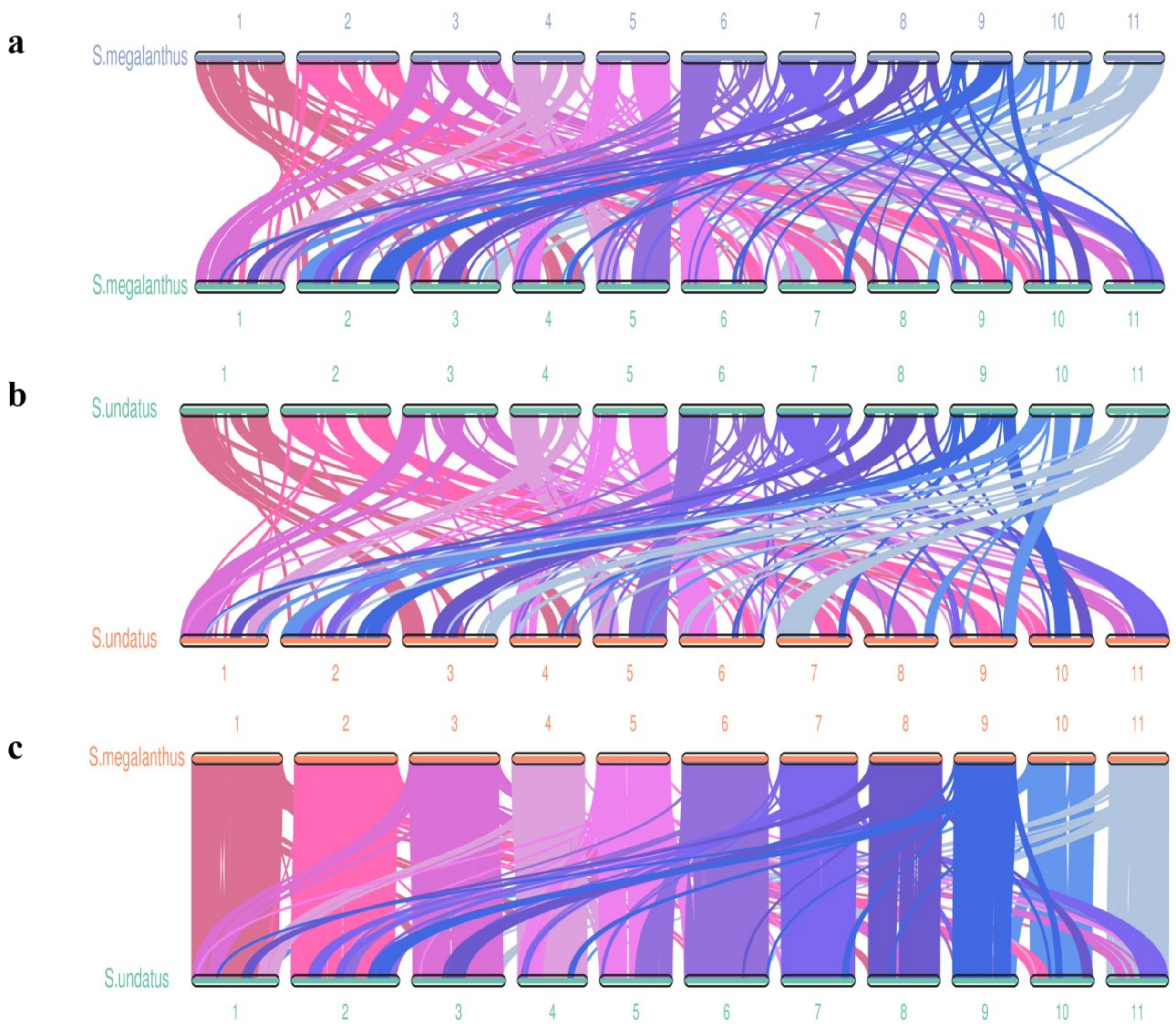


**Supplementary Figure 9: Syntenic collinearity diagram of diploid and polyploid pitaya species. a)** Syntenic blocks of yellow pitaya (*S. megalanthus*) of each chromosome are exhibited that have the same order on another chromosome of the same species. **b)** Syntenic blocks of red-peel pitaya (*S. undatus*) of each chromosome are shown that have the same order on another chromosome of red-peel pitaya. **c)** Syntenic blocks of *S. megalanthus* exhibited collinearity with the orthologs of *S. undatus*.

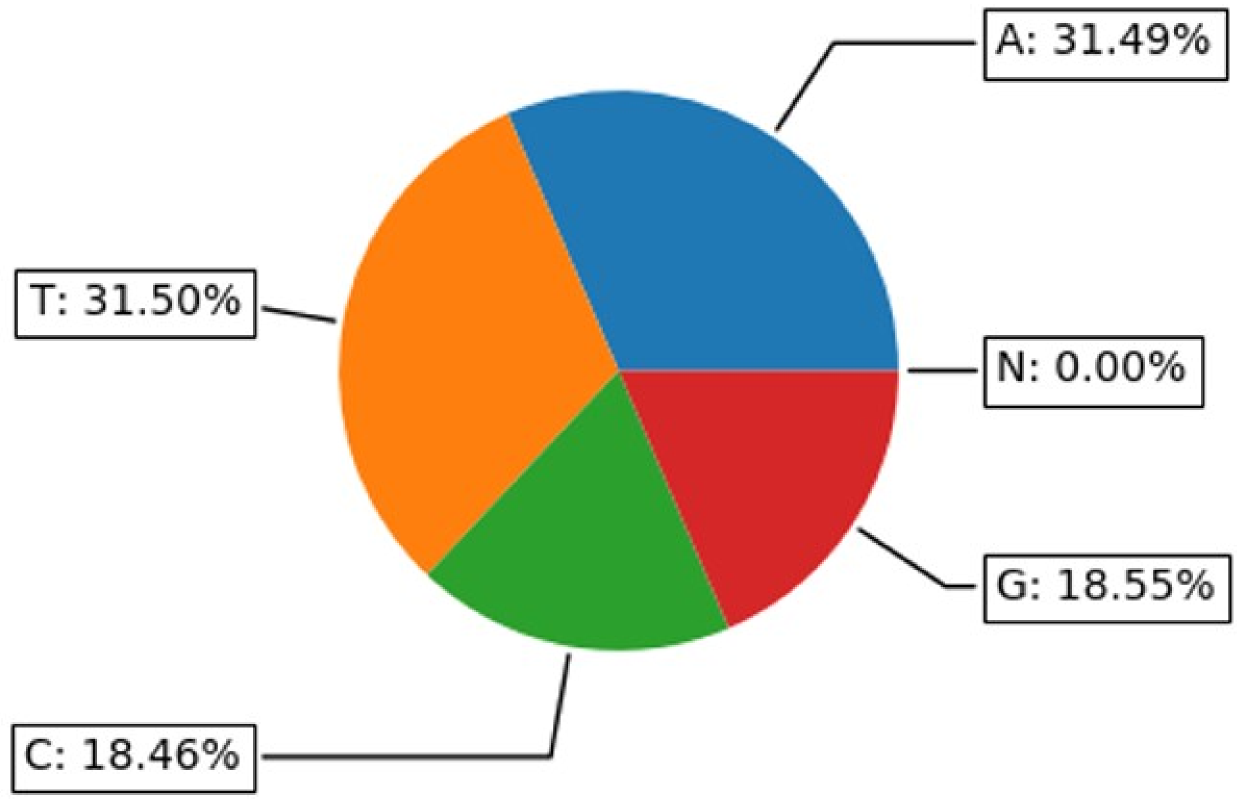


**Supplementary Figure 10:** Statistics on the distribution of bases (Adenine, Guanine, Cytosine, Thymine) composition in *S. megalanthus* genome.

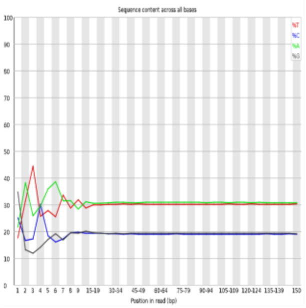

***S. undatus*** **Read 1, Replication 1**

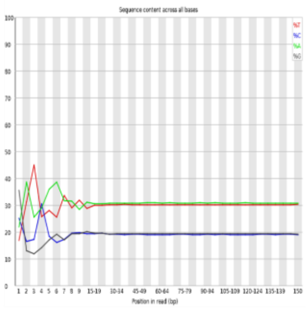

***S. undatus*** **Read 1, Replication 2**

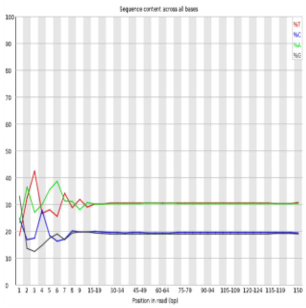

***S. undatus*** **Read 2, Replication 1**

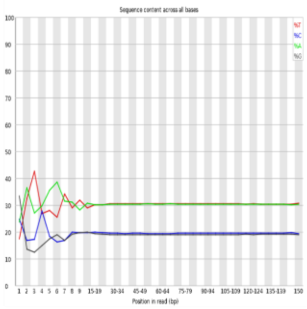

***S. undatus*** **Read 2, Replication 2**

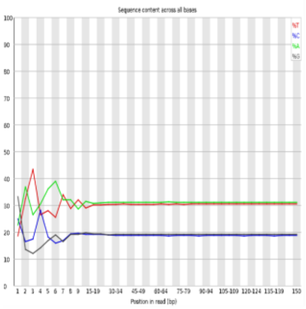

***S. megalanthus*** **Read 1, Replication 1**

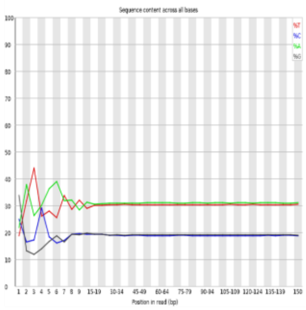

***S. megalanthus*** **Read 1, Replication 2**

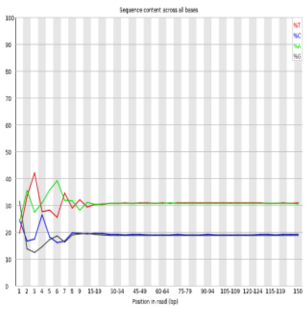

***S. megalanthus*** **Read 2, Replication 1**

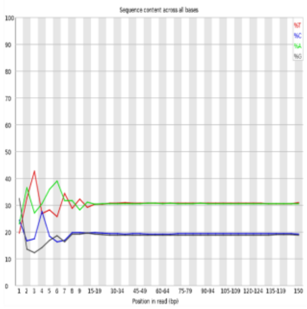

***S. megalanthus*** **Read 2, Replication 2**

**Supplementary Figure 11: Base content distribution of diploid and polyploid pitaya species along the length of read 1 and read 2.** Note: The horizontal axis denotes the base position of the reads, and the longitudinal axis represents the proportion of ATCG bases at that position.

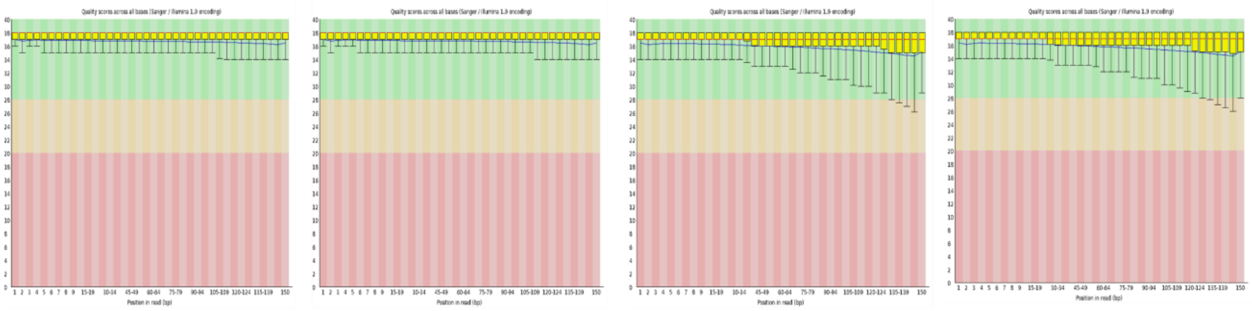

***S. undatus*** **Read 1, Replication 1** ***S. undatus*** **Read 1, Replication 2** ***S. undatus*** **Read 2, Replication 1** ***S. undatus*** **Read 2, Replication 2**

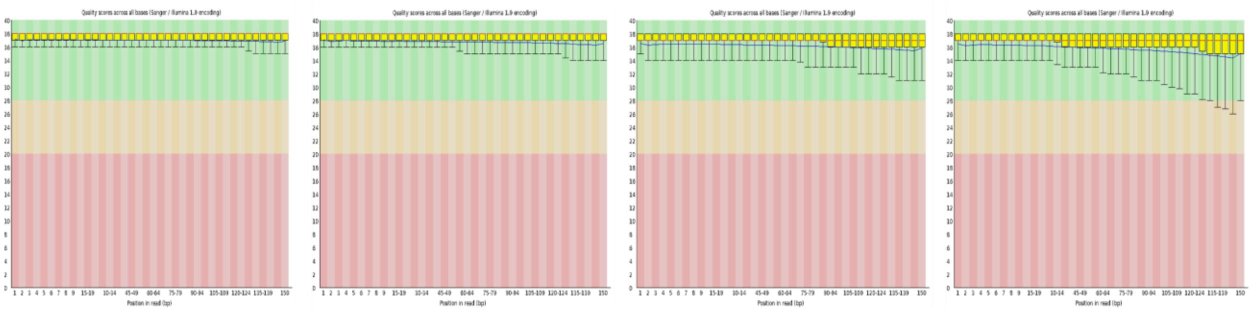

***S. megalanthus*** **Read 1, Replication 1** ***S. megalanthus*** **Read 1, Replication 2** ***S. megalanthus*** **Read 2, Replication 1** ***S. megalanthus*** **Read 2, Replication 2**

**Supplementary Figure 12: Base mass distribution of diploid and polyploid pitaya species along the length of read 1 and read 2.** Note: The horizontal axis exhibits the base position of the reads, and the longitudinal axis represents the average mass value of the base at that position.

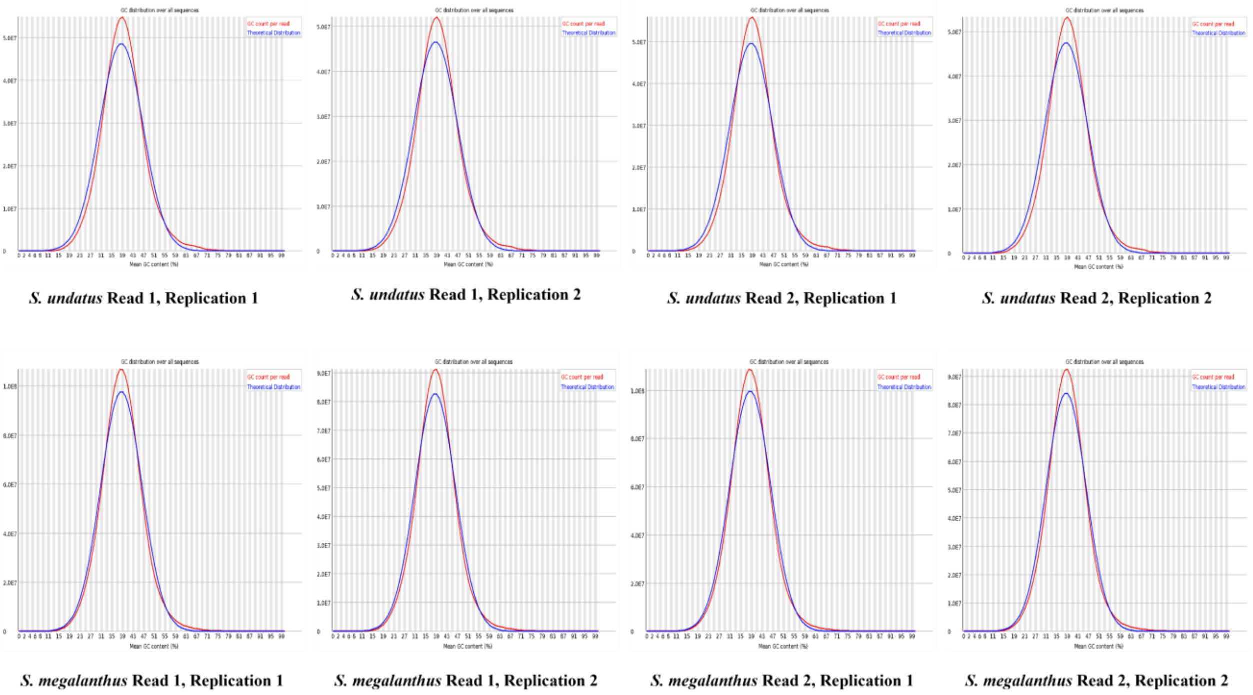


**Supplementary Figure 13: GC content distribution of diploid and polyploid pitaya species along the length of read 1 and read 2.** Note: The horizontal axis shows the percentage of GC content while the vertical axis exhibits the number of reads of this GC content.

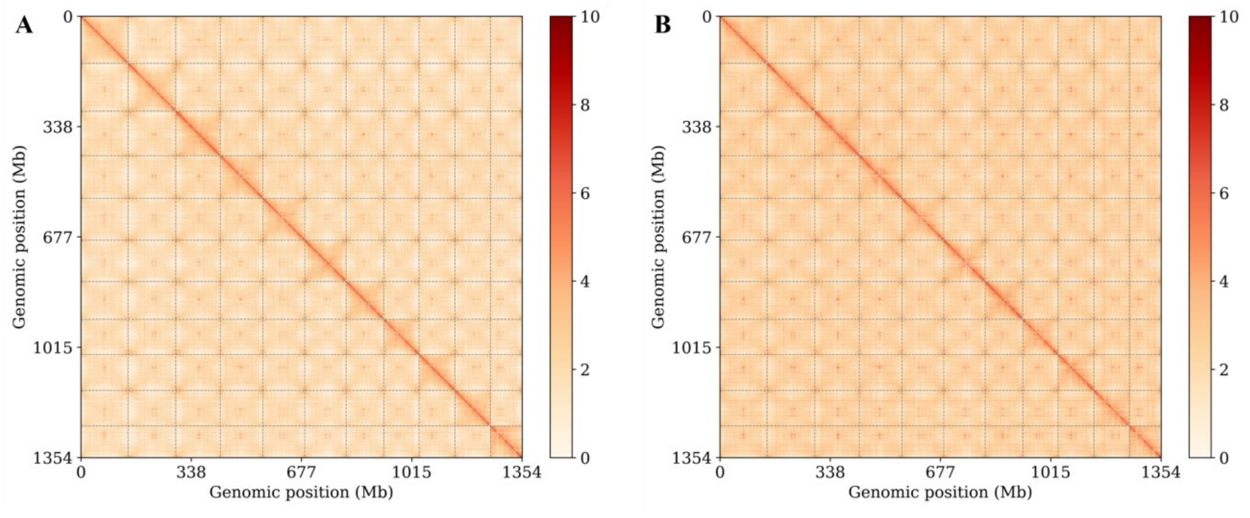


**Supplementary Figure 14: Heatmap of genome-wide interactions at 100-kb resolution. A)** *S. undatus* **B)** *S. megalanthus*. Note: the horizontal and vertical axes denote the positions on the reference genome. Color bars exhibit the intensity of the interaction, the higher the intensity, the stronger the interaction. The box along the diagonal region represents the interaction within the same chromosome (Cis). Unit: Bin

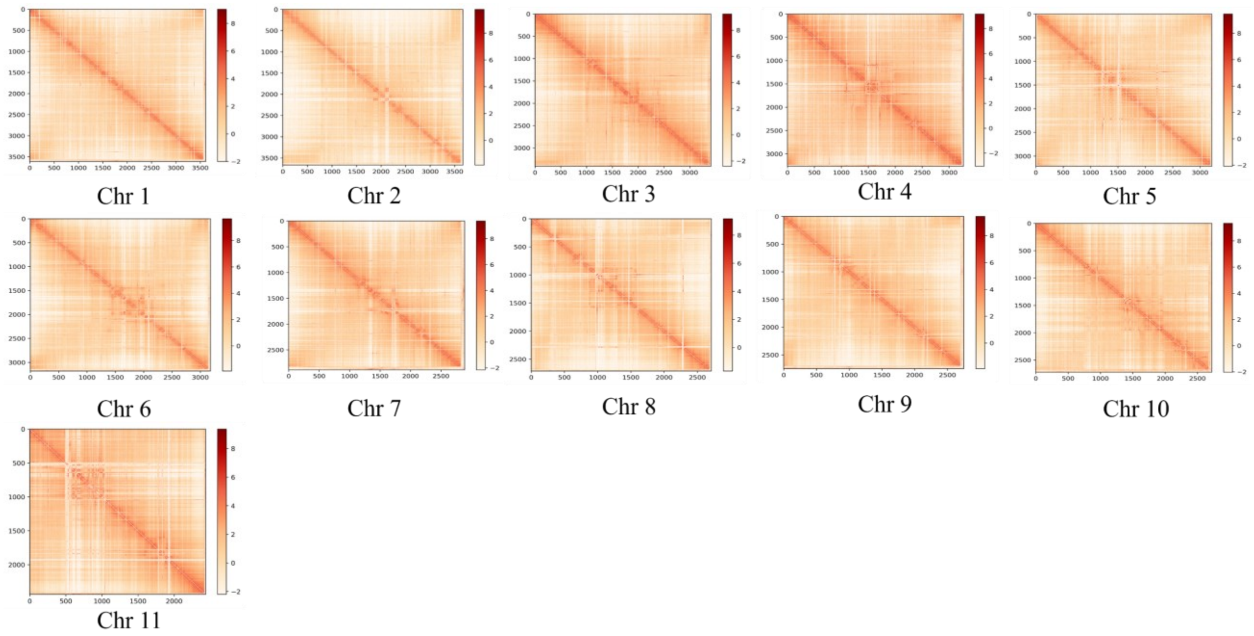


**Supplementary Figure 15: Single-chromosomal interaction heatmap with a resolution at 40-kb of diploid pitaya (*S. undatus*).** Note: The horizontal and vertical axis shows the position of the chromosome. The color bar exhibits the intensity of the interaction, the higher its intensity, the stronger the interaction. Along the diagonal region: the close interaction is strong, forming a three-dimensional structure. Unit: Bin.

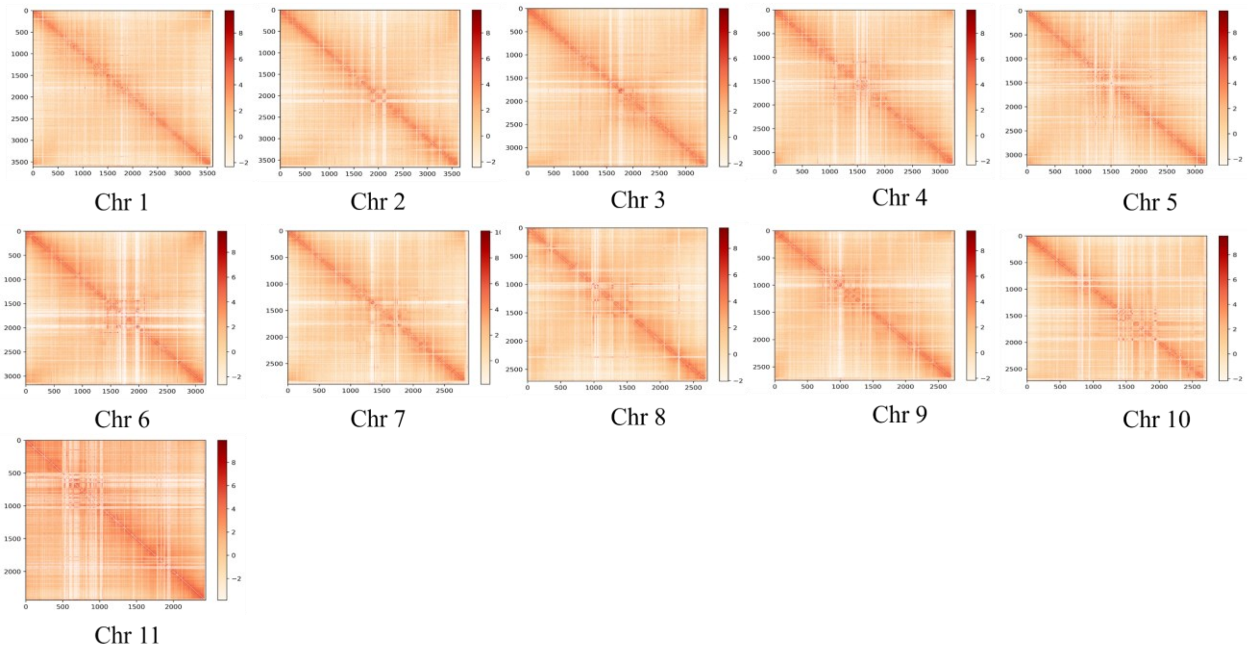

**Supplementary Figure 16: Single-chromosomal interaction heatmap with a resolution of 40 kb of polyploid pitaya (*S. megalanthus*).** Note: The horizontal and vertical axis shows the position of the chromosome. The color bar exhibits the intensity of the interaction, the higher its intensity, the stronger the interaction. Along the diagonal region: the close interaction is strong, forming a three-dimensional structure. Unit: Bin.

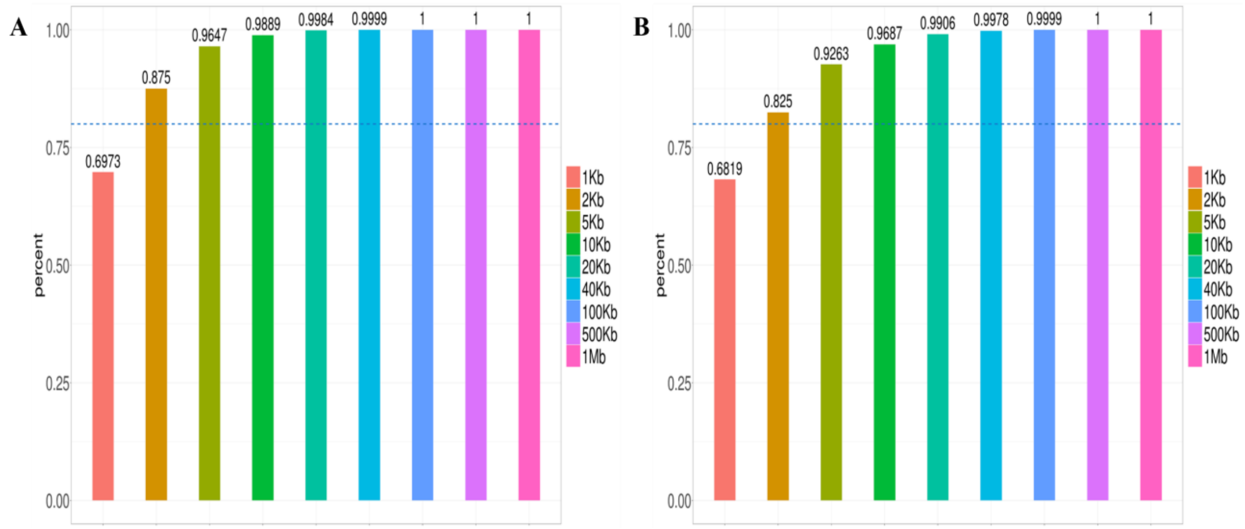


**Supplementary Figure 17: Resolution chart. A) *S. undatus* B) *S. megalanthus*.** Note: The horizontal axis shows different resolutions. The vertical axis exhibits the proportion of bins with more than 1000 contacts in the total bin. The horizontal dashed line denotes 80%, more than the horizontal dashed line, which means that the resolution supports subsequent analysis.

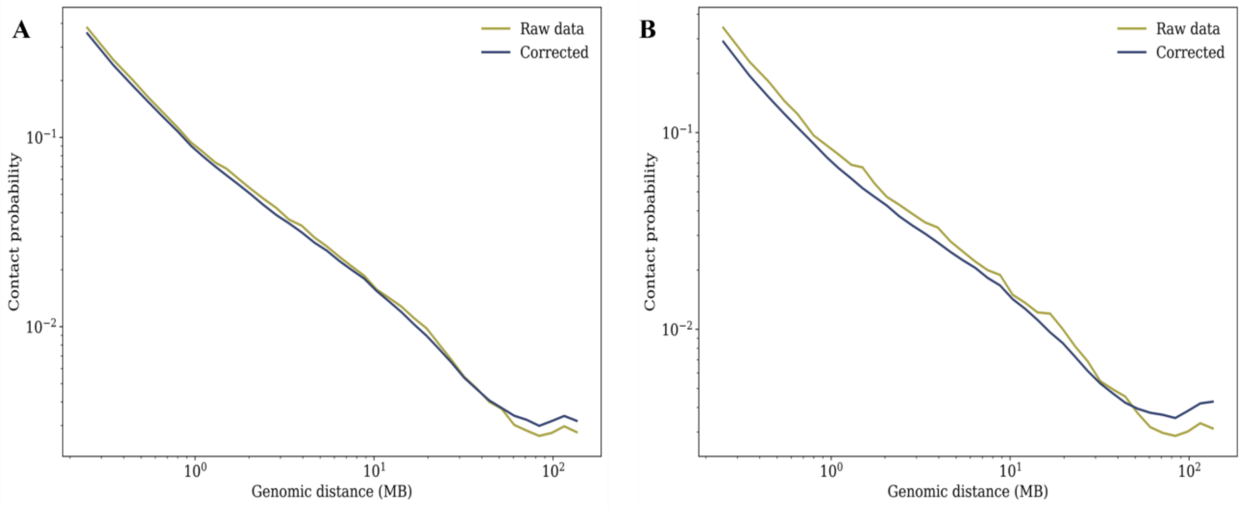

**Supplementary Figure 18: Genome-wide interaction attenuation curve. A)** *S. undatus* **B)** *S. megalanthus*. The transverse axis denotes the relative distance between cis-interaction sites, Longitudinal axis exhibits the interaction frequency of this distance. unit: Mb.

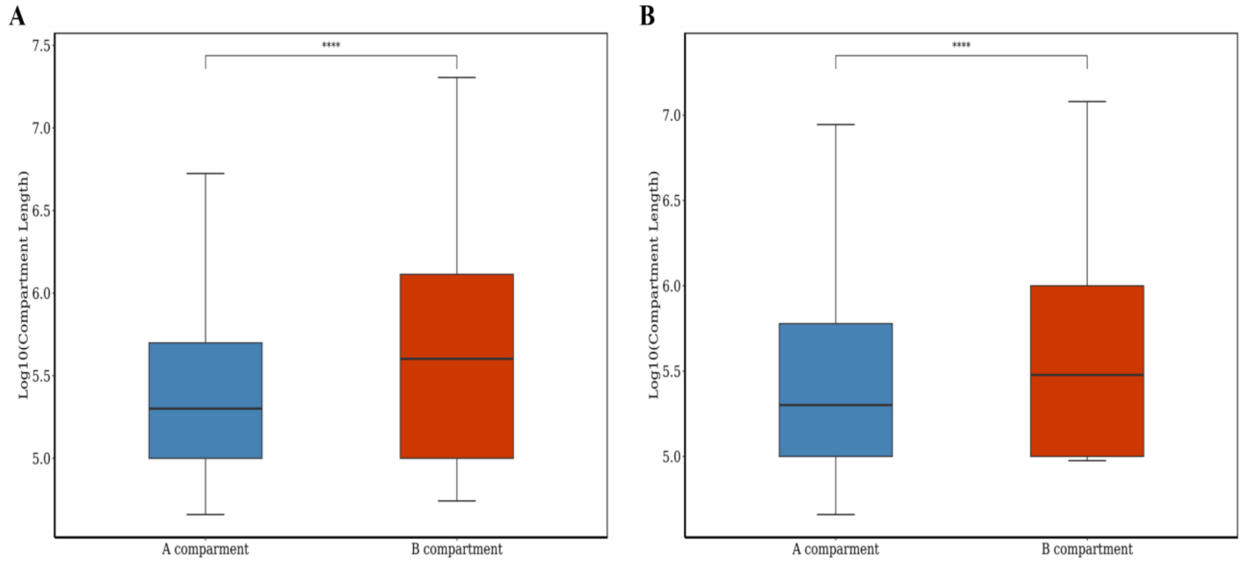


**Supplementary Figure 19: Length distribution of the compartments. A)** *S. undatus* **B)** *S.* megalanthus. The horizontal axis exhibits Compartment A and Compartment B. The Vertical axis denotes the length of the compartment (unit: bp), taking log10. Each box plot has five statistics: after removing the discrete values, the top to bottom are the maximum, upper quartile, median, lower quartile, and minimum values. The significant test results in the figure **** represent 0.0001 ≥ p.

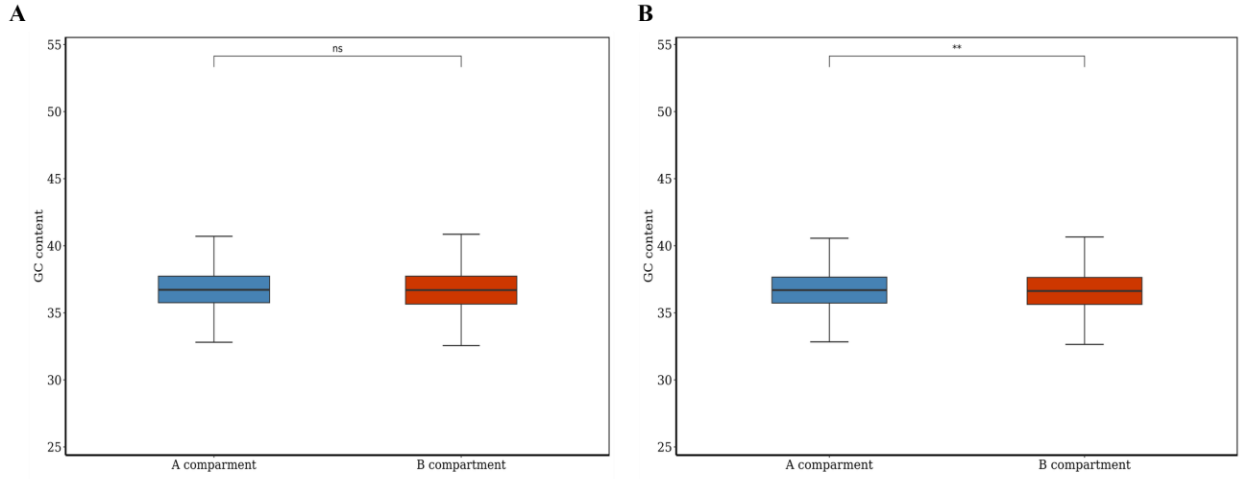

**Supplementary Figure 20: GC content distribution of the compartments. A)** *S. undatus* **B)** *S.* megalanthus. The horizontal axis exhibits Compartment A and Compartment B. The vertical axis denotes GC content in each bin in Compartment, unit: %. Each box plot has five statistics: after removing the discrete values, the top to bottom are the maximum, upper quartile, median, lower quartile, and minimum values. The significant test results in figure "ns" represents $p > 0.05$ and ** represents $0.01 \geq p > 0.001$.

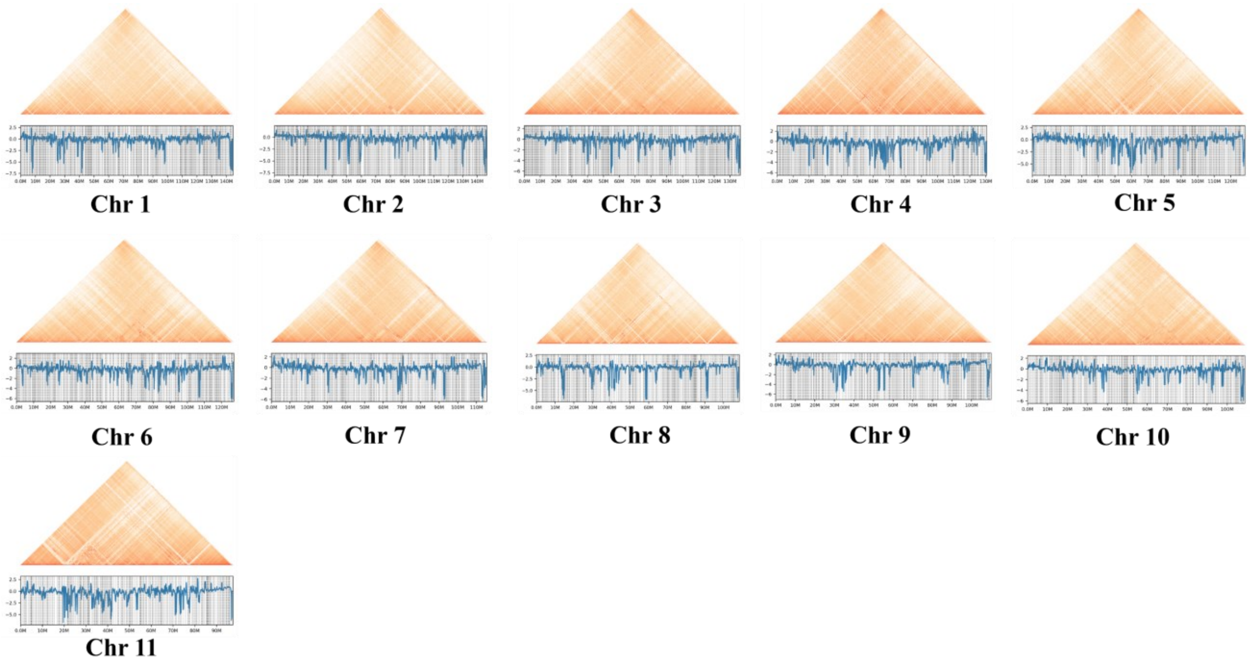


**Supplementary Figure 21: Single chromosome topologically associated domains of diploid pitaya (*S. undatus*).** Note: The horizontal axis represents a position on the reference genome (Unit: Mb). The upper part of the figure shows a heat map of the interaction of a single chromosome. However, the lower part of the figure exhibits a vertical axis, the blue line shows the Insulation Score, and the gray line denotes the TAD boundary.

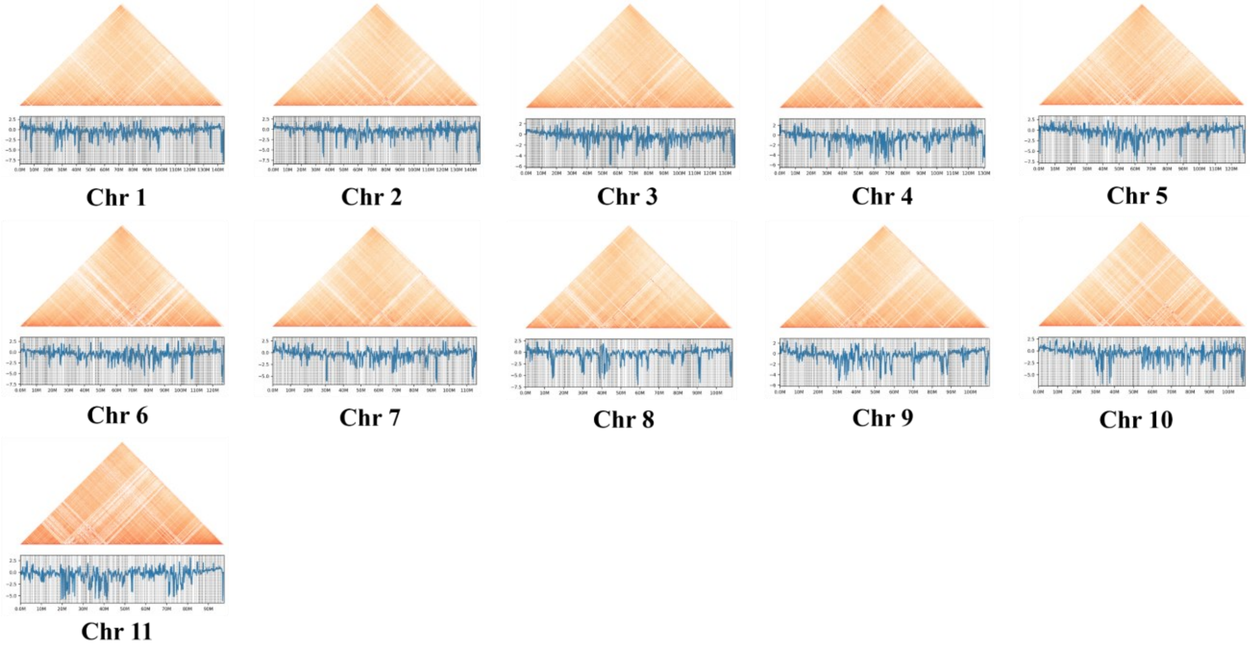


**Supplementary Figure 22: Single chromosome topologically associated domains of polyploid pitaya (*S. megalanthus*).** Note: The horizontal axis represents a position on the reference genome (Unit: Mb). The upper part of the figure shows a heat map of the interaction of a single chromosome. However, the lower part of the figure exhibits a vertical axis, the blue line shows the Insulation Score, and the gray line denotes the TAD boundary.

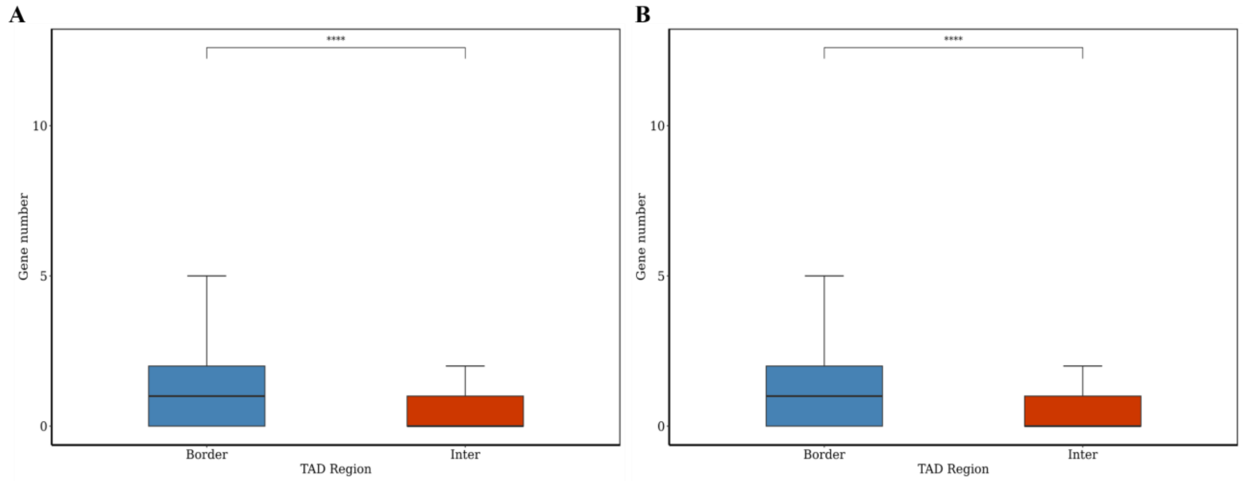


**Supplementary Figure 23: Map of the number of genes within the boundaries of the TAD. A)** *S. undatus* **B)** *S.* megalanthus. The horizontal axis denotes the TAD boundary (Border) and TAD internal (Inter). The longitudinal axis represents the number of genes in each bin in the TAD boundary (Border) or TAD (Inter) (unit: 3). The significance test results in the figure **** represents 0.0001 ≥ p.

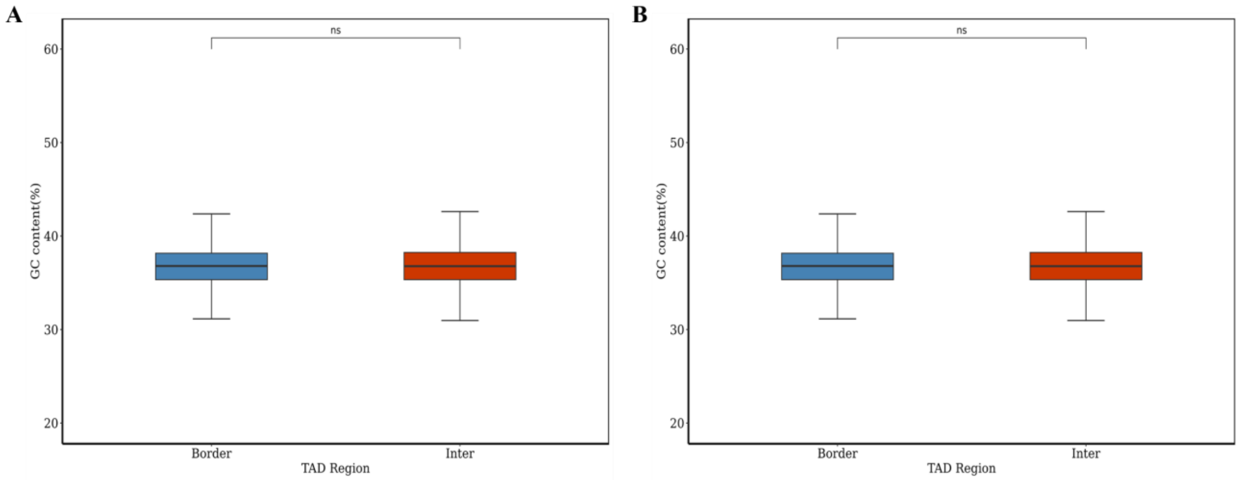


**Supplementary Figure 24: The distribution of GC content at the boundary of TAD and within TAD. A)** *S. undatus* **B)** *S.* megalanthus. Note: The horizontal axis represents the TAD boundary (Border) and TAD internal (Inter). The longitudinal axis exhibits the GC content in each bin in the TAD boundary (Border) or TAD internal (Inter) (unit: %). The significance test results in the graph ns represents $p > 0.05$.

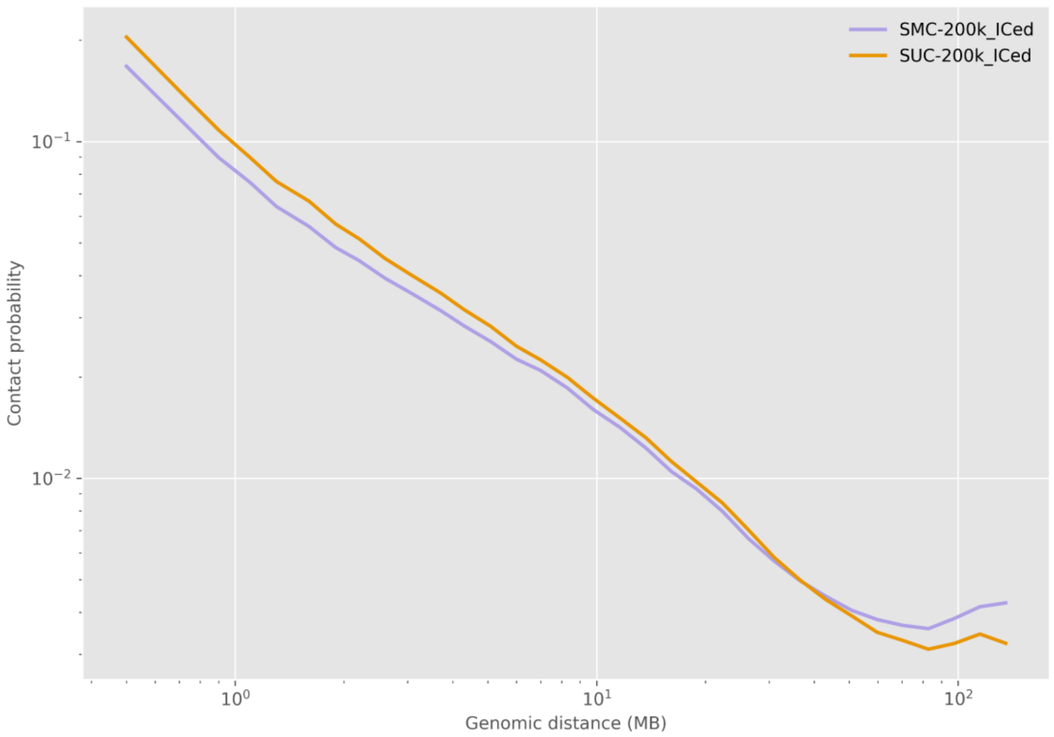


**Supplementary Figure 25: Genome-wide distance-interaction frequency plot at 40Kb resolution between diploid and polyploid pitaya.** We combined the interaction attenuation curve of the whole genome of diploid pitaya and polyploid pitaya at 40-kb resolution to perform a comparative analysis of the differences in distance and the interaction frequency in both species. Note: SMC denotes the sample of polyploid pitaya (*Selenicereus megalanthus*) subjected to Hi-C sequencing and SUC exhibits the sample of diploid pitaya (*Selenicereus undatus*) subjected to Hi-C sequencing.

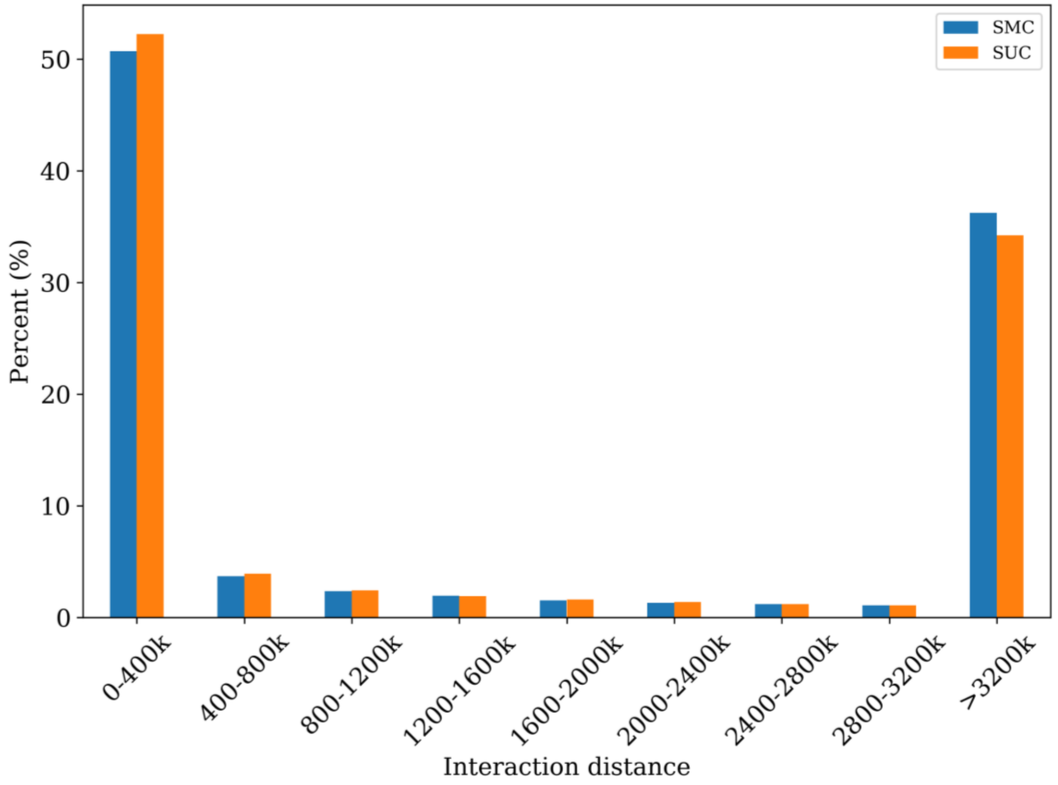


**Supplementary Figure 26: Proportional distribution of interactions at different distances between diploid pitaya and polyploid pitaya.** SMC denotes the sample of polyploid pitaya (*Selenicereus megalanthus*) subjected to Hi-C sequencing and SUC exhibits the sample of diploid pitaya (*Selenicereus undatus*) subjected to Hi-C sequencing.

.

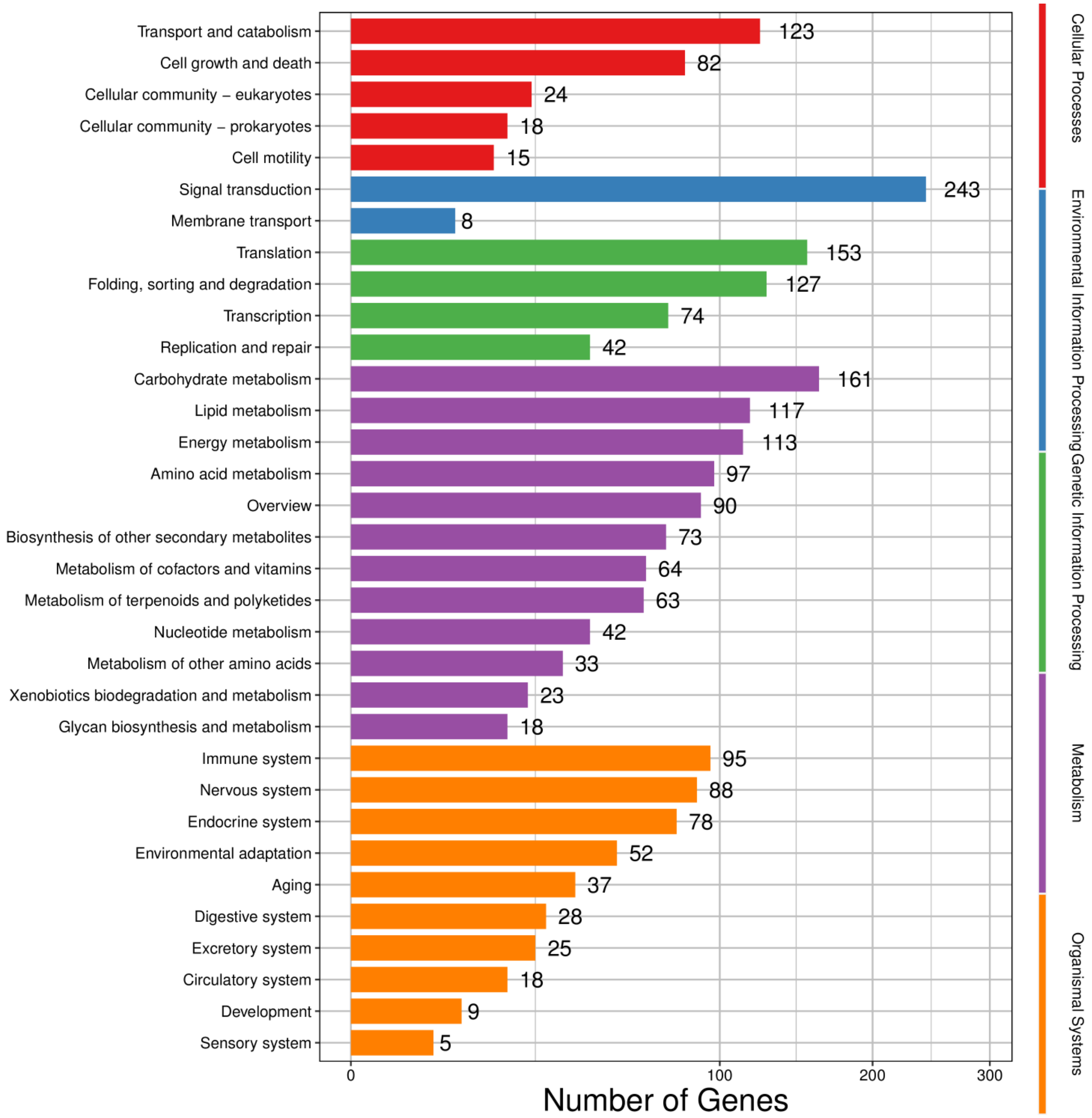


**Supplementary Figure 27: KEGG annotation of genes in differential compartments of diploid and polyploid pitaya.** The horizontal axis represents the number of genes annotated to the pathways. The vertical axis exhibits the name of the KEGG metabolic pathways. KEGG metabolic pathways are divided into six components including cellular process, environmental information, genetic information, metabolism, and organism systems.

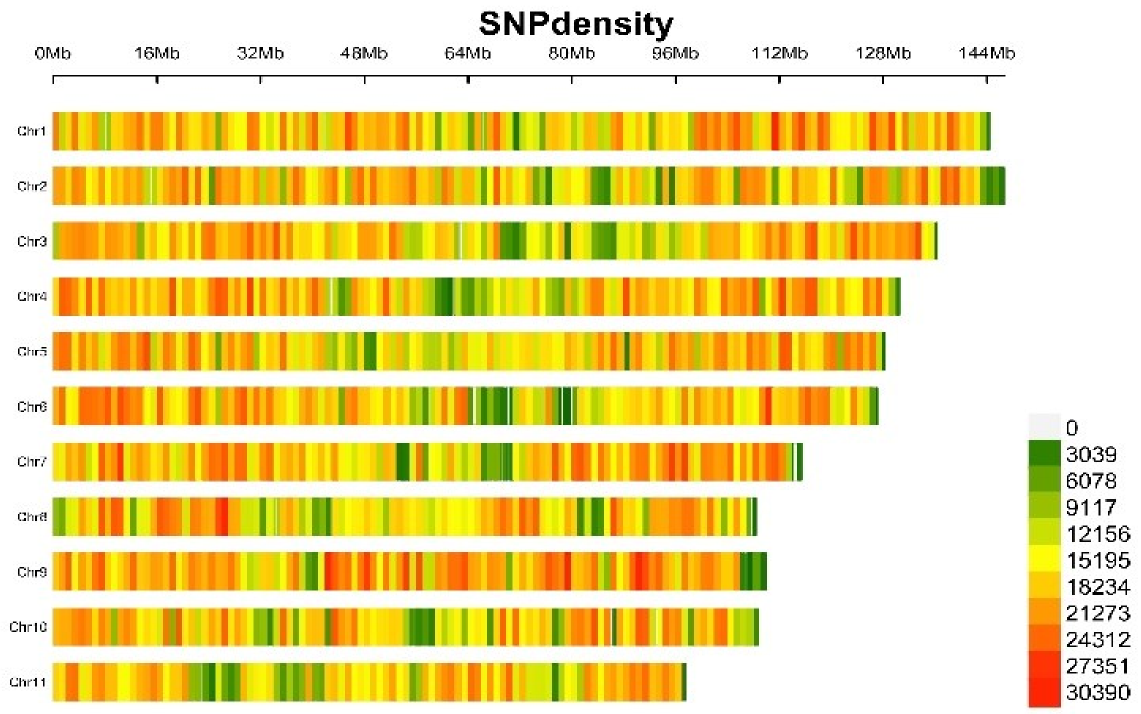


**Supplementary Figure 28:** The SNP density across chromosomes 1 to 11 of the yellow pitaya genome. The x-axis shows the physical distance alongside each chromosome (1Mb window) and the color gradient from red (high SNP density) to green (low SNP density) demonstrates the distribution of genetic variation.

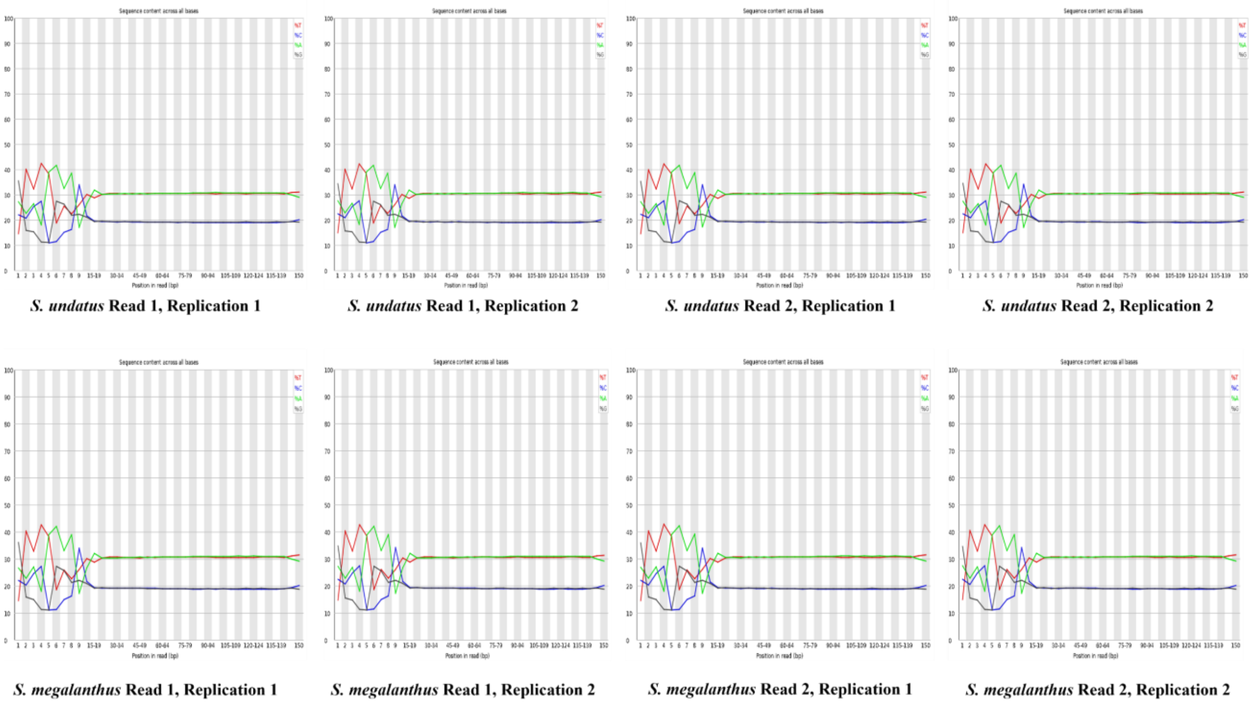


**Supplementary Figure 29: Base content distribution map of Read1 and Read2 of diploid and polyploid pitaya.** The horizontal axis is the base position of the reads while the vertical axis is the proportion if ATCG  bases.

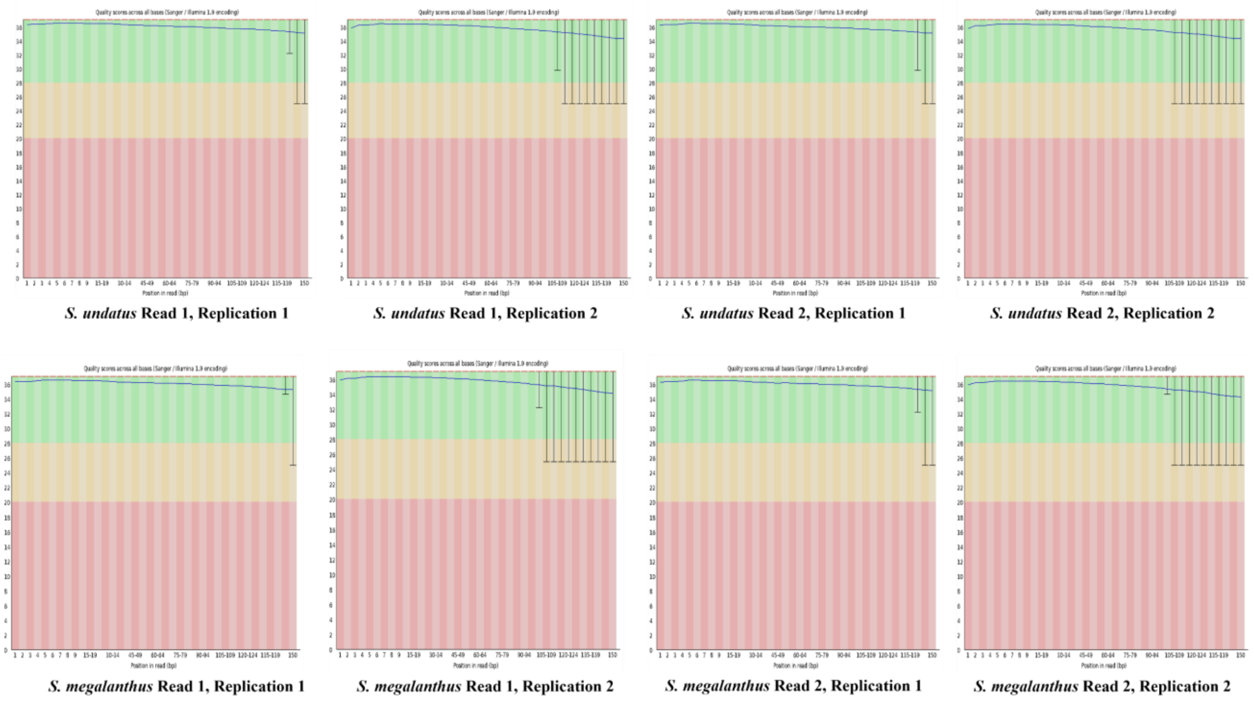

**Supplementary Figure 30: Base mass distribution map of Read and Read 2 of diploid and polyploid pitaya.** The horizontal axis is the base position of the reads while the vertical axis is the average mass value of the base.

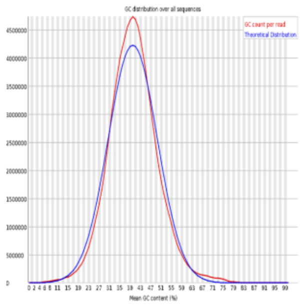


***S. undatus*** **Read 1, Replication 1**

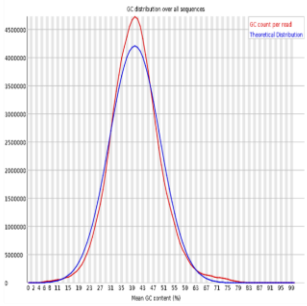


***S. undatus*** **Read 1, Replication 2**

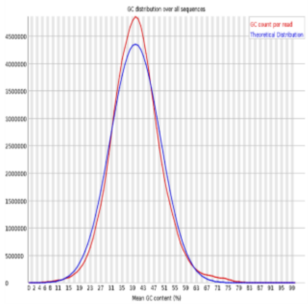


***S. undatus*** **Read 2, Replication 1**

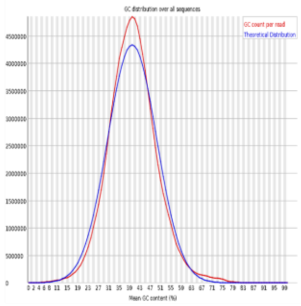


***S. undatus*** **Read 2, Replication 2**

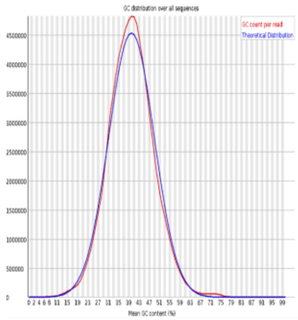


***S. megalanthus*** **Read 1, Replication 1**

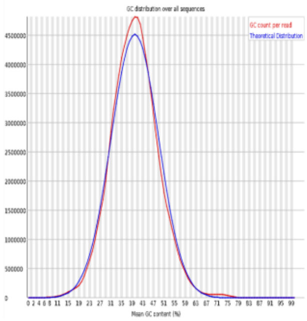


***S. megalanthus*** **Read 1, Replication 2**

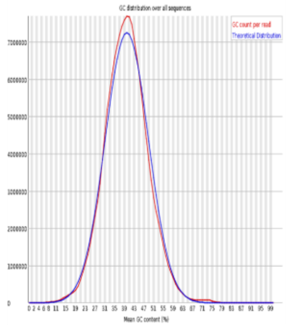


***S. megalanthus*** **Read 2, Replication 1**

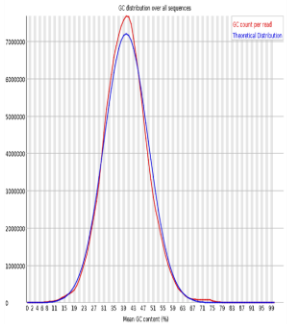


***S. megalanthus*** **Read 2, Replication 2**

**Supplementary Figure 31: GC content distribution of Read1 and Read2 of diploid and polyploid pitaya.** The horizontal axis is the percentage of GC content while the vertical axis is the number of reads for the GC content.

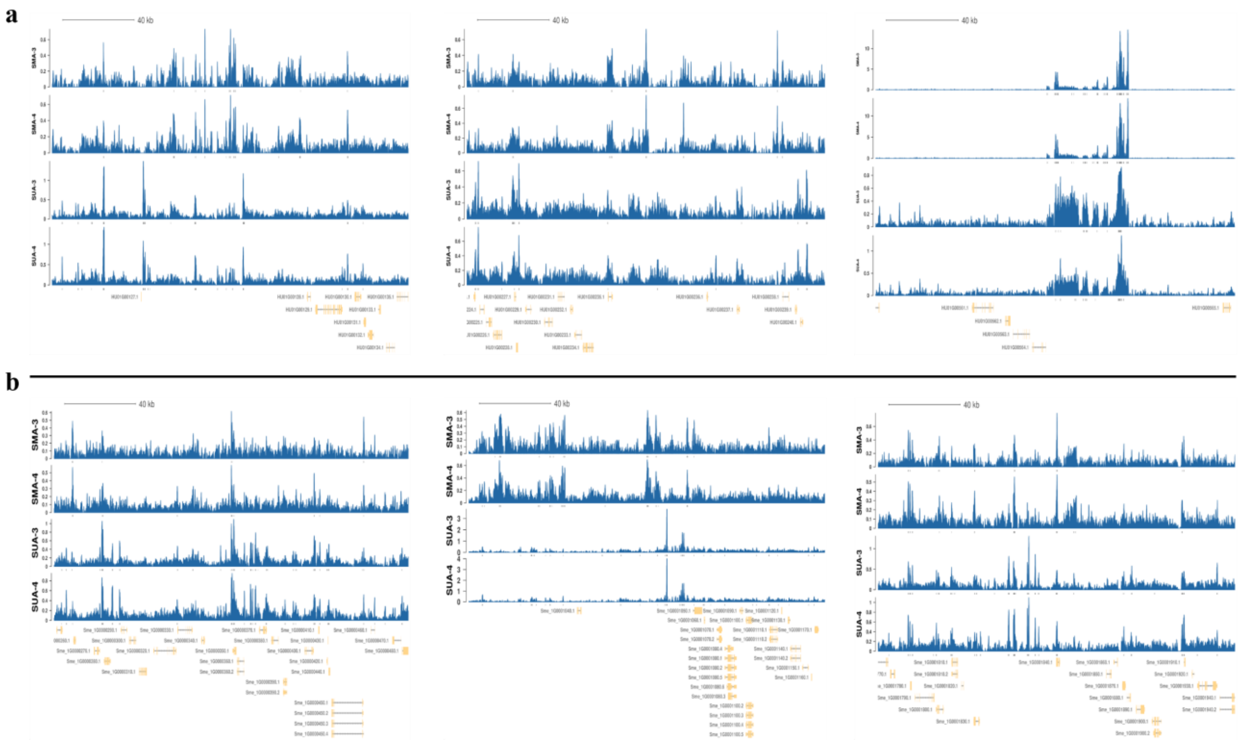


**Supplementary Figure 32: Visualization of ATAC-seq data to observe the significantly enriched areas in diploid and polyploid pitaya.** The horizontal axis is the position of the genome, and the vertical axis is the peak and gene annotation on **a)** diploid genome (*S. undatus*) **b)** polyploid genome (*S. megalanthus*).

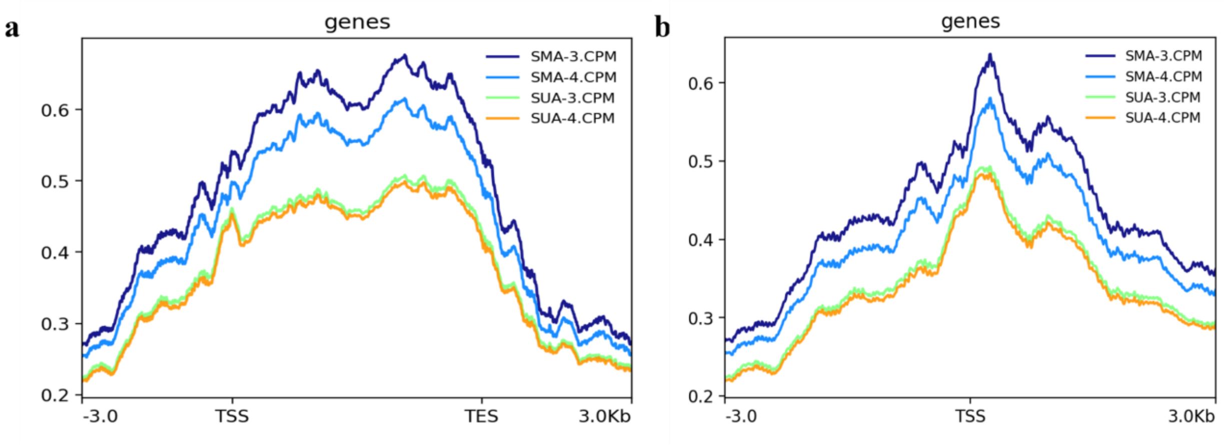


**Supplementary Figure 33: a) Trend plot of ATAC-seq signal distribution in the gene body region.** The horizontal axis is the position of the Gene body region and its upstream and downstream 3kb region, and the vertical axis is the average signal value of ATAC-seq. **b)** Trend plot of ATAC-seq signal distribution around TSS. The horizontal axis is the position of the 3kb upstream and downstream area of the TSS, and the vertical axis is the average signal value of ATAC-seq.

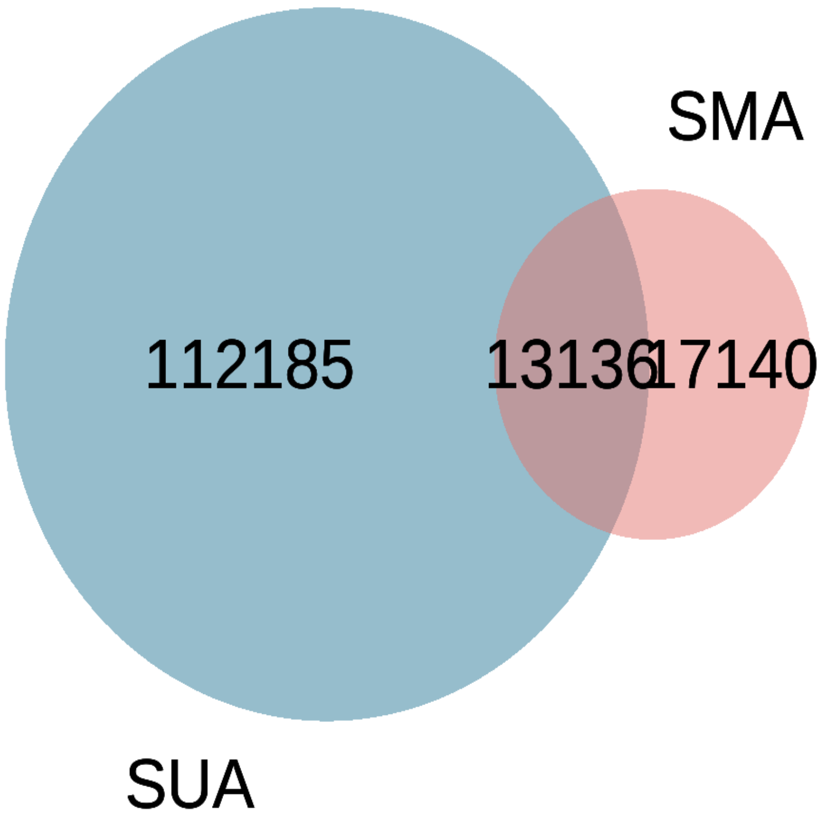


**Supplementary Figure 34: Venn plot of the overlapping peaks in diploid and polyploid pitaya.** SMA denotes the sample of polyploid pitaya (*Selenicereus megalanthus*) subjected to ATAC sequencing and SUA exhibits the sample of diploid pitaya (*Selenicereus undatus*) subjected to ATAC sequencing. 13136 peaks were shown as common between both species of pitaya.

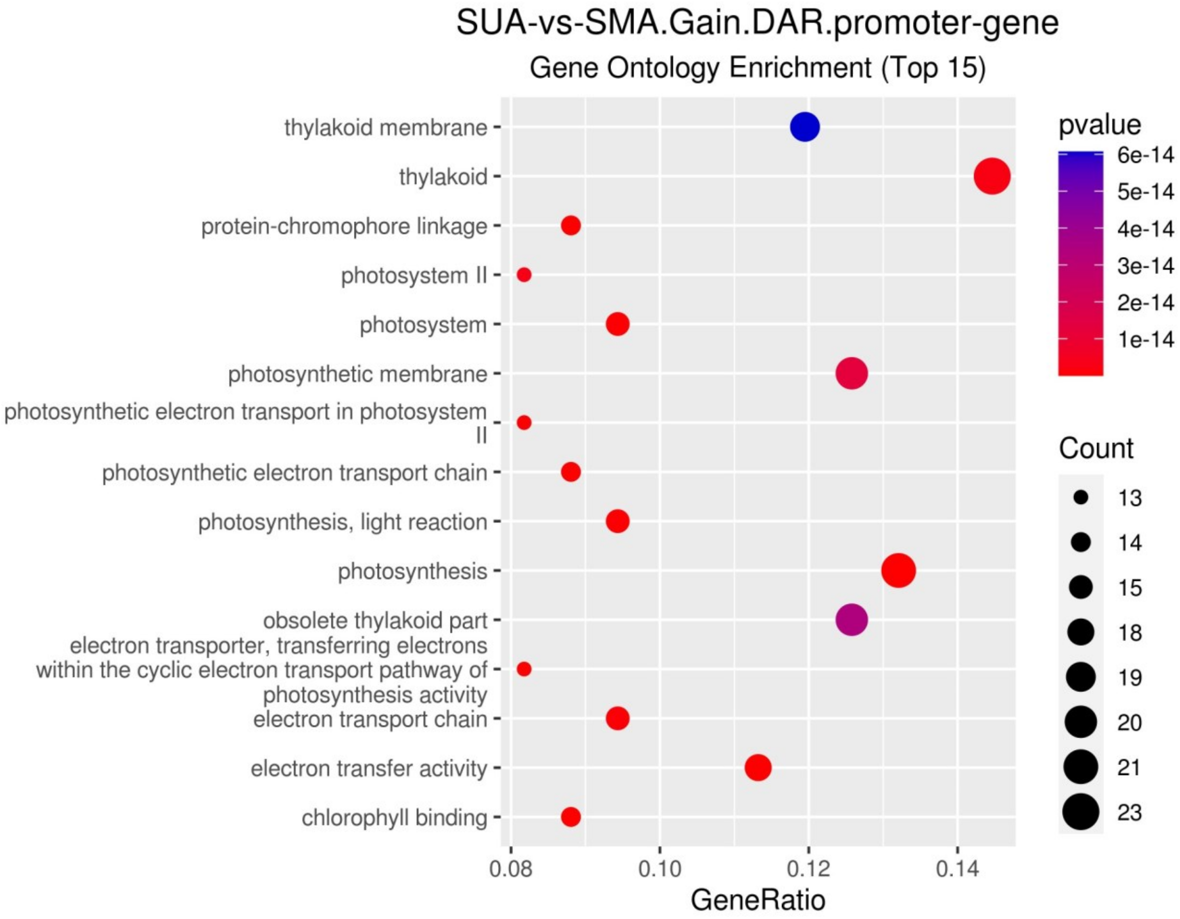


**Supplementary Figure 35: GO functional enrichment of Promoter genes associated with Gain DAR.** The horizontal axis is GeneRatio, the proportion of the number of genes enriched to the GO term in the gene set, and the vertical axis is the GO term. The color in the figure is the size of the p-value, and the circle is the size of the number of genes.

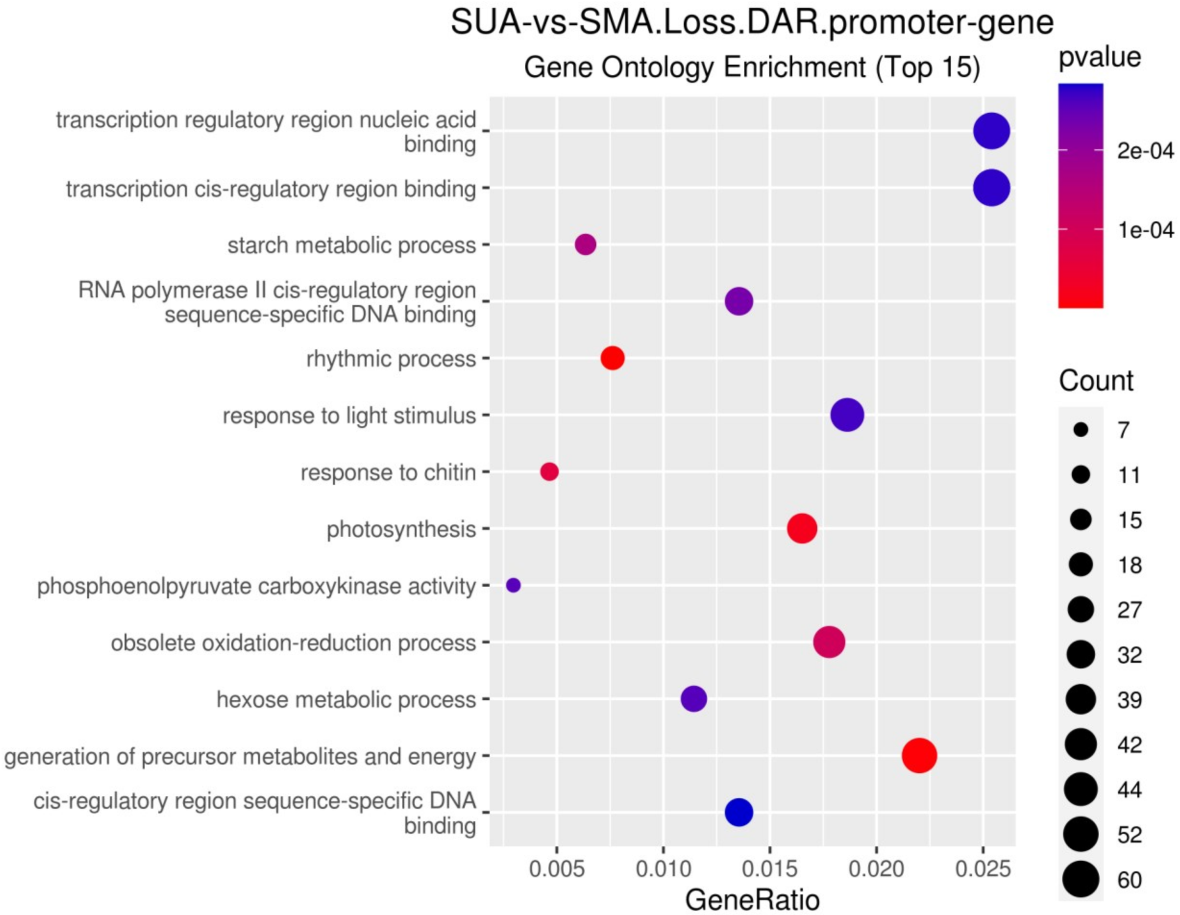


**Supplementary Figure 36: GO functional enrichment of Promoter genes associated with Loss DAR.** The horizontal axis is GeneRatio, the proportion of the number of genes enriched to the GO term in the gene set, and the vertical axis is the GO term. The color in the figure is the size of the p-value, and the circle is the size of the number of genes.

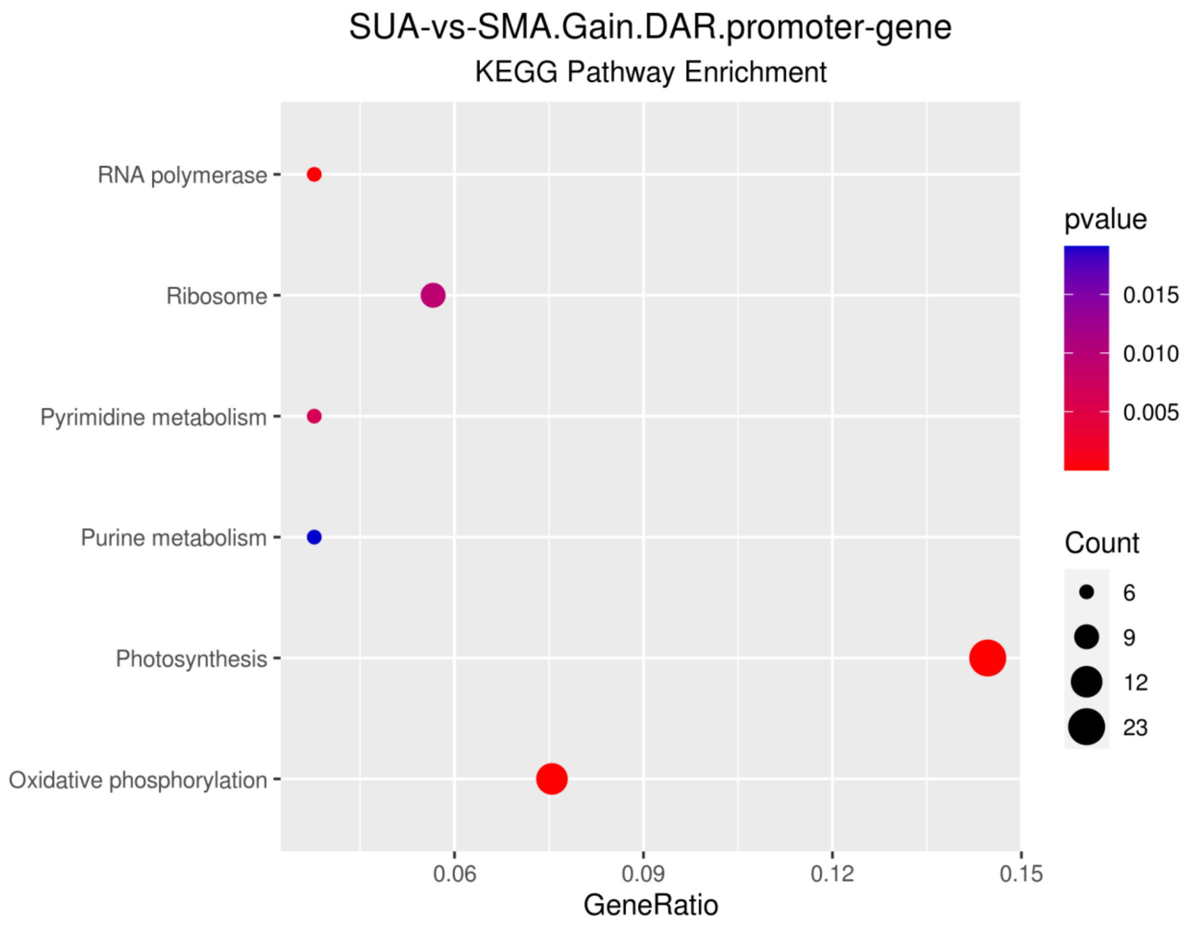


**Supplementary Figure 37: KEGG functional enrichment of the promoter gene of Gain DAR.** The horizontal axis is the GeneRatio, the proportion of the number of genes enriched to the pathway in the gene set, and the vertical axis is the Pathway. The color in the figure is the size of the p-value, and the circle is the size of the number of genes.

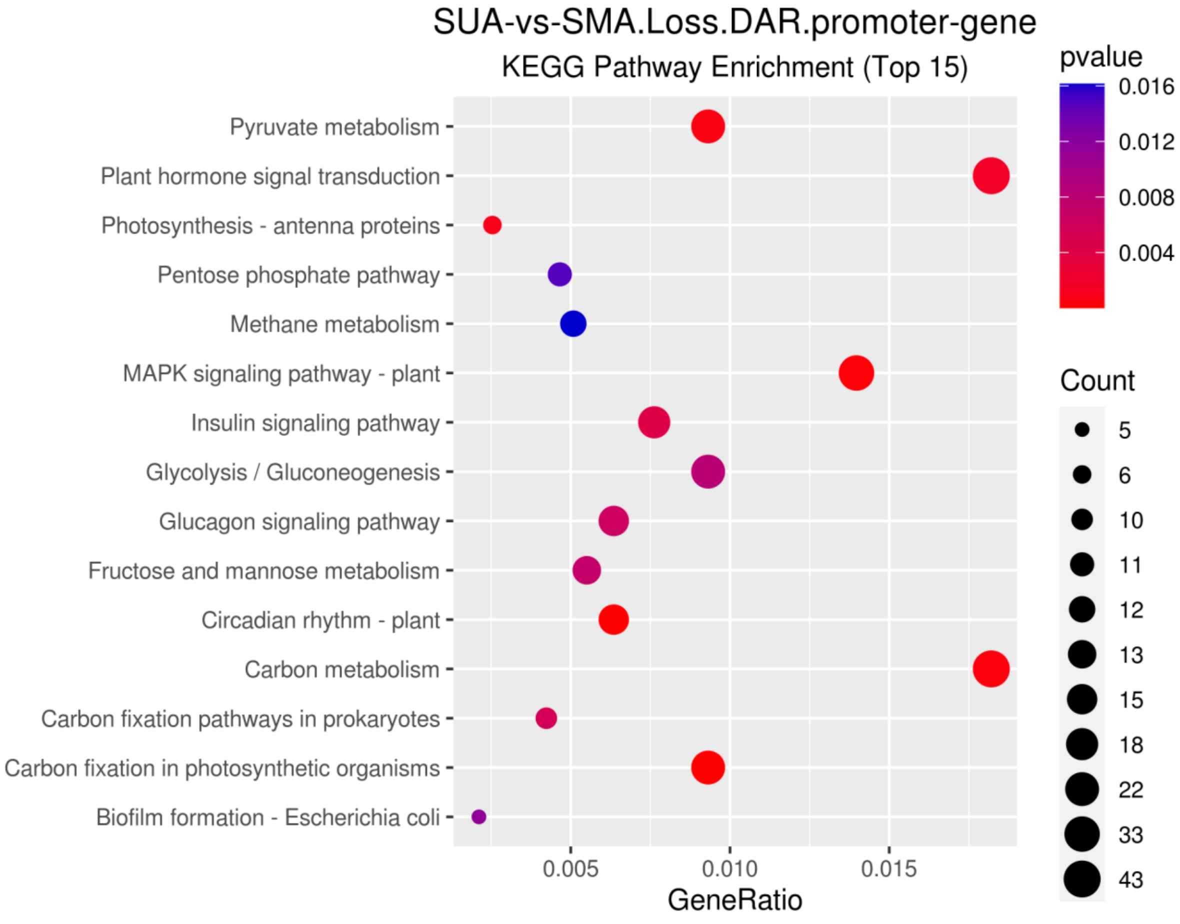


**Supplementary Figure 38: KEGG functional enrichment of the promoter gene of Loss DAR.** The horizontal axis is the GeneRatio, the proportion of the number of genes enriched to the pathway in the gene set, and the vertical axis is the Pathway. The color in the figure is the size of the p-value, and the circle is the size of the number of genes.

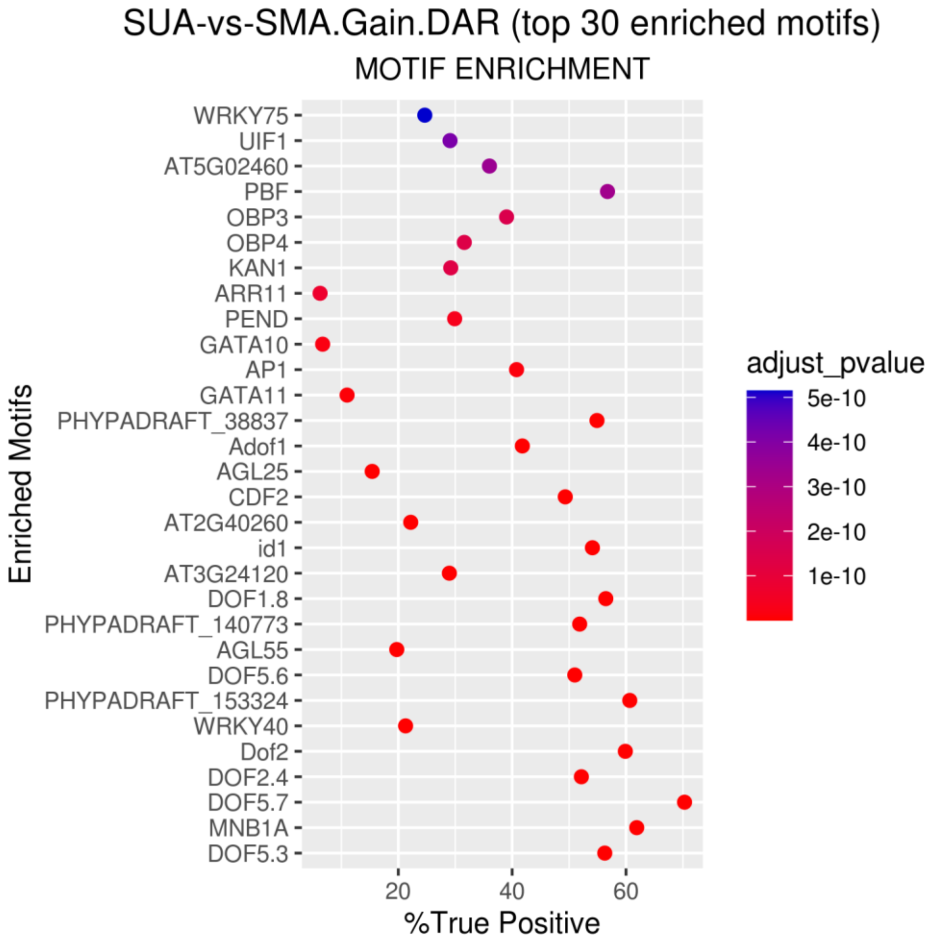


**Supplementary Figure 39: Motif enrichment results for Gain DAR.** The horizontal axis is the proportion of the number of DARs with the Motif in the DAR set, and the vertical axis is the Motif. The color in the figure is the size of the corrected p-value.

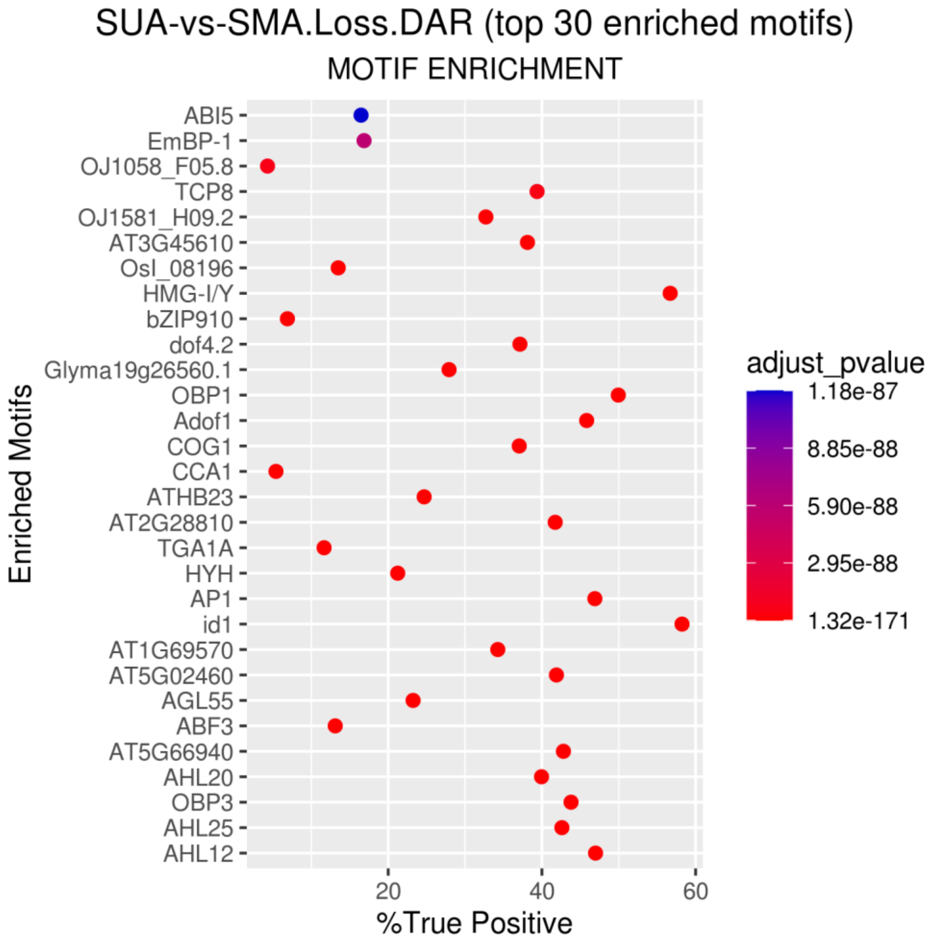


**Supplementary Figure 40: Motif enrichment results for Loss DAR.** The horizontal axis is the proportion of the number of DARs with the Motif in the DAR set, and the vertical axis is the Motif. The color in the figure is the size of the corrected p-value.

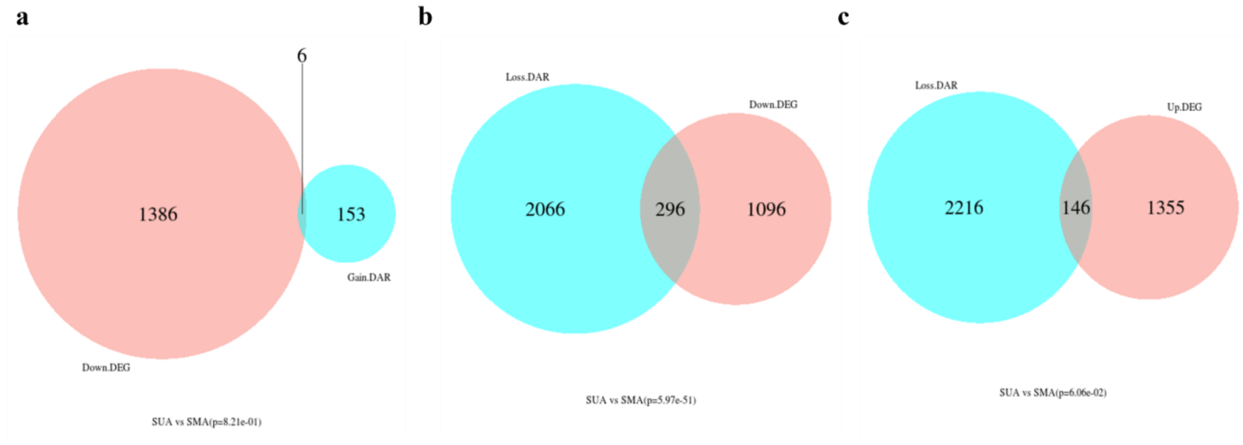

**Supplementary Figure 41: Comparison of DAR-associated genes and differentially expressed genes in *S. undatus* vs *S. megalanthus*. a)** Venn plot of gain DAR vs down DEGs **b)** Venn plot of loss DAR vs up-DEGs **c)** Venn plot of loss DAR vs down DEGs
